# Activation loop plasticity and active site coupling in the MAP kinase, ERK2

**DOI:** 10.1101/2023.04.15.537040

**Authors:** Laurel Pegram, Demian Riccardi, Natalie Ahn

## Abstract

Changes in the dynamics of the protein kinase, ERK2, have been shown to accompany its activation by dual phosphorylation. However, our knowledge about the conformational changes represented by these motions is incomplete. Previous NMR relaxation dispersion studies showed that active, dual-phosphorylated ERK2 undergoes global exchange between at least two energetically similar conformations. These findings, combined with measurements by hydrogen exchange mass spectrometry (HX-MS), suggested that the global conformational exchange involves motions of the activation loop (A-loop) that are coupled to regions surrounding the kinase active site. In order to better understand the contribution of dynamics to the activation of ERK2, we applied long conventional molecular dynamics (MD) simulations starting from crystal structures of active, phosphorylated (2P), and inactive, unphosphorylated (0P) ERK2. Individual trajectories were run for (5 to 25) *µ*s and totaled 727 *µ*s. The results showed that the A-loop is unexpectedly flexible in both 2P- and 0P-ERK2, and able to adopt multiple long-lived (>5 *µ*s) conformational states. Simulations starting from the X-ray structure of 2P-ERK2 (2ERK) revealed A-loop states corresponding to restrained dynamics within the N-lobe, including regions surrounding catalytic residues. One A-loop conformer forms lasting interactions with the C-terminal L16 segment and shows reduced RMSF and greater compaction in the active site. By contrast, simulations starting from the most common X-ray conformation of 0P-ERK2 (5UMO) reveal frequent excursions of A-loop residues away from a C-lobe docking site pocket and towards a new state that shows greater dynamics in the N-lobe and disorganization around the active site. Thus, the A-loop in ERK2 appears to switch between distinct conformational states that reflect allosteric coupling with the active site, likely occurring *via* the L16 segment. Analyses of crystal packing interactions across many structural datasets suggest that the A-loop observed in X-ray structures of ERK2 may be driven by lattice contacts and less representative of the solution structure. The novel conformational states identified by MD expand our understanding of ERK2 regulation, by linking the activated state of the kinase to reduced dynamics and greater compaction surrounding the catalytic site.

## Introduction

The MAP kinases, ERK1 and ERK2, are key effectors in the MAP kinase cascade, a signaling pathway downstream of RAS that is essential for cell proliferation, differentiation, motility, and survival [28, 58]. ERKs are activated by dual phosphorylation of specific threonine and tyrosine residues on the activation loop (A-loop), both catalyzed by upstream MAP kinase kinases 1 and 2 (MKK1/2 *aka* MEK1/2). MKK1/2 in turn are activated by members of the RAF family of protein kinases in all cells, and by c-MOS in germ cells. The prevalence of oncogenic mutations in RAS and RAF has motivated the successful development of inhibitors towards B/C-RAF and MKK1/2 for the treatment of melanomas and other cancers. Preclinical outcomes suggest the efficacy of ERK inhibitors towards cancers resistant to RAF or MKK inhibitors [17, 37]. Therefore, mechanisms involved in ERK activation are important to understand when targeting it for cancer therapeutics [54].

X-ray crystallographic studies of the phosphorylated (2P) and unphosphorylated (0P) states of ERK2 have provided a framework for understanding structural changes associated with kinase activation [8, 73] (**Fig. 1**). The largest conformational change occurs in the activation loop (A-loop), which contains the phosphorylation sites. Remodeling of the A-loop results in salt bridge interactions between pT183 and pY185 (rat ERK2 numbering throughout) and multiple Arg residues in the kinase N- and C-lobes (**Fig. 1A**). The reorientation of pY185 opens a proposed recognition site for proline-directed sequence motifs in ERK substrates [8], and a rearrangement of residues N-terminal to the phosphorylation sites (F181, L182) exposes a C-lobe binding site for a hydrophobic docking motif (“DEF”) found in ERK substrates and effectors[30].

**Figure 1.**
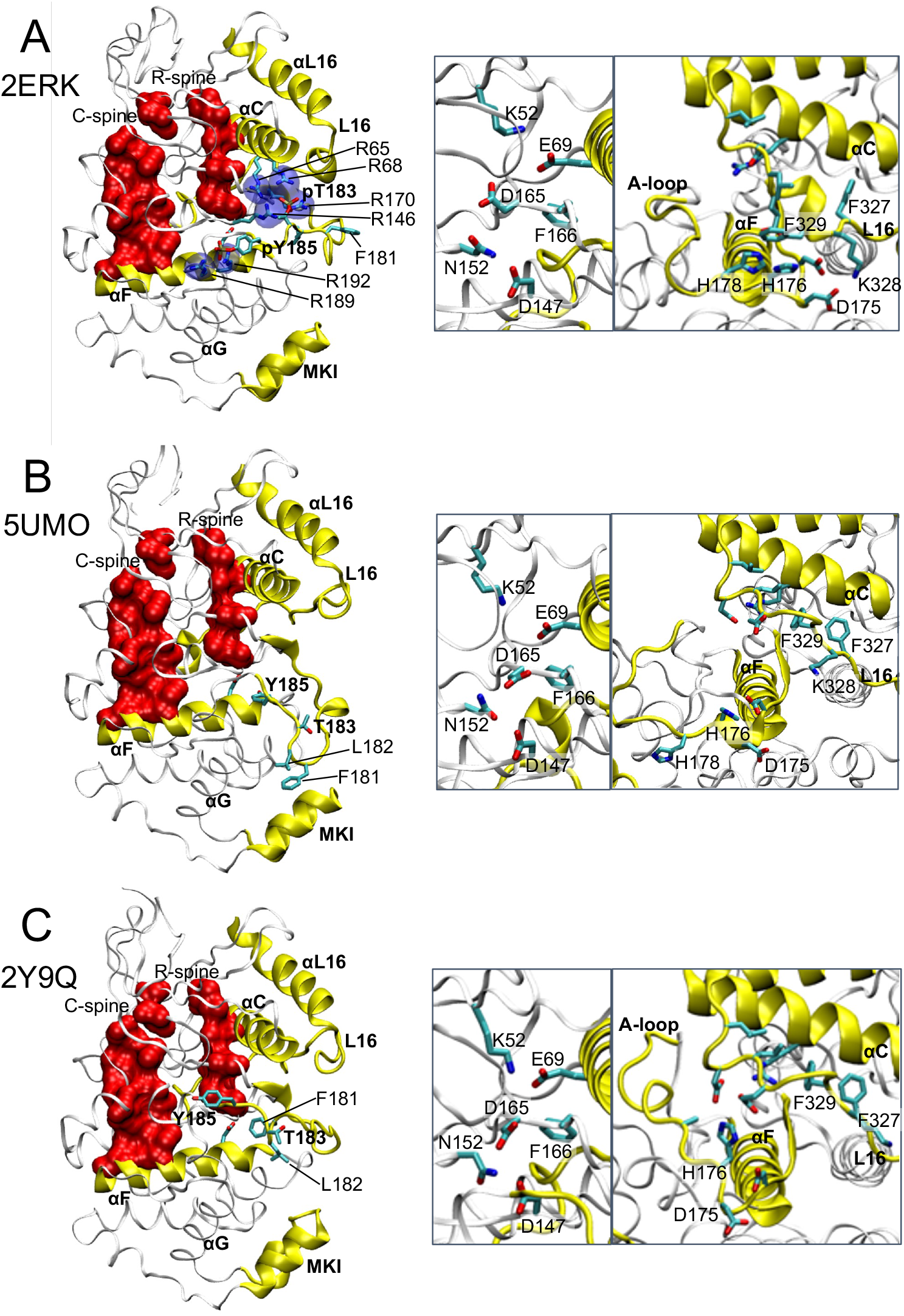
Crystal structures of 2P-ERK2 and 0P-ERK2. (**A**) The structure of the 2P-ERK2 apoenzyme, PDBID:2ERK, used as the starting state for all 2P-ERK2 simulations. (**B,C**) Structures of 0P-ERK2, as (**B**) apoenzyme, PDBID:5UMO, and (**C**) a peptide ligand complex, PDBID:2Y9Q, with the peptide removed, were used as starting states for 0P-ERK2 simulations. A third starting state for 0P-ERK2 used PDBID:2ERK after removing the phosphate groups from pT183 and pY185. Labelled residues show the pT183 and pY185 phosphorylation sites and salt-bridged Arg residues in 2ERK; T183, Y185, and A-loop residues F181 and L182 interact with the C-lobe in 5UMO. Close-up structures show (left panel) positions of active site residues that participate in catalysis, and (right panel) residue contacts between helices *α*C, *α*F, and the L16 segment.

Despite the large differences in the A-loop, structural differences between the active sites in the crystal structures of 0P-ERK2 and 2P-ERK2 are minimal. This contrasts with other protein kinases, where X-ray structures reveal significant conformational shifts that commonly accompany the switch from active to inactive states [64]. These include rotation of helix *α*C and consequent disruption of a critical Lys-Glu salt bridge (K52-E69 in ERK2) that coordinates phosphate oxygens in ATP; a “DFG flip” backbone rotation that buries a catalytic Asp residue (D165 in ERK2) needed for Mg^2+^ coordination; and disrupted alignments of regulatory-spine (R-spine) and catalytic-spine (C-spine) residues involved in nucleotide binding and phosphoryltransfer [12, 25, 34, 71]. Oddly, the positions of these active site residues are largely invariant between the crystal structures of the active 2P and inactive 0P forms of ERK2. Thus, ERK2 has been considered as a prototype to investigate regulatory mechanisms in kinases that do not display substantial conformational rearrangements at the active site.

Solution measurements have revealed changes in protein dynamics following ERK2 phosphorylation and activation. Studies using hydrogen-deuterium exchange mass spectrometry (HX-MS) showed that phosphorylation of ERK2 altered rates of deuterium uptake in localized regions where X-ray structures were invariant [19]. NMR Carr-Purcell-Meiboom-Gill (CPMG) relaxation dispersion measurements of [methyl-^13^C,^1^H]-Ile, Leu and Val residues in ERK2 revealed that activation by phosphorylation led to global exchange behavior within the N- and C-lobes and surrounding the active site. This exchange was modeled by an equilibrium between two energetically similar conformational states, named “L” and “R”, interconverting on a millisecond timescale [69]. Importantly, residues in the ERK2 A-loop were included in the global exchange, and mutations in the A-loop blocked formation of the R-state, demonstrating allosteric coupling from the A-loop to residues surrounding the active site [20]. Furthermore, different ATP-competitive inhibitors of ERK2 displayed conformational selection for the L and R states, shifting the L⇌R equilibrium in opposite directions [42, 52]. These inhibitors induced changes in HX protection within the P+1 segment adjoining the A-loop, confirming coupling from the active site to the A-loop [42]. Together, the results revealed an allosteric mechanism in 2P-ERK2, where the A-loop is not found in one settled state but instead interconverts between discrete states that are in turn coupled to motions in the active site. The nature of these states and how they contribute to ERK2 activation are unknown.

Inspired by this solution-phase evidence for the importance of dynamics in ERK2 activation, we applied long conventional molecular dynamics simulations to characterize potential motions in ERK2 and their structural framework. The results show multiple long-lived conformations of the A-loop that have not previously been observed by crystallography. Notably, settled conformations of the phosphorylated A-loop variably formed stable interactions with the N-lobe L16 segment or the C-lobe MAPK insert (MKI), or disrupted canonical salt-bridges between pY185 and the C-lobe to allow novel contacts with helix *α*C. Simulations of 0P-ERK2 showed new settled states of the A-loop that exposed the Y185 phosphorylation site to solvent. Difference contact network analyses and RMSF calculations revealed correlations between A-loop movements and dynamics of active site residues. Settled states of 2P-ERK2 were correlated with reduced dynamics and compactness of the N-lobe and active site, while 0P-ERK2 states suggested greater N-lobe motions and higher levels of active site disorganization. The results reveal unexpected flexibility in the A-loop, and a role for multiple conformational states in regulating active site dynamics in a phosphorylation-dependent manner.

## Results

Crystal structures of 2P-ERK2 and 0P-ERK2 were used as starting models to explore the dynamics of the A-loop by MD. The structure of the 2P-ERK2 apoenzyme (PDBID: 2ERK) shows extensive salt-bridge contacts between pT183 and pY185 in the A-loop and Arg residues in the N- and C-lobes (pT183 with R65, R68, R146, R170 and pY185 with R189, R192) (**Fig. 1A**). In addition, side chain interactions are formed between H176 in the A-loop and F329 in L16, a C-terminal segment which is uniquely found in MAP kinases [38]. Following energy minimization, this structure was used as the starting model for active, phosphorylated ERK2 (2erk 2P). The structure of the 0P-ERK2 apoenzyme (PDBID: 5UMO) shows the A-loop in a folded conformation with T183 exposed and Y185 buried (**Fig. 1B**). The A-loop residues F181 and L182 contact the C-lobe in a pocket formed between helix *α*G and the MAP kinase insert (MKI), the latter is also unique to MAP kinases and has been shown to recognize a sequence motif (“docking domain for ERK2, FXF (DEF)”) found in substrates and effectors. A second X-ray structure of 0P-ERK2 complexed with a kinase interaction motif (KIM) peptide (PDBID: 2Y9Q) shows an alternative A-loop conformation, with F181 and L182 removed from the *α*G-MKI DEF pocket (**Fig. 1C**). These were used as starting models for inactive 0P-ERK2 (5umo 0P, 2y9q 0P). A third starting model for 0P-ERK2 was constructed from 2ERK by removing the phosphates on pT183 and pY185 (2erk 0P). Each of these structures show similarities in the positions of key active site residues that participate in catalytic turnover (**Fig. 1A-C**). These include the K52-E69 salt bridge, which enables hydrogen bonding to P*α* and P*β* oxygens in ATP; D165 and N152, which coordinate Mg^2+^ complexed with ATP; and D147, which is the general base for phosphoryltransfer.

In order to examine the ability of the A-loop to sample new conformational states, initial experiments used long conventional simulations between 10 *µ*s and 25 *µ*s (“1° seeds”) performed at four temperatures (285 K, 300 K, 315 K, 330 K). These were followed by multiple short simulations (“2° seeds”), typically 5.7 *µ*s each performed at 300 K, and started from different frames in 1° seed trajectories. In this way, the simulations combined complementary strategies of long trajectories at varying temperatures, with short runs from multiple starting states at a single temperature. Nomenclature and details of runs are summarized in **Table 1**. The accumulated sampling times summed over all temperatures and seeds reached 369.5 *µ*s for 2P and 357.7 *µ*s for 0P starting models, totaling 727.2 *µ*s over all seeds.

**Table 1.** Summary of nomenclature and trajectory sampling. Each time listed represents that of a single trajectory. For example, 14 additional 2° seeds of 2erk 2P.pY-R65 were each 5.70 µs at 300 K.

| PDBID | State | Starting model | 1° seeds |  |  |  | 2° seeds (300 K) |  |
| --- | --- | --- | --- | --- | --- | --- | --- | --- |
| | | | T (K) | Seed | MD state | Time (μs) | No. Seeds | Time (μs) |
| 2ERK | 2P | 2erk_2p | 285 | 1 | 2erk_2P.pY-R65 | 23.33 | 14 | 5.70 |
|  |  |  | 300 | 1 | 2erk_2P.L16 | 26.98 | - | - |
|  |  |  | 300 | 2 | 2erk_2P.solv | 21.60 | 16 | 5.70 |
|  |  |  | 315 | 1 | 2erk_2P.solv | 21.58 | - | - |
|  |  |  | 315 | 2 | 2erk_2P.solv | 19.50 | - | - |
|  |  |  | 330 | 1 | 2erk_2P.MKI | 17.10 | 12 | 5.70 |
| 5UMO | 0P | 5umo_0p | 285 | 1 | 5umo_0P.MKI | 23.69 | - | - |
|  |  |  | 300 | 1 | 5umo_0P.MKI | 20.50 | - | - |
|  |  |  | 300 | 2 | 5umo_0P.MKI | 20.10 | - | - |
|  |  |  | 315 | 1 | 5umo_0P.MKI | 21.80 | - | - |
|  |  |  | 315 | 2 | 5umo_0P.FL | 22.50 | 5 | 3 x 6.30<br>2 x 4.92 |
|  |  |  | 330 | 1 | 5umo_0P.MKI | 14.40 | - | - |
| 2ERK | 0P | 2erk_0p | 285 | 1 | 2erk_0P.solv | 23.05 | - | - |
|  |  |  | 300 | 1 | 2erk_0P.solv | 20.99 | - | - |
|  |  |  |  | 2 | 2erk_0P.solv | 20.72 | - | - |
| | | | 315 | 1 | 2erk_0P.Y-αC | 21.79 | 8 | 5.70 |
|  |  |  |  | 2 | 2erk_0P.solv | 19.80 | - | - |
|  |  |  | 330 | 1 | 2erk_0P.solv | 14.40 | - | - |
| 2Y9Q | 0P | 2y9q_0P | 300 | 1 | 2y9q_0P.F/Y | 9.90 | - | - |
|  |  |  |  | 2 | 2y9q_0P.F/Y | 9.90 | - | - |
|  |  |  |  | 3 | 2y9q_0P.F/Y | 9.90 | - | - |
|  |  |  |  | 4 | 2y9q_0P.F/Y | 9.90 | - | - |

### MD simulations of phosphorylated ERK2

#### Novel conformational states of the A-loop

Six 1° seeds were started from 2erk 2P and run for (18 to 25) *µ*s at varying temperatures (**Table 1**). In all runs, the A-loop (res. 170-186) deviated from the starting model within (3 to 5) *µ*s as measured by the RMSD of C*α* atoms. Typically, the RMSD of the A-loop segment increased to more than 5 Å after the simulations began and remained high, only rarely and transiently falling below 3 Å (**Fig. 2A-D**). By contrast, the rest of the kinase “core” backbone structure (res. 16-169 and 187-348) remained largely unchanged with RMSD mostly below 2.5 Å (**Fig. 2A-D**). The initial increases in RMSD for the A-loop were matched by increased fluctuations measured by RMSF (**Fig. 2E-H**). Although the trajectory at lowest temperature (285 K) (**Fig. 2C,G**) retained the A-loop of the starting state structure for the longest time, it was still followed by a rapid, sharp increase in RMSD and RMSF after 3 *µ*s (**Fig. 2G**). In three 1° seeds, residues in the A-loop formed new intramolecular interactions, dampening their fluctuations to reflect new settled conformations persisting with low RMSF for more than 5 *µ*s (regions with hatched lines in **Fig. 2A-C, E-G**). These were identified by scanning 1 *µ*s segments of each trajectory for regions with average A-loop RMSF less than 1.2 Å. This revealed three distinct “settled states” of the A-loop based on their low RMSF and long lifetimes (> 5 *µ*s), each deviating from the 2ERK crystal structure with RMSD > 5 Å.

**Figure 2.**
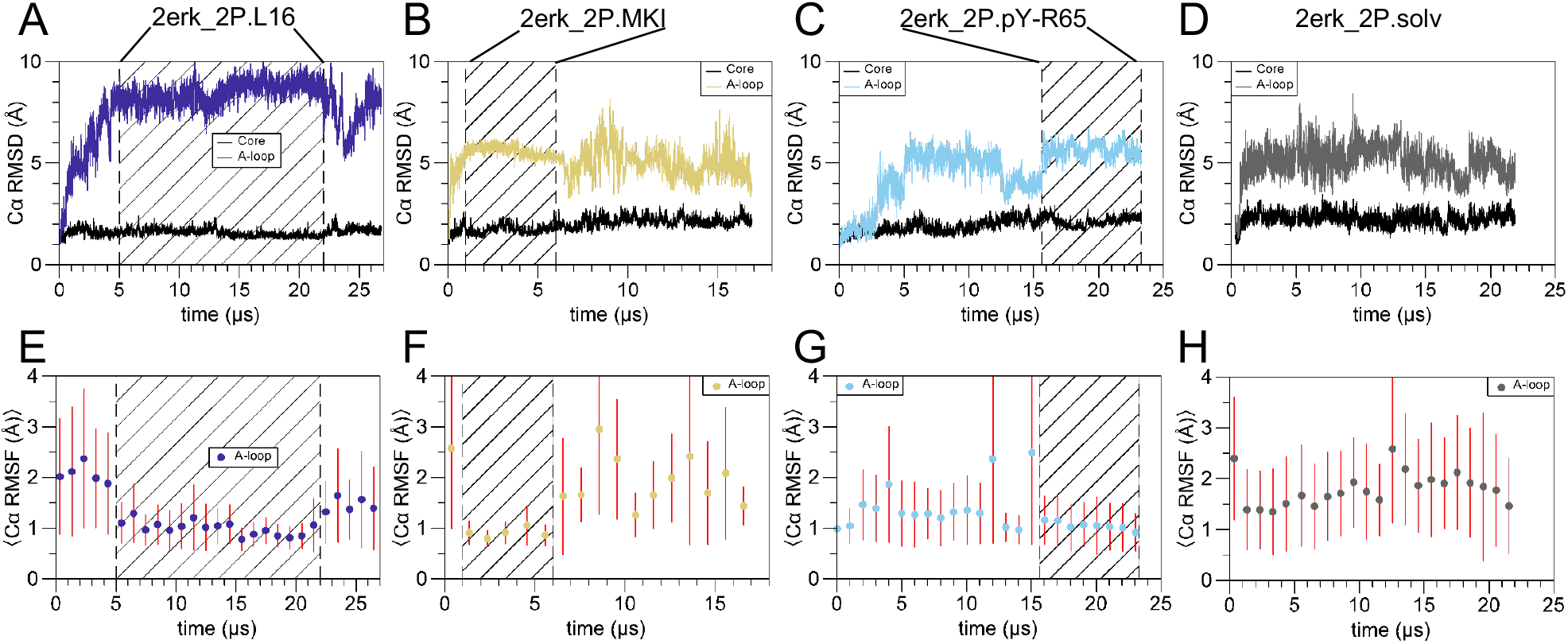
RMSD and RMSF plots identify settled states of the A-loop in 1° seed trajectories of 2P-ERK2. Trajectories corresponding to settled states (denoted by hatched lines) for **(A,E)** 2erk 2P.L16 (2erk 2P 1° seed 1 at 300 K), **(B,F)** 2erk 2P.MKI (2erk 2P 1° seed 1 at 330 K), and **(C,G)** 2erk 2P.pY-R65 (2erk 2P 1° seed 1 at 285 K), as well as **(D,H)** a representative trajectory where no settled state is reached, 2erk 2P.solv (2erk 2P 1° seed 1 at 315 K). **(A-D)** Plots of RMSD calculated for C*α* atoms in the A-loop (res. 170-186, colors) and kinase core (res. 16-169 and 187-348, black). **(E-H)** Plots of A-loop RMSF calculated for C*α* atoms showing averages and standard deviations for 1 *µ*s segments across each trajectory.

The three settled A-loop states are illustrated in **Fig. 3A-C** as overlays of configurations (colors) alongside the 2erk 2P starting model (black). **Fig. 3A** illustrates the “2erk 2P.L16” conformation, so named because of the A-loop movement towards the N-lobe where it forms contacts with residues in the C-terminal L16 segment. In this state, pT183 and pY185 engage in salt bridges with the same six Arg partners observed in 2ERK. The N-terminal region of the A-loop however, is remarkably different from 2ERK, with RMSD ∼8 Å between res. 170-186 (**Fig. 2A**). As a result, the side chain contacts between F329 in L16 and H176 in the A-loop of 2ERK (**Fig. 1A**) are replaced by contacts between F329 and P174 in 2erk 2P.L16 (**Fig. 3A**). In addition, F181 which is solvent-facing in 2ERK, remodels to contact L16 residues D335, L336, and P337 in 2erk 2P.L16. Thus, 2erk 2P.L16 introduces more side chain contacts between the A-loop and L16, which may enhance the stability of the new fold.

**Figure 3.**
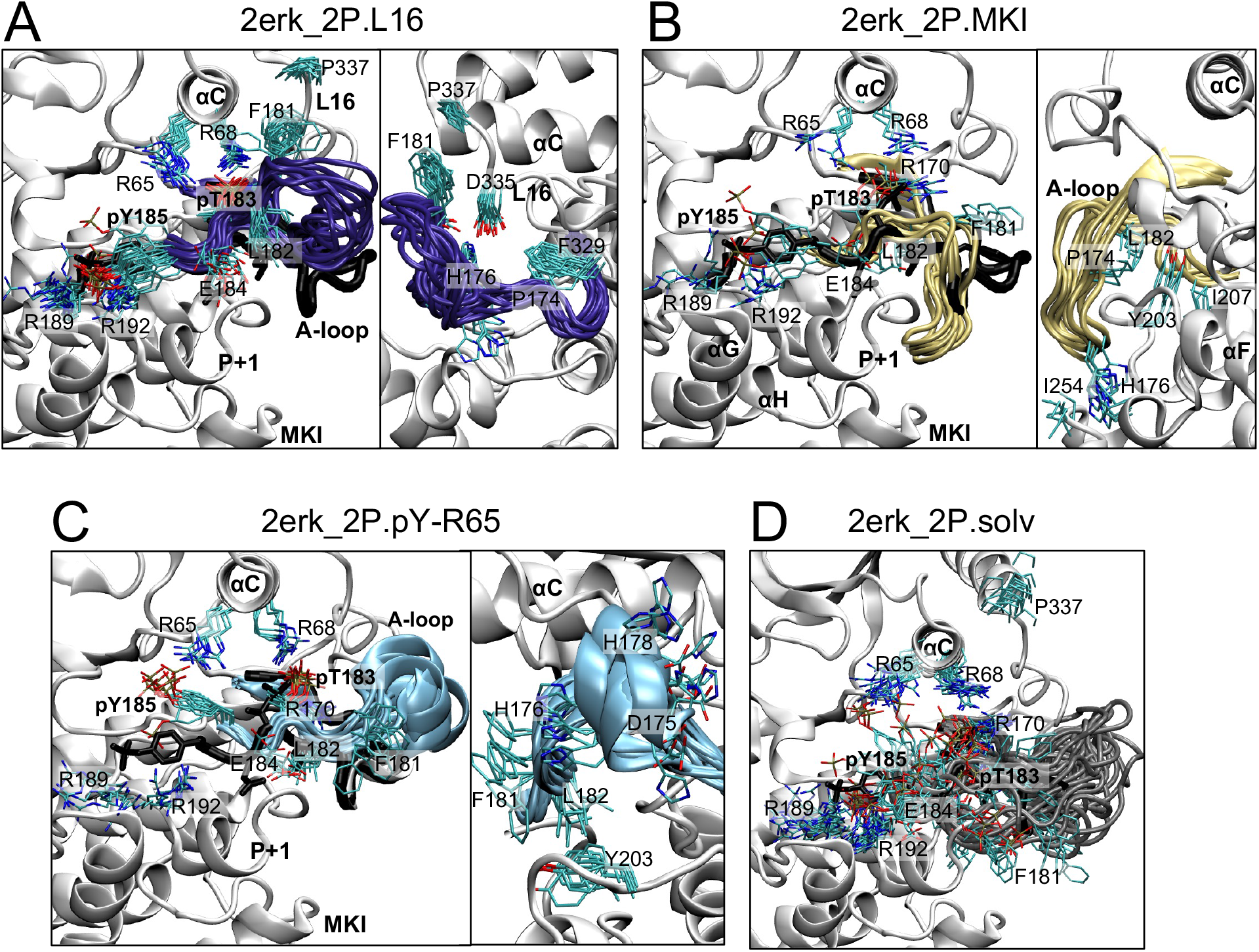
Settled states of the A-loop in 2P-ERK2. Overlays of A-loop conformers corresponding to settled states of **(A)** 2erk 2P.L16 (2erk 2P 1° seed 1, 300 K), **(B)** 2erk 2P.MKI (2erk 2P 1° seed 1, 330 K), **(C)** 2erk 2P.pY-R65 (2erk 2P 1° seed 1, 285 K), and **(D)** a representative trajectory with no settled A-loop state, 2erk 2P.solv (2erk 2P 1° seed 1, 315 K). Frames shown are separated by 1 *µ*s and derived from trajectories in Fig. 2. In each panel, the A-loop backbone and pT-E-pY side chains from 2ERK are shown in black and the kinase core is shown in white. Alternative orientations in panels **A-C** correspond to rotation by ∼90° about the vertical axis.

The “2erk 2P.MKI” conformation is displayed in **Fig. 3B**, so named because the A-loop extends towards the MAPK insert in the C-lobe. Such C-lobe interactions are reminiscent of the 5UMO crystal structure (**Fig. 1B**), except that the pT183-Arg and pY185-Arg salt bridges seen in 2ERK are retained. Like 5UMO, the A-loop in 2erk 2P.MKI forms many contacts with the P+1 segment, located between the A-loop and helix *α*F. The detailed contacts of A-loop residues, however, are quite different. Notably, the side chains of F181 and L182, which contact helix *α*G and MKI in 5UMO, are displaced in 2erk 2P.MKI. Here, F181 and L182 swing away from the C-lobe, such that F181 becomes solvent-exposed and engaged in cation-*π* interactions with R170, while L182 moves into a hydrophobic pocket with P174, Y203, and I207 in helix *α*F. Consequently, H176 contacts the C-lobe in a pocket formed between I254 in MKI and a loop between helices *α*H-*α*I. Thus, although the A-loop fold in 2erk 2P.MKI superficially resembles 5UMO, its specific residue contacts are remodeled to accommodate the phosphorylation of T183 and Y185. Relative to 2erk 2P.L16, greater variations occur in the pT183 and pY185 side chains of 2erk 2P.MKI, suggesting greater flexibility within the C-terminal region of the A-loop.

The 2erk 2P.pY-R65 conformation, displayed in **Fig. 3C**, breaks the salt-bridges between the pY185 phosphate and residues R189 and R192 in the P+1 loop of 2ERK, allowing pY185 to move towards helix *α*C and form a new salt-bridge with R65. Although transient proximity between pY185 and R65 occurred in many trajectories, their long-lived contacts with A-loop RMSF < 2 Å were observed in only one 1° seed (**Fig. 2C**). In this settled state, stable positioning of the phosphorylation motif (pT183-E184-pY185) was maintained, despite the large rotation of the pY185 sidechain; N-terminal A-loop residues formed additional internal hydrogen bonding interactions within a short helical segment.

The remaining trajectories displayed highly dynamic segments with solvent-exposed A-loop conformations (**Fig. 2D,H, Fig. 3D**). These were designated “2erk 2P.solv” conformers and, due to their high RMSF, were not considered a distinct settled state. In the 2erk 2P.solv ensemble, pT183 and pY185 both largely maintained crystallographic Arg salt-bridge interactions (along with occasional excursions of both residues) but with a highly mobile backbone around the phosphorylation motif compared to the more ordered settled states (**Fig. 3D**). Significantly, 2erk 2P.solv conformers often appeared before and after each of the three settled states above. This suggests that the 2erk 2P.solv state constitutes an intermediary region connecting A-loop settled states.

#### Mapping trajectory frames to different A-loop conformational states

Multiple frames (typically separated by 300 ns) were extracted from trajectory regions corresponding to three settled states (2erk 2P.L16, 2erk 2P.MKI, and 2erk 2P.pY-R65). These were used as the starting configurations for “2° seed” simulations, each carried out for 5.7 *µ*s at 300 K. The 2° seed trajectories were then characterized using the collective fraction of “native contacts” (*Q_A−loop_*) associated with a given settled state. In the present study, “native contacts” correspond to contacts between heavy atoms (within 4.5 Å) of the A-loop and kinase core for a given settled-state reference configuration (see Methods). The *Q_A−loop_* values for each settled state were calculated for every frame of all trajectories; all 1° and 2° trajectories run at 300 K were then separated into settled-state frame collections using *Q_A−loop_ >* 0.67 as the threshold. Any frame below the threshold for all settled states was assigned to 2erk 2P.solv (see Methods).

A schematic and example of the reference frame selection for 2erk 2P.MKI are shown in **Fig. 4A,B**. Secondary seed 43 was selected based on the uniformity of this settled state to the first frame across the trajectory. The reference structure is taken as the frame with the lowest RMSD from the average coordinates across the 2° trajectory (**Fig. 4C**). Similarly, reference frames were identified for 2erk 2P.L16 and 2erk 2P.pY-R65 (**Fig. 4D,E**). The reference frame structures illustrated the differences in contacts to the kinase core made by the A-loop, each differing between the three settled states and from the 2erk 2P starting state (**Fig. 4C-F**, red). Plots of the numbers of contacts made by each residue showed distinct patterns in the center of the A-loop (**Fig. S1**). In 2erk 2P.L16, F181 formed the largest number of contacts, reflecting interactions with Q60 (in helix *α*C), and D335, L336 and P337 (in L16). In 2erk 2P.MKI, H176 and D177 formed the highest number of contacts, interacting with residues C252 (MKI) and S200 (P+1), while in 2erk 2P.pY-R65, H178 contacted main chain atoms in D330-M331 (L16). Examples of *Q_A−loop_* values for trajectories are illustrated, where *Q_A−loop_ >* 0.67 for 2erk 2P.MKI revealed the presence of this settled state in a 1° seed between (2 to 7) *µ*s (**Fig. 4G**) or a 2° seed throughout the trajectory (**Fig. 4H**). Another 2° seed transitioned into the 2erk 2P.solv state within the first *µ*s (**Fig. 4I**). Thus, *Q_A−loop_* enabled the presence of the A-loop settled states to be measured over the course of each trajectory.

**Figure 4.**
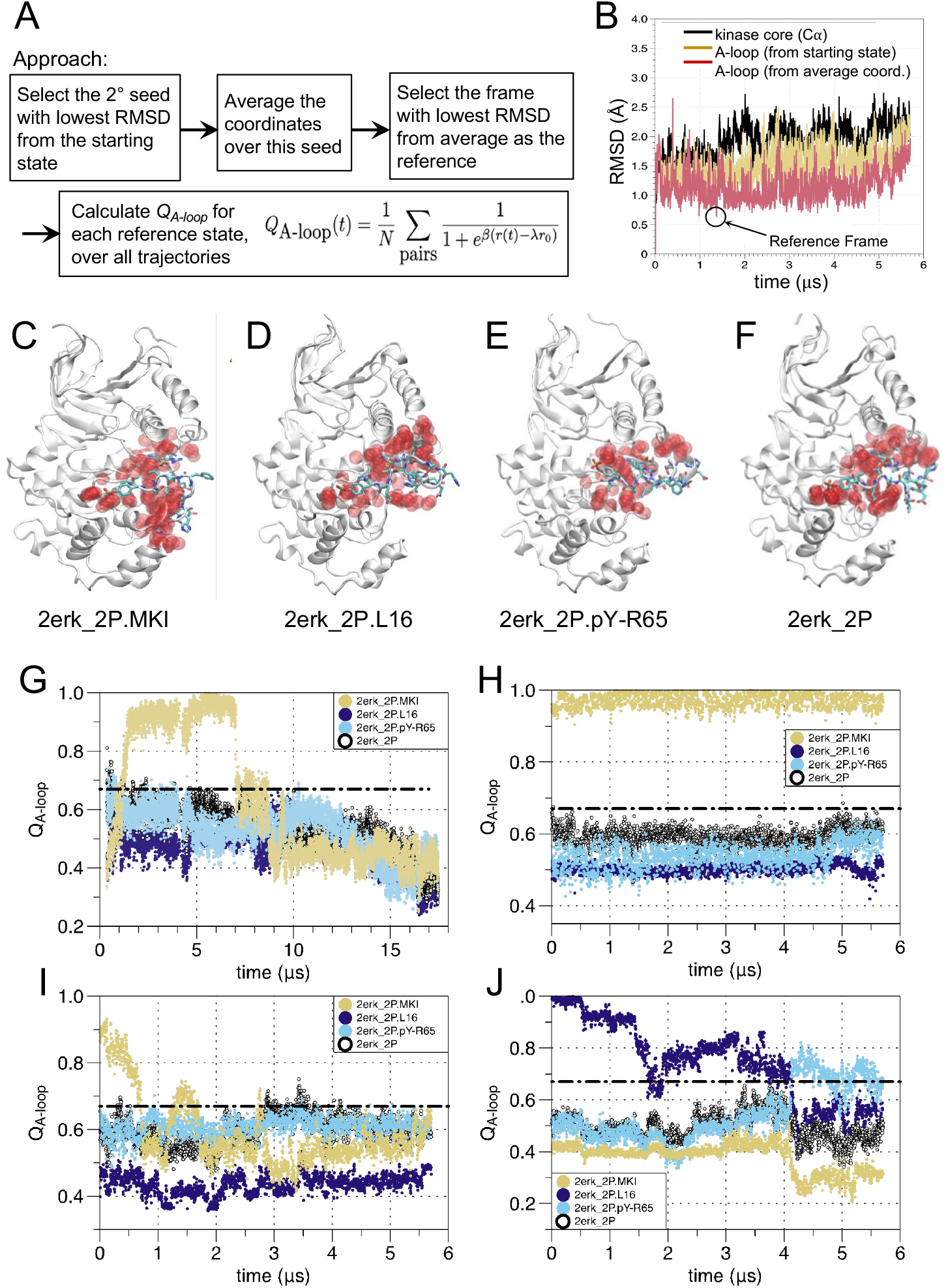
Selection of reference structures and *Q_A−loop_* plots for settled A-loop states in 2P-ERK2. **(A)** Schematic approach for determining reference structures and calculating *Q_A−loop_*. **(B)** An example shows how the reference structure for 2erk 2P.MKI was identified. Coordinates were averaged over a 2° trajectory, and the representative structure was determined from the frame that had A-loop heavy atom coordinates with the lowest RMSD from the averaged coordinates. **(C-E)** Reference structures for each settled state of 2P-ERK2, rendering A-loop residue sidechains (licorice) and their partner atoms from the kinase core that are within 4.5 Å (red spheres). **(F)** Corresponding representation of the 2erk 2P starting state. **(G-J)** Representative trajectories plotting *Q_A−loop_* calculated for each 2erk 2P state shown in panels **(C-F)**: **(G)** 2erk 2P 1° seed 1 at 330 K; **(H)** 2erk 2P 2° seed 43; **(I)** 2erk 2P 2° seed 42; **(J)** 2erk 2P 2° seed 39. *Q_A−loop_* and RMSD plots for all 2erk 2P 1° and 2° seed trajectories are provided in **Figs. S2** and **Figs. S3-S4**, respectively. The number of heavy atom contacts with A-loop residues for each reference state are shown in **Fig. S1**.

**Figs. S2**, **S3**, and **S4** show *Q_A−loop_* calculations and local A-loop RMSDs for the starting state and each reference state across all 1° and 2° seed trajectories. In all 1° seeds, *Q_A−loop_* for 2erk 2P quickly decayed while RMSD increased, indicating rapid divergence of the A-loop away from the starting state (**Fig. S2**, grey). This divergence was maintained in all 2° seed trajectories (**Figs. S3** and **S4**, grey). For the most part, agreement between *Q_A−loop_ >* 0.67 and RMSD < 2 Å occurred in trajectory regions corresponding to settled A-loop states. However in some trajectories, reduced RMSD suggested movements of the A-loop towards a settled state, even while *Q_A−loop_* remained below threshold (e.g. **Fig. S2** 2erk 2p.300.seed 2, gold, 8 *µ*s to 12 *µ*s). Other trajectories revealed elevated RMSD away from a settled state, even while *Q_A−loop_* remained high (e.g. **Fig. S2** 2erk 2p.285.seed 1, cyan, 13 *µ*s to 17 *µ*s). Thus, *Q_A−loop_* was a more reliable indicator for settled states of the A-loop than RMSD.

Analyses of the 1° and 2° seeds revealed evidence for switching between A-loop states. As noted above, 2erk 2P.MKI decayed in a 1° seed at 330 K and one 2° seed (**Fig. 4G,I**, gold). However, 2erk 2P.MKI extended over the full 5.7 *µ*s in 11 other 2° seeds (**Figs. S3**, **S4** 2erk 2p.300.seeds 3-10, 41, 43, 44, gold). This suggests that 2erk 2P.MKI forms transiently with lifetime > 5 *µ*s. Similarly, a trajectory leading to 2erk 2P.pY-R65 maintained this state for over 7 *µ*s in its 1° seed (**Fig. 2C**, cyan, 16 *µ*s to 23 *µ*s, **Fig. S2** 2erk 2p.285.seed 1, cyan), and throughout each of the 2° seeds started from this state (**Figs. S3**, **S4** 2erk 2p.300.seeds 11-24, cyan). By contrast, the 1° trajectory leading to 2erk 2P.L16 maintained a lifetime of more than 25 *µ*s (**Fig. S2** 2erk 2p.300.seed 1, purple), but varying lifetimes among 2° seeds, with *Q_A−loop_* falling below threshold in 10 of 16 cases (**Figs. S3**, **S4**, 2erk 2p.300.seeds 25-40, purple). In most instances, 2° seeds starting from 2erk 2P.L16 decayed to 2erk 2P.solv. In two trajectories, 2erk 2P.L16 decay was followed by increasing *Q_A−loop_* for 2erk 2P.pY-R65 (**Fig. 4J**, purple/cyan; **Figs. S3, S4**, 2erk 2p.300.seeds 27, 39, purple/cyan). Together, the results show that 2erk 2P.MKI, 2erk 2P.pY-R65, and 2erk 2P.L16 are all thermally accessible and populated in solution, with evidence for transitions to 2erk 2P.solv and between settled states.

#### Difference contact network analysis by A-loop state

Difference contact network analysis (dCNA) was applied to determine how variations between A-loop conformations led to changes in residue-residue contacts in other regions of the kinase. A-loop conformation ensembles were accumulated from 1° and 2° seed trajectories run at 300 K using a threshold *Q_A−loop_ >* 0.67 for each state (2erk 2P.L16, 2erk 2P.MKI or 2erk 2P.pY-R65). Frames with *Q_A−loop_* falling below the threshold for all three states were assigned to 2erk 2P.solv. Contact probability matrices for each ensemble were calculated as the fraction of the total frames containing contacts between residues, defined as any two heavy atoms within 4.5 Å. These symmetric, sparse matrices (where each dimension is the number of residues) were used to calculate a matrix of differences in contact probability for each residue pair between any two ensembles, as described by Hamelberg [13, 68, 72]. The contact probability matrix ranges from 0.0 to 1.0, and, thus, the difference in contact probability between two states ranges from -1.0 (probability decreases, contact is broken), through 0.0 (no difference between two states, e.g. diagonal elements), to 1.0 (probability increases, contact is formed). **Fig. 5A-C** illustrates increases (blue) and decreases (red) in contact probability between settled state ensembles (matrix elements for selected residues summarized in **Table S1**); the thickness of each bar indicates the magnitude of the probability change, with large magnitude differences characteristic of significant conformational changes, and small magnitudes expected for more subtle shifts around a single native state.

**Figure 5.**
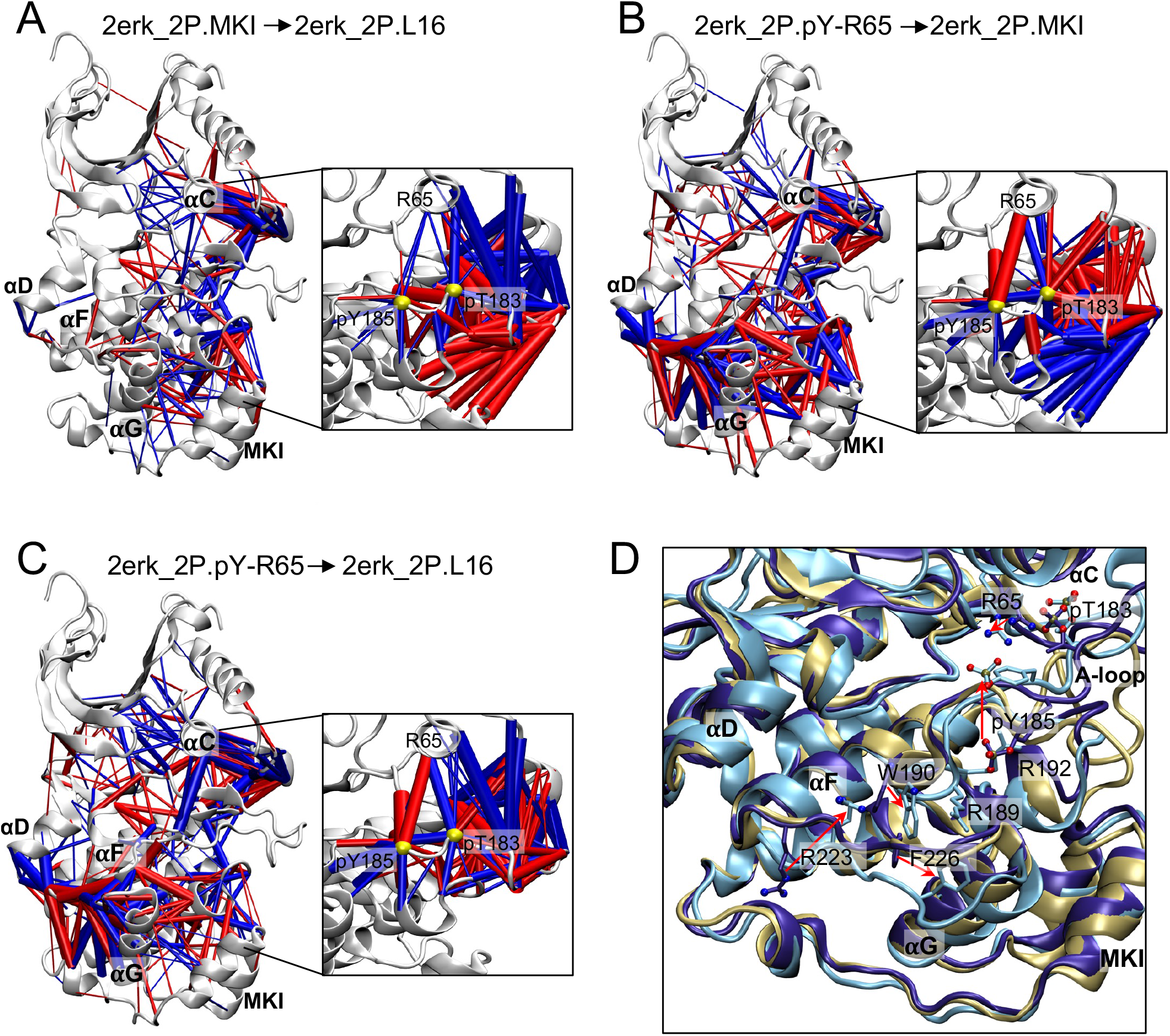
Difference contact network analyses (dCNA) for settled A-loop states in 2P-ERK2. Contact probabilities are accumulated for residue-residue pairs when their heavy atoms are within 4.5 Å from each other. All contact probability differences with absolute value *≥* 0.1 are shown, rendered as bars connecting C*α* atoms with radii scaled by the magnitude of the probability differences. Contact probabilities that increase or decrease between indicated states are shown in red or blue, respectively, going from **(A)** 2erk 2P.MKI to 2erk 2P.L16, **(B)** 2erk 2P.pY-R65 to 2erk 2P.MKI, and **(C)** 2erk 2P.pY-R65 to 2erk 2P.L16. For clarity, the differences in contacts made with residues in the A-loop are shown in insets, separately from contacts made only between residues in the kinase core. Yellow spheres in inset figures show C*α* atoms for pT183 and pY185. **(D)** Overlay of all three reference structures, showing the displacement of the backbone between 2erk 2P.L16 (purple), 2erk 2P.MKI (gold), and 2erk 2P.pY-R65 (cyan). Side chains highlight selected residues in 2erk 2P.L16 and 2erk 2P.pY-R65. Red arrows highlight major conformational movements of R223 and F226 in the loop between helices *α*F and *α*G and pY185 in the A-loop, and W190 in the P+1 segment.

Contacts to A-loop residues appeared with thick blue and red bars (**Fig. 5A-C**, insets), as expected from the large systematic structural variations between states (**Fig. 3A-C**). Residues surrounding the pT and pY phosphorylation sites (shown as yellow spheres in **Fig. 5A-C**, insets) formed a hub, reporting pronounced changes in contacts to kinase core residues. For example, bars between the A-loop and helix *α*C reflected the closer proximity between pY185 and R65 in 2erk 2P.pY-R65 compared to 2erk 2P.L16 or 2erk 2P.MKI, while bars between pT183 and R65 reflected their shifts away from each other in 2erk 2P.pY-R65 (**Fig. 5B,C**, insets). Likewise, bars connecting residues in the N-terminal A-loop shifted closer to the C-lobe in 2erk 2P.MKI compared to 2erk 2P.L16 or 2erk 2P.pY-R65 (**Fig. 5A,B**, insets). These reflected the closer interactions between the A-loop and the P+1, MKI, and *α*H-*α*I segments that are unique to 2erk 2P.MKI (**Fig. 3B**).

Changes in contact probabilities were also apparent between pairs of residues exclusively located outside of the A-loop. Two regions showed systematic differences between states. First, large changes between C-lobe residues were seen upon comparing 2erk 2P.pY-R65 to either 2erk 2P.L16 or 2erk 2P.MKI (**Fig. 5B,C**). These highlighted large shifts in residues in the loop between helices *α*F and *α*G (N222-F226) relative to those in helix *α*D, helix *α*G and MKI. Inspection of the reference structures revealed an obvious conformational change in the C-lobe of 2erk 2P.pY-R65 (**Fig. 5D**, cyan), where helix *α*G and the *α*F-*α*G loop moved towards MKI and away from the hinge and *α*D-*α*E loop compared to 2erk 2P.L16 and 2erk 2P.MKI (**Fig. 5D**, purple and gold). Trajectory overlays showed greater variability in main-chain and side-chain conformers in 2erk 2P.pY-R65, which, notably, drove R223 in the *α*F-*α*G loop from a solvent-exposed environment to one that was partially buried next to W190 in the P+1 loop (**Fig. 5D**, **Fig. S5**). These conformational movements in the C-lobe can be ascribed to the disruption of salt bridges between pY185 and the P+1 loop residues R189 and R192, which reoriented R189 towards helix *α*G, thus moving *α*G and the *α*F-*α*G loop towards MKI.

A second region with significant contact probability differences between A-loop settled state ensembles occurred in the N-lobe and active site. In this region, overlays between states showed only minor differences in backbone or side chain positioning (**Fig. 6A**). Nevertheless, a cluster of blue bars within the N-lobe revealed increased contacts in 2erk 2P.L16 compared to 2erk 2P.pY-R65 or 2erk 2P.MKI (**Fig. 5A,C**). These blue bars connected residues within conserved regions involved in ATP binding and phosphoryltransfer, including the Gly loop (G32, A33), *β*3 (K52), *α*C (Y62, E69) and the *β*9/DFG motif (D165, F166, G167). Examples illustrated in **Fig. 6B** show probability densities exhibiting shifts in these residues between different A-loop ensembles. Thus, K52-E69 which coordinates ATP, toggles between 2.9 Å and 4.5 Å with 2erk 2P.L16 mostly centered around an intact salt bridge, and 2erk 2P.pY-R65 shifted towards a disrupted salt bridge (**Fig. 6B**). Y34-G167, which reports contacts between the Gly loop and DFG, showed a dominant population around 4.2 Å in 2erk 2P.L16 which shifted towards 6 Å in 2erk 2P.pY-R65 and 2erk 2P.MKI. Similarly, shifts to longer distance were seen in A33-Y62, which reports contacts between the Gly loop and helix *α*C. Together, these findings suggest longer periods of Gly loop closure and closer proximity between K52 and D165 in the 2erk 2P.L16 ensemble. At the same time, red bars indicated greater separation between the DFG and HRD motifs in 2erk 2P.L16 (**Fig. 5A,C**). Here, 2erk 2P.L16 showed increased distances between residues D146 (catalytic base) and D165 or N152 (Mg^2+^-coordinating) compared to 2erk 2P.pY-R65 or 2erk 2P.MKI (**Fig. 6B**).

**Figure 6.**
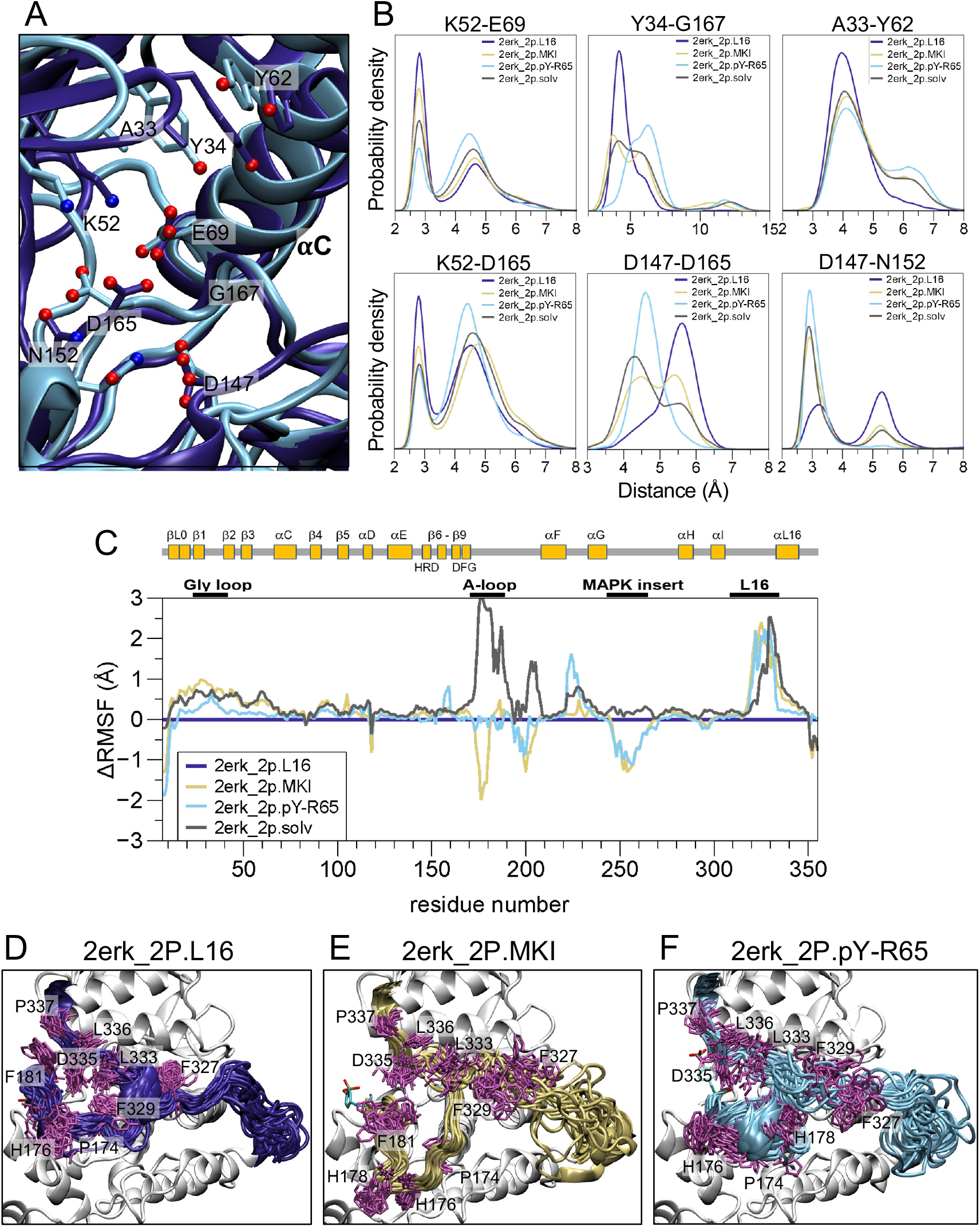
Differences in regional dynamics between settled A-loop states in 2P-ERK2. **(A)** Overlay of reference structures for 2erk 2P.L16 (purple) and 2erk 2P.pY-R65 (cyan), highlighting residues within the active site. **(B)** Probability densities for distances between residue pairs; each curve integrates to 1.0 for the region shown. Shifts in probability density reveal shortened distances involving catalytic residues in the K52-E69 salt bridge, the HRD motif (D147), and the DFG motif (D165, G167), and lengthened distances between HRD (D147, N152) and DFG (D165) motifs in 2erk 2P.L16 compared to other states. **(C)** Differences in RMSF between states, normalized to the 2erk 2P.L16 settled state (horizontal purple line). **(D-F)** Overlays of frames (every 250 ns) showing the regions of the A-loop and L16 segment for settled states **(D)** 2erk 2P.L16 (2erk 2P 2° seed 29), **(E)** 2erk 2P.MKI (2erk 2P 2° seed 43), and **(F)** 2erk 2P.pY-R65 (2erk 2P 2° seed 13).

In summary, dCNA revealed changes in contact between spatially clustered residues in different states of the A-loop. These reflected obvious conformational shifts that resulted from the movement of pY185 towards helix *α*C in 2erk 2P.pY-R65, disrupting its interactions with the P+1 loop and resulting in the movement of helix *α*G and the *α*F-*α*G loop towards MKI. At the same time, systematic changes in distance occurred in 2erk 2P.L16, which reflected greater compactness around conserved motifs involved in ATP binding and a shift towards greater opening around the catalytic base.

These results together with the absence of major conformational differences within the N-lobe suggested that changes in dynamics may contribute to residue compactness, such that 2erk 2P.L16 represents a more rigid, less dynamic mode of the active site than 2erk 2P.MKI or 2erk 2P.pY-R65. Inspection of root mean square fluctuations (RMSF) supported these changes in dynamics. **Fig. 6C** shows changes in fluctuations of C*α* atom coordinates compared to those of 2erk 2P.L16 (**Fig. 6C**, purple horizontal line centered at zero). As expected from **Fig. 3**, RMSF around the A-loop was largest for 2erk 2P.solv compared to other states (**Fig. 6C**, black), lowest for 2erk 2P.MKI (**Fig. 6C**, gold), and comparable between 2erk 2P.pY-R65 and 2erk 2P.L16. By contrast, RMSF values in 2erk 2P.MKI, 2erk 2P.pY-R65 and 2erk 2P.solv were systematically higher within the conserved regions of the N-lobe that form the ATP binding site and R-spine (Gly loop, helix *α*C, *β*3-*β*4-*β*5) (**Fig. 6C**, gold, cyan, black). Therefore, the shift towards compactness between N-lobe residues observed by dCNA is associated with more restrained dynamics in 2erk 2P.L16 compared to the other states.

The interactions between the A-loop and kinase core suggest an explanation for the unique ability of 2erk 2P.L16 to modulate distal N-lobe regions. Multiple contacts formed between the A-loop and L16 segment in 2erk 2P.L16 (e.g., P174-F329, F181-P337) were absent in 2erk 2P.MKI or 2erk 2P.pY-R65 (**Fig. 6D-F**). These resulted in a restrained L16 loop with reduced dynamics compared to the other states, as apparent from RMSF plots and trajectory overlays (**Fig. 6C-F**). As a result, the F327 side chain packs with hydrophobic residues in helix *α*C, which in turn connect to a network of hydrophobic residues in *β*3, *β*4, *β*5, and *β*9 that surround K52-E69 and DFG (**Fig. S6**). By contrast, larger movements of F327 in 2erk 2P.MKI and 2erk 2P.pY-R65 disrupted the interactions between L16 and the N-lobe. Thus, the reduced motions of L16 in 2erk 2P.L16 may explain the lower fluctuations and greater compaction in elements of the Gly loop, helix *α*C, and *β*3-*β*4-*β*5 compared to other states. This suggests that mutual interactions with L16 could allow the A-loop conformation to control the dynamics of essential residues in the active site.

### Simulations of unphosphorylated ERK2

#### Conformational ensembles of the unphosphorylated A-loop and *Q_A−loop_*analyses

Behaviors of 0P-ERK2 were examined using MD approaches similar to those described for 2P-ERK2. Three starting states were used, corresponding to the X-ray structure of the unphosphorylated apoenzyme (5UMO; starting state 5umo 0P, **Fig. 1B**), the structure of 2P-ERK2 (2ERK) after replacing pT183 and pY185 with unphosphorylated Thr and Tyr (2erk 0P, **Fig. 1A**), and a structure of 0P-ERK2 complexed with a kinase interaction motif (KIM) docking peptide (2Y9Q) after removing the peptide ligand (2y9q 0P, **Fig. 1C**).

Primary trajectories showed multiple A-loop conformations with low RMSF and lifetimes > 5 *µ*s (**Fig. 7**). Two distinct states were observed in 1° seeds started from 5umo 0P. One state, named “5umo 0P.MKI”, largely preserved the fold seen in the 5UMO crystal structure, with F181 anchored to the C-lobe and Y185 buried within the P+1 loop (**Fig. 7A**). In this state, L182 was also anchored to the C-lobe, exposing T183 to solvent, as in 5UMO. The RMSD of the A-loop remained within (3 to 4) Å from the starting state, while the kinase core backbone remained within 2 Å. After identifying a reference frame for 5umo 0P.MKI (**Fig. S7A**), *Q_A−loop_* and RMSD values were calculated across all 1° trajectories (**Fig. S8A,B**, green). The 5umo 0P.MKI conformation was observed in five of six 1° seed trajectories, displaying *Q_A−loop_ >* 0.67 and RMSD < 2 Å relative to its reference frame.

**Figure 7.**
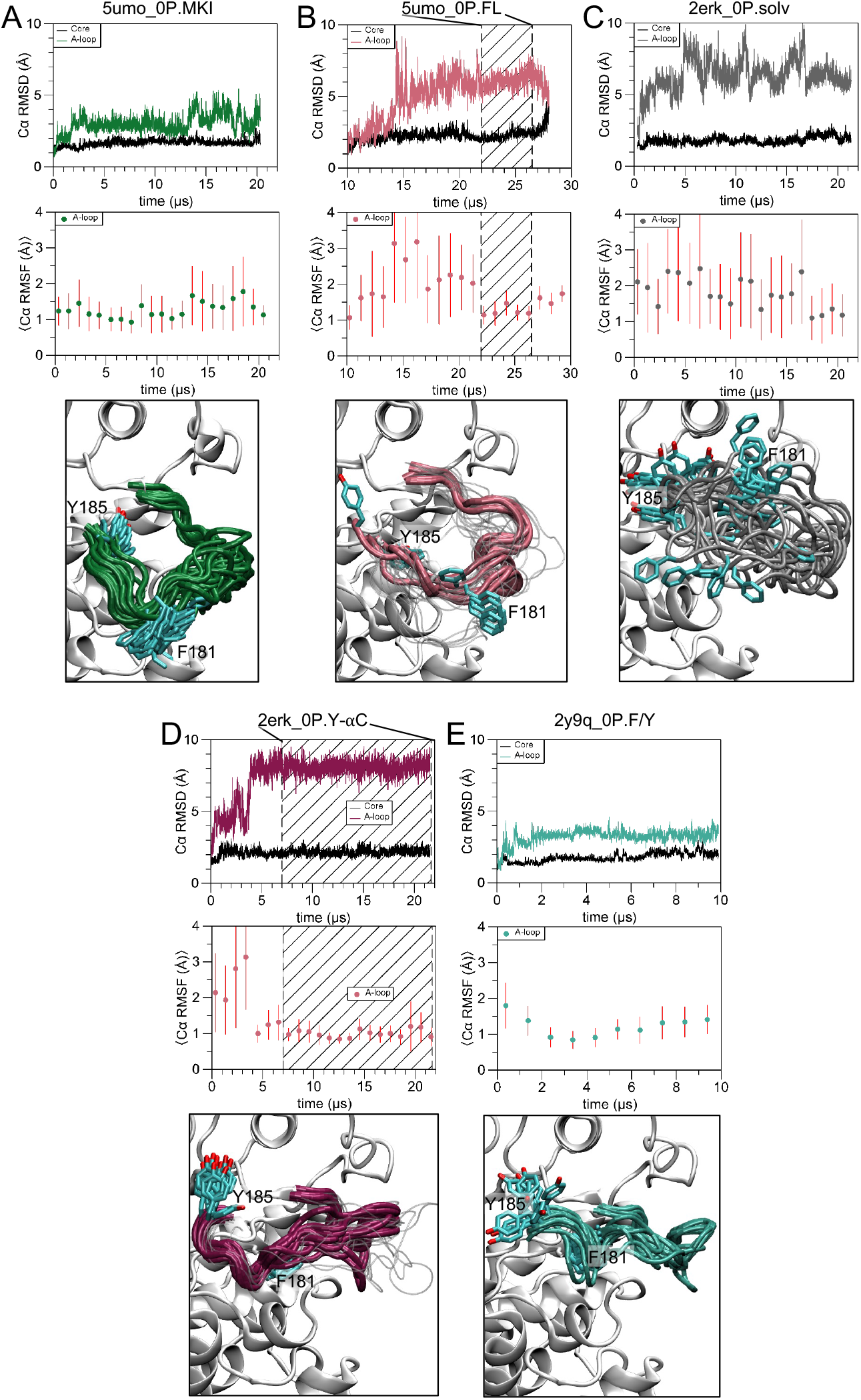
RMSD and RMSF plots identify settled states of the A-loop in 0P-ERK2. RMSD and RMSF plots for C*α* atoms, and overlay of frames corresponding to A-loop settled states, for **(A)** 5umo 0P.MKI (5umo 0P 1° seed 1 at 300 K), **(B)** 5umo 0P.FL (5umo 0P 1° seed 2 at 315 K), **(C)** a representative example of a seed that fails to form a settled state, 2erk 0P.solv (2erk 0P 1° seed 1 at 300), **(D)** 2erk 0P.Y-*α*C (2erk 0P 1° seed 1 at 315 K), and **(E)** 2y9q 0P.F/Y (2y9q 0P 1° seed 3 at 300 K). RMSD plots are shown for the A-loop (res. 170-186, colors) and kinase core (res. 16-169 and 187-348, black). RMSF plots show averages and standard deviations of the A-loop RMSF for 1 *µ*s segments across each trajectory. Overlays of A-loop conformers show frames separated by 1 *µ*s for each entire trajectory. For the two novel states (panels **B,D**), the settled states are shown by frames in color, and the frames outside the settled states are shown in gray. *Q_A−loop_* and RMSD plots are provided for all 0P-ERK2 1° and 2° seed trajectories in **Figs. S8** and **S9**, respectively.

A second state appeared in the remaining 1° seed started from 5umo 0P, reaching a new settled conformation with average A-loop RMSF < 1.5 Å between (22 to 27) *µ*s of the trajectory (**Fig. 7B**, **Fig. S8** 5umo 0p.315.seed 2). Here, interactions of F181 and L182 with the C-lobe were broken, exposing F181 and Y185 to solvent (**Fig. 7B**). This new state was therefore named “5umo 0P.FL” and was used to initiate multiple 2° seeds (**Fig. S9** 5umo 0p.300.seeds 3-7, salmon). After defining the reference states for 5umo 0P.MKI and 5umo 0P.FL (**Fig. S7A,B**), *Q_A−loop_* comparisons showed that once the 5umo 0P.FL A-loop conformation formed, it never returned to 5umo 0P in any 1° or 2° seed (**Fig. S8** 5umo 0p.315.seed 2, **Fig. S9** 5umo 0p.300.seeds 3-7, salmon).

In 1° seeds starting from 2erk 0P, most trajectories immediately deviated from the initial conformation, leading to a largely disordered A-loop with RMSD > 5 Å and average A-loop RMSF > 2 Å, which was named “2erk 0P.solv” (**Fig. 7C**). However, one 1° seed reached a settled state where the average A-loop RMSF decreased to < 1.2 Å after 7 *µ*s and persisted for the remainder of the trajectory (**Fig. 7D**, **Fig. S8** 2erk 0p.315.seed 1, maroon). In this new A-loop conformation, the main chain around the T-E-Y phosphorylation motif rotates, moving Y185 towards helix *α*C. Here Y185 remains solvent exposed, burying F181 into a pocket formed between HRD, P+1 and *α*F (**Fig. 7D**). This state was named “2erk 0P.Y-*α*C” (**Fig. S7C**). Multiple 2° seeds retained this conformer, which was clearly distinct from all others (**Fig. S9** 2erk 0p.300.seeds 3-10, maroon). Finally, 1° seeds starting from 2y9q 0P retained the A-loop conformation seen in the 2Y9Q X-ray structure (**Fig. 7E**), where F181 and L182 interactions with the C-lobe were disrupted, displacing Y185 to solvent. This state was named “2y9q 0P.F/Y” (**Fig. S7D**). RMSD and *Q_A−loop_* measurements showed that all 1° trajectories largely remained in the 2y9q 0P.F/Y state (**Fig. S8** 2y9q 0p.300.seeds 1-4, teal), therefore 2° seeds were not performed.

Together, these results revealed considerable conformational variation in the A-loop of 0P-ERK2. Importantly, each of the starting models accessed settled A-loop states with substantial residue exposure to solvent (5umo 0P.FL, 2erk 0P.Y-*α*C, 2y9q 0P.F/Y), as well as a disordered state (2erk 0P.solv). This was particularly significant for residue Y185, whose phosphorylation by MKK1/2 is kinetically favored over T183 [18]. The buried conformation of Y185 in the 5UMO crystal structure has obfuscated the order of phosphorylation. However, the experimentally observed order can be readily explained by these results from MD, which demonstrate multiple conformations of the A-loop that promote solvent exposure of Y185 and accessibility to MKK1/2.

#### Comparing A-loop states 0P- and 2P-ERK2 by difference contact network analysis

Upon examining different states of 0P-ERK2 by dCNA (**Fig. 8**), the largest red and blue bars reflected conformational differences in the A-loop and proximal core regions. Thus, 5umo 0P.MKI showed increased contacts of the A-loop with P+1, helix *α*G, and MKI in the C-lobe, compared to 2erk 0P.Y-*α*C and 2y9q 0P.F/Y (**Fig. 8A,B**). These reflected major conformational changes in F181 and L182 in the A-loop as well as Y203 in the P+1 segment between 5umo 0P.MKI, 2erk 0P.Y-*α*C and 2y9q 0P.F/Y, each of which formed or disrupted many heavy atom contacts between states.

**Figure 8.**
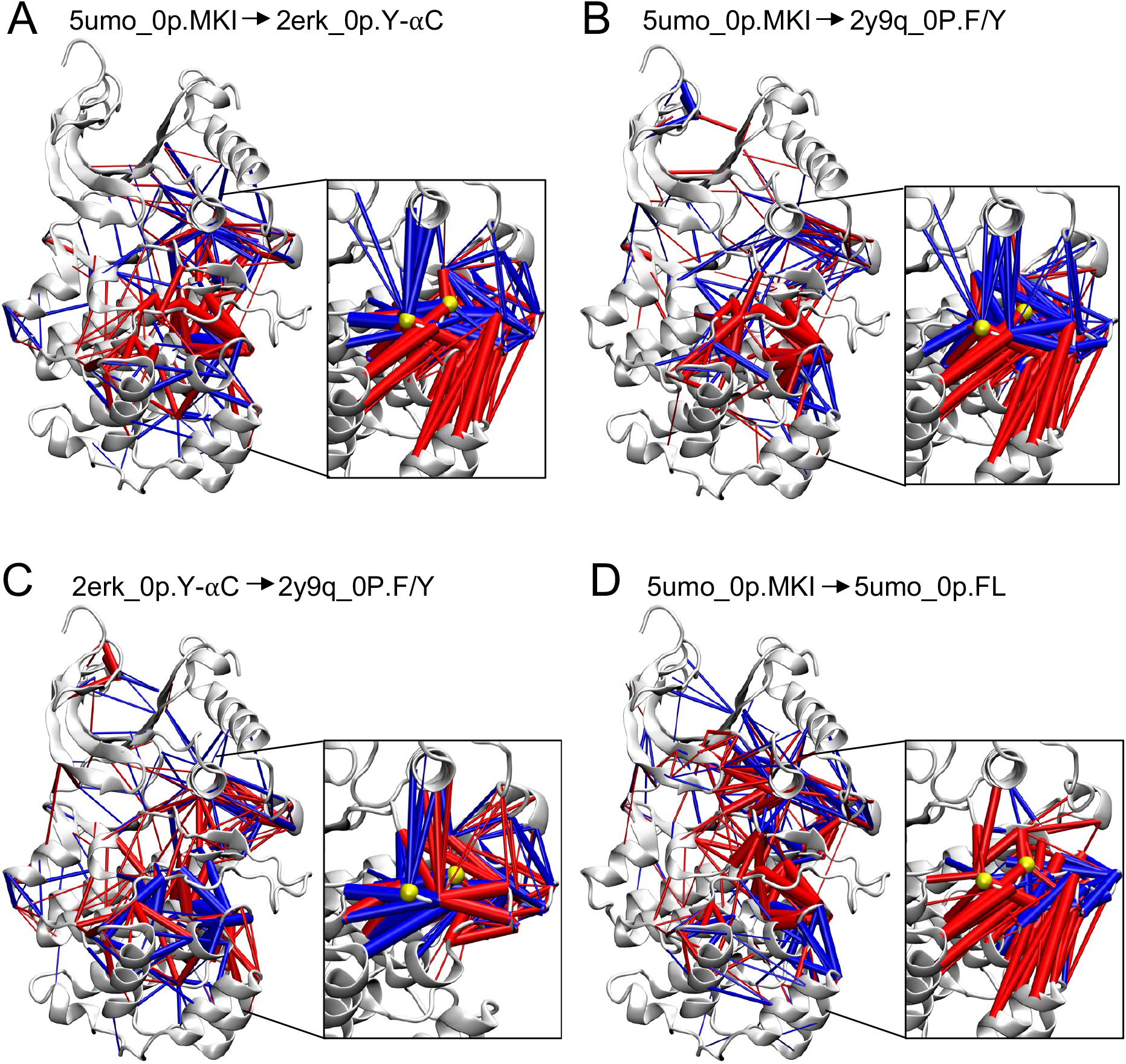
Difference contact network analyses (dCNA) for settled A-loop states in 0P-ERK2. Contact probabilities are calculated and displayed as described in Fig. 5. Shown are contact differences going from **(A)** 5umo 0P.MKI to 2erk 0P.Y-*α*C, **(B)** 5umo 0P.MKI to 2y9q 0P.F/Y, **(C)** 2erk 0P.Y-*α*C to 2y9q 0P.F/Y, and **(D)** 5umo 0P.MKI to 5umo 0P.FL. For clarity, the contact differences involving A-loop residues are shown in insets, separately from those made only between kinase core residues. Yellow spheres in inset figures show C*α* atoms for T183 and Y185.

Outside of the A-loop region, differences in residue contacts were less extensive between 5umo 0P.MKI, 2erk 0P.Y-*α*C, and 2y9q 0P.F/Y than between settled states of 2P-ERK2. For example, conformational changes in the C-lobe between the reference structures of 0P-ERK2 (**Fig. S7A-D**) were smaller than the movements seen in 2erk 2P.pY-R65 (**Fig 5D**, **Fig. S5**). Furthermore, the systematic shifts in contact probabilities that reflected compaction between N-lobe residues in 2erk 2P.L16 (**Fig. 5A,C**, **Fig 6B**) were largely absent between different 0P states (**Fig. 8**). Thus, for the most part, dCNA differences between 0P-ERK states were localized to the A-loop and regions in proximity. This contrasted with 2P-ERK2, where conformational variants of the A-loop were associated with perturbations at distal N-lobe regions.

#### A-loop residues F181 and L182 leave the DEF pocket in 5umo 0P.MKI to form a more dynamic state

The 5umo 0P.FL settled state was an exception to the observations above. Here, dCNA bars revealed substantial disruption of contacts between N-lobe residues in helix *α*C and the active site compared to 5umo 0P.MKI (**Fig. 8D**). Transitioning from 5umo 0P.MKI to 5umo 0P.FL consistently reduced contacts throughout the active site region and enhanced contacts to the MKI. This resulted in enhanced dynamics of 5umo 0P.FL, which we examined in greater detail. In order to more closely examine the transition from 5umo 0P.MKI to the 5umo 0P.FL settled state, the distances between C-lobe residue L232 (helix *α*G) and A-loop residues F181 and L182 were plotted for all 1° seeds of 5umo 0P (**Fig. 9**). Surprisingly, F181 transiently leaves the *α*G-MKI pocket (DEF motif binding site) in all trajectories, but stably exits the pocket only after 14 *µ*s in one seed (**Fig. 9A**, **Fig. 7B**, **Fig. S8** 5umo 0p.315.seed 2). By contrast, L182 remains in the *α*G-MKI pocket in all 1° trajectories except this seed, where it follows the F181 excursion at 15 *µ*s, fluctuating briefly before stably transitioning away from *α*G-MKI at 17 *µ*s (**Fig. 9B**). The RMSF of the A-loop increases sharply (**Fig. 7B**) and then decreases as the loop settles into its new state, with L182 buried near Y185. The results show that 5umo 0P.MKI accommodates transient excursions of F181 away from the C-lobe, until a point where L182-C-lobe contacts are disrupted, leading to cooperative movements of the A-loop to form the 5umo 0P.FL state. This provides novel insight into the importance of contacts formed by L182, in controlling the dynamics of A-loop movement.

**Figure 9.**
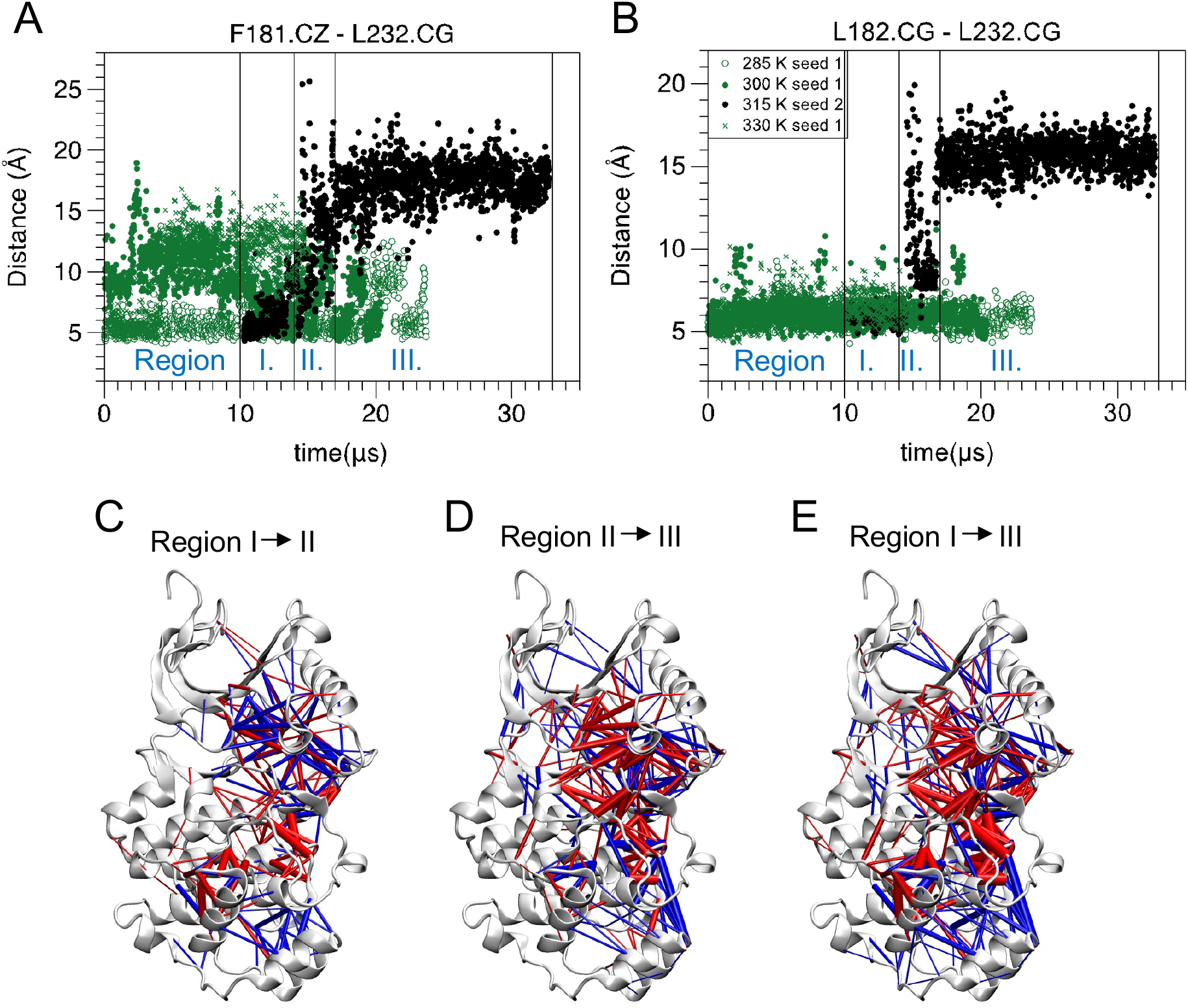
Dynamics of the A-loop and kinase core in the 5umo 0P.FL trajectory. Overlays of 1° seed trajectories, each starting from 5umo 0P. Distances between residues **(A)** F181 in the A-loop and L232 in the C-lobe, and **(B)** L182 in the A-loop and L232. Plots show frames every 12.5 ns. **(C-E)** Difference contact network analyses (dCNA) showing contact probability differences between three regions of the trajectory where 5umo 0P.FL appears (5umo 0P 1° seed 2 315 K), corresponding to time windows in Regions I (10 *µ*s to 14 *µ*s), II (14 *µ*s to 17 *µ*s) and III (17 *µ*s to 33 *µ*s).

This transition to the new A-loop state was explored further by examining the time dependence of dCNA, comparing three regions of the 1° seed trajectory (**Fig. 9C-E**). dCNA comparing the first 4 *µ*s (Region I) to the next 3 *µ*s (Region II) reveals the disruption of contacts between the A-loop and core regions involving the P+1 segment and loop preceding helix *α*F and helix *α*G; within the N-lobe, numerous dynamic changes were reflected by blue bars connecting helix *α*C to the active site and L16 (**Fig. 9C**). Completing the transition, dCNA comparing Region II to the remaining trajectory (Region III) reveals a large number of reduced contacts throughout the kinase core, as well as new contacts formed with the MKI (**Fig. 9D**). Overall, the dCNA between Region I to Region III reflects similar changes in contact described between 5umo 0P.MKI and 5umo 0P.FL (**Fig. 8D**).

Interestingly, contacts formed and broken appeared to fluctuate between the dCNA for Regions I *vs* II and for Regions II *vs* III (**Fig. 9C,D**), implying transient movements in localized regions of the kinase core. The fluctuations, in part, reflected large movements of helix *α*C, as measured by 80°-120° shifts in the pseudo-dihedral angle (*φ*) between helices *α*C and *α*E (**Fig. 10A,B**). Further separation of this trajectory into eight regions based on *φ* revealed large variations in contacts by dCNA (**Fig. S10A-G**). Time-dependent shifts in red and blue bars revealed fluctuations in conformation involving the DFG and HRD motifs and helix *α*C, which rapidly exchanged as *φ* increased or decreased. By contrast, conformational fluctuations were more restrained in trajectories of 2erk 2p, as illustrated for the 1° seed for 2erk 2P.MKI (**Fig. S11A-C**, **Fig. 4G**, **Fig. S2** 2erk 2p.330.seed 1). Here, *φ* ranged between 90°-105° and time-dependent dCNA reflected changes in contacts that were fewer and lower in magnitude (thinner red and blue bars) compared to the 5umo 0P.FL trajectory.

**Figure 10.**
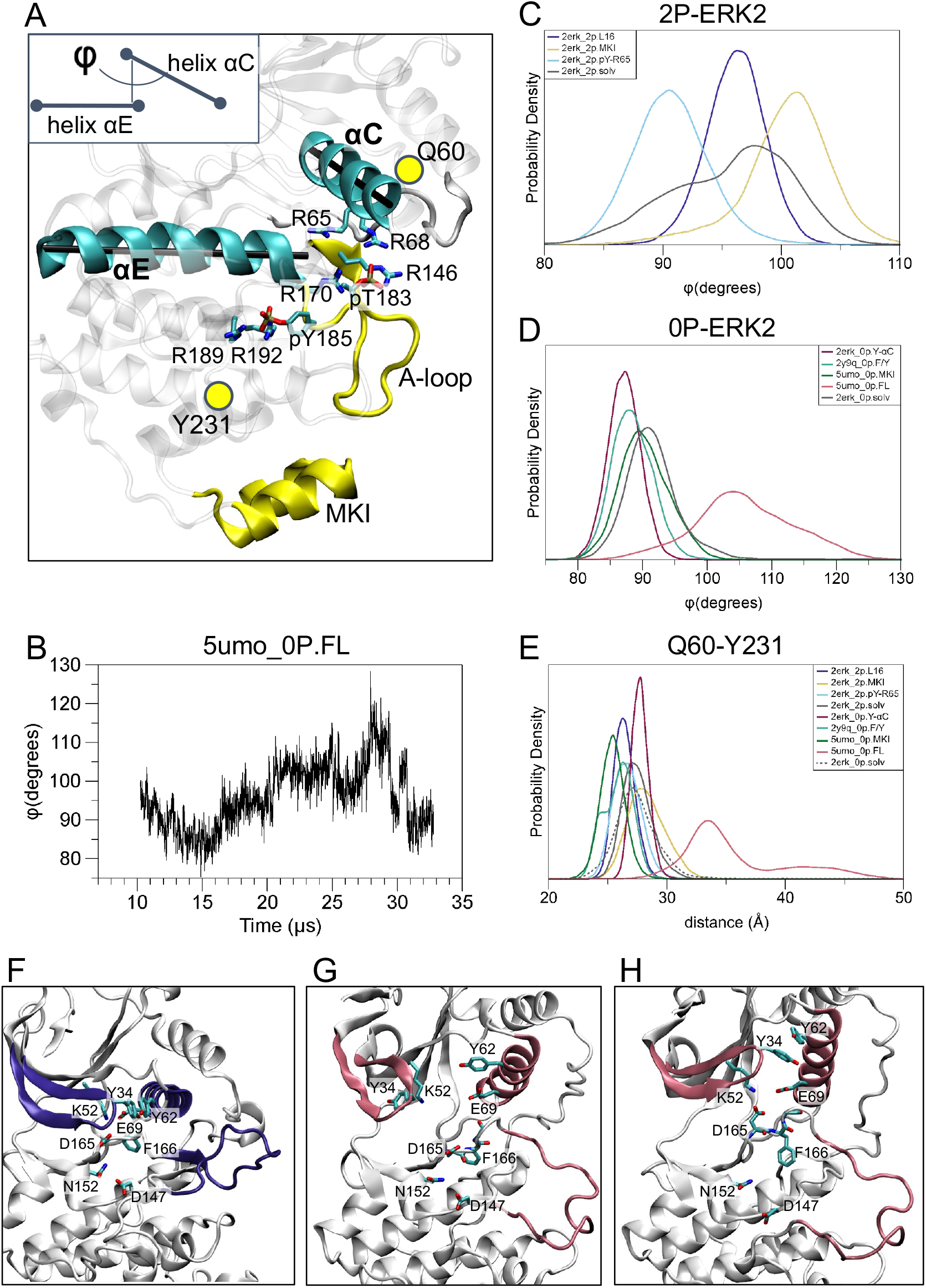
Variations in dihedral angles and domain separation in different A-loop states of 2P-ERK2 and 0P-ERK2. **(A)** The pseudo-dihedral angle between helices *α*C and *α*E (*φ*) is shown schematically for the 2erk 2P.MKI reference structure. **(B)** Fluctuations in *φ vs* time across the 1° trajectory containing 5umo 0P.FL (5umo 0p seed 2 at 315 K). **(C,D)** Probability densities for *φ* are shown for different A-loop states in **(C)** 2P-ERK2 and **(D)** 0P-ERK2. **(E)** Probability densities for the distance between Q60 and Y231 in different A-loop states of 2P-ERK2 and 0P-ERK2. **(F-H)** Representative frames illustrating variations in structure in 5umo 0P.FL.

#### Variations in the *α*C-*α*E pseudo-dihedral angle between 0P- and 2P-ERK2

The pseudo-dihedral angle between helices *α*C and *α*E has been used to report domain movements in ERK2 by measuring the degree of rotation between N- and C-lobes [2, 42]. Plots of *φ* for trajectories starting from 2erk 2P showed a probability distribution that was narrower for 2erk 2P.L16 relative to 2erk 2P.pY-R65 or 2erk 2P.MKI, and that broadened even further in 2erk 2P.solv (**Fig. 10C**). This is consistent with a greater degree of N-lobe compaction in 2erk 2P.L16 compared to other states. In addition, the magnitude of *φ* decreased in 2erk 2P.pY-R65 relative to 2erk 2P.L16, reflecting a shift of helix *α*C inwards, due to salt-bridge formation between pY185 and R65. The magnitude of *φ* increased in 2erk 2P.MKI relative to 2erk 2P.L16, reflecting movement of helix *α*C outwards, due to disruption of A-loop interactions with the N-lobe.

By contrast, the pseudo-dihedral angles were systematically lower in magnitude in nearly all trajectories of 0P-ERK2, with probability densities more comparable to 2erk 2P.pY-R65 than 2erk 2P.L16 or 2erk 2P.MKI (**Fig. 10D**). The distribution was shifted to higher values for 5umo 0P.MKI compared to 2erk 0P.Y-*α*C and 2y9q 0P.F/Y, a trend similar to that observed for 2erk 2P.MKI compared to the other 2erk 2p states. The most dramatic shift in the distribution was observed for 5umo 0P.FL, which shifted *φ* to higher angle with a probability density that was significantly broadened compared to the others. As noted above, the dihedral angle for each frame varied considerably when plotted over the course of the 1° seed trajectory where 5umo 0P.FL appeared (**Fig. 10B**), even after the A-loop reached its settled conformation (**Fig. 7B**). Density plots of the distance between Q60 and Y231 which measures N- and C-lobe separation (**Fig. 10E**), as well as frames captured during this trajectory (**Fig. 10F-H**) illustrate the large rotations of helix *α*C underlying this variation. Together, the results reveal broader motions within the kinase core in states of 0P-ERK2 compared to 2P-ERK2, and especially large backbone motions in 5umo 0P.FL.

#### Variations in RMSF and active site residue distances between 0P- and 2P-ERK2

RMSF values for C*α* atoms were examined for the different states of 0P-ERK2 (**Fig. 11A**). Each state was normalized to 2erk 2P.L16, in order to directly compare them to the RMSF plots in 2P-ERK2 (**Fig. 6C**). Outside of the A-loop and MKI segments, nearly all states of 0P-ERK2 (5umo 0P.MKI, 2erk 0P.Y-*α*C, and 2y9q 0P.F/Y) showed RMSF values that were largely comparable across the kinase (**Fig. 11A**). The exception again was 5umo 0P.FL, which displayed RMSF values elevated far above the others, revealing higher dynamics in all regions of the enzyme. Importantly, larger fluctuations in the N-lobe and the L16 segment were observed in all forms of 0P-ERK2 relative to 2erk 2P.L16, and were comparable to fluctuations in 2erk 2P.pY-R65 and 2erk 2P.MKI (**Fig. 6C**). Although the L16 segments in these states appeared organized, greater variability in associated L16 side chain interactions with the N-lobe (e.g. F327) were apparent (**Fig. 11B-E**). The results are consistent with a model in which disrupting interactions of the A-loop with L16 elevates the RMSF in key regions of the N-lobe, enabling conformations of the A-loop to control dynamics in the active site. Overall, the range of motions in 0P-ERK2 appeared systematically enhanced relative to 2P-ERK2.

**Figure 11.**
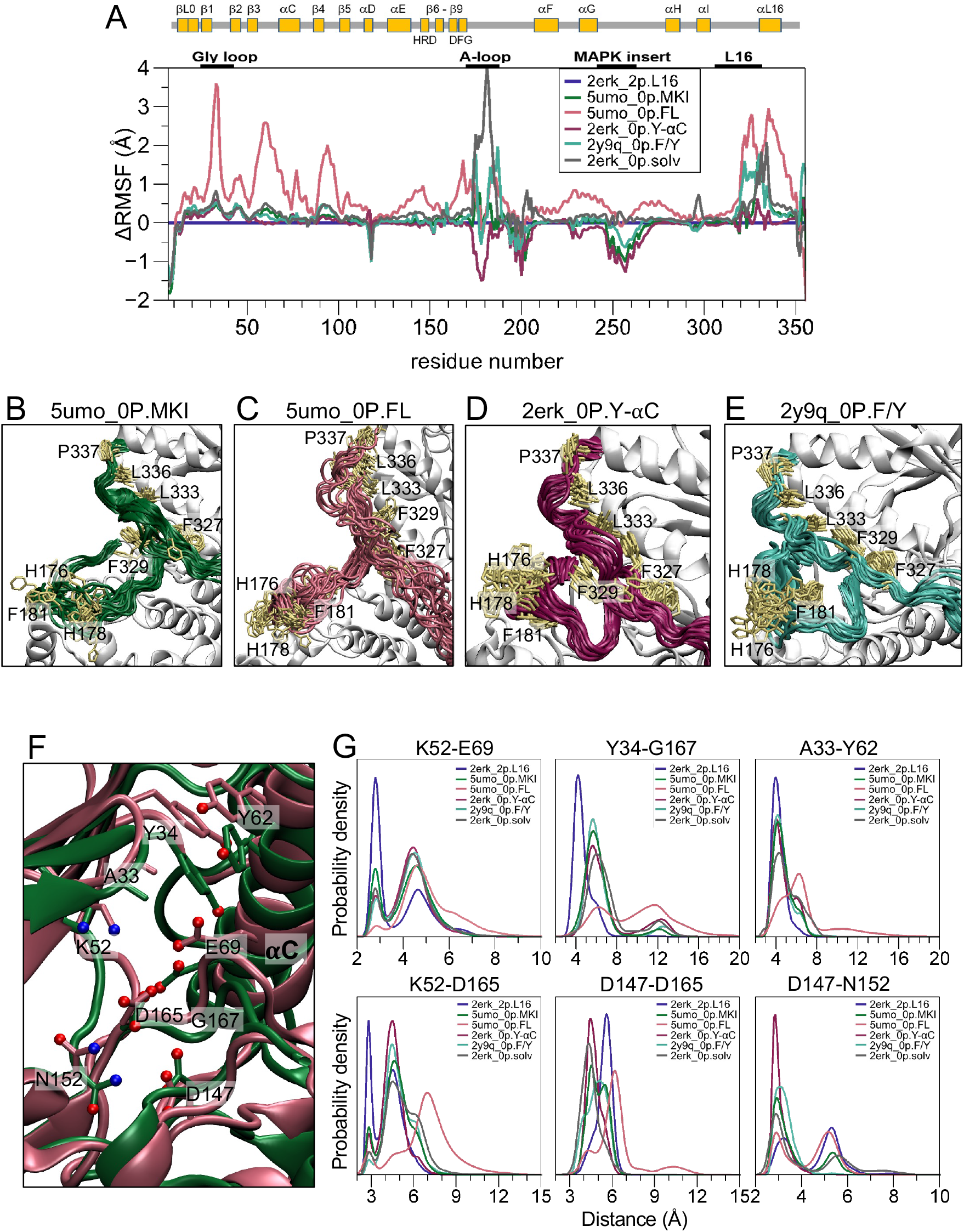
Differences in regional dynamics between settled A-loop states in 0P-ERK2. **(A)** Differences in RMSF between states, normalized to the 2erk 2P.L16 settled state (horizontal purple line). **(B-E)** Overlays of frames (every 250 ns) showing regions of the A-loop and L16 segment for settled states **(B)** 5umo 0P.MKI (5umo 0P 1° seed 1 300 K), **(C)** 5umo 0P.FL (5umo 0P 2° seed 4 300 K), **(D)** 2erk 0P.Y-*α*C (2erk 0P 1° seed 1 315 K), and **(E)** 2y9q 0P.F/Y (2y9q 0P 1° seed 3 300 K). **(F)** Overlay of reference structures for 5umo 0P.MKI (green) and 2erk 0P.Y-*α*C (maroon), highlighting residues within the active site, as in Fig. 6A. **(G)** Probability densities for distances between residue pairs; each curve integrates to 1.0 for the region shown. Shifts in probability density reveal longer distances involving catalytic residues in states from 0P-ERK2, referenced to 2erk 2P.L16.

Accordingly, density plots revealed lengthening of distances between residue pairs in the N-lobe of 0P states relative to 2erk 2P.L16 (**Fig. 11F,G**), as observed with the other states of 2P-ERK2 (**Fig. 6A,B**). Thus, the K52-E69 salt bridge showed a significant shift to longer distances in 5umo 0P.MKI, 2erk 0P.Y-*α*C, 2y9q 0P.F/Y, and 5umo 0P.FL. Similarly, lengthening between Y34-G167, A33-Y62, and K52-D165 was observed in all 0P states relative to 2erk 2P.L16. Notably, 5umo 0P.MKI, 2erk 0P.Y-*α*C, and 2y9q 0P.F/Y showed populations that were mostly comparable to those seen in 2erk 2P.pY-R65 (**Fig. 6B**, **Fig. 11G**), while contacts between all residue pairs were even more strongly disrupted in 5umo 0P.FL. This revealed larger conformational fluctuations in 5umo 0P.FL, including for example a population of conformers whose Y34-G167 distances ranged between 10 Å and 15 Å, corresponding to an autoinhibited state with the Y34 side chain folded underneath the Gly loop (**Fig. 10G**). Overall, 5umo 0P.FL and other A-loop states of 0P-ERK2 revealed greater disruption of residue contacts in the active site, compared to 2P-ERK2.

In summary, the results from MD show substantial differences between A-loop states sampled by 0P- and 2P-ERK2. In all simulations of 2P-ERK2, the A-loop moves away from the 2ERK X-ray conformation, forming settled conformers with variable contacts to the kinase core. The differences between these contacts lead to differences in dynamics in regions surrounding the ATP binding site, and suggest dynamic restraints introduced through A-loop interactions with the L16 segment. By contrast, simulations of 0P-ERK2 maintain starting conformations and introduce new ones. Many of the settled states deviate from the 5UMO X-ray structure, resulting in disrupted A-loop contacts with MKI and solvent-exposure of Y185. Importantly, these 0P states reflect a lower degree of organization and reduced dynamic restraints in the N-lobe, compared to 2P-ERK2.

### Survey of crystal structures of 0P- and 2P-ERK2

Two striking outcomes of the MD analysis were the persistent movements of the A-loop away from the 2ERK starting state in all trajectories of 2P-ERK2, and the significant variations of the A-loop in 5umo 0P.FL and other 0P settled states. Sang et al. [57] have proposed that crystal packing interactions stabilize the A-loop conformation in the 5UMO crystal structure. Furthermore, recent MD simulations of p38*α* MAP kinase [27] and Abl kinase [1] also found large variations in solution-phase A-loop conformations; both studies established the role of crystal packing in stabilizing/selecting respective A-loop conformers using simulations of the crystal environment. Therefore, we explored the conformational diversity of ERK2 in more detail, using a structural survey of the RCSB PDB. We would expect MD simulations that included the crystal environment to maintain contacts observed by X-ray diffraction even at elevated, room temperatures, as observed in the above-mentioned studies [1, 27].

**Table S2** provides a summary of all ERK2 crystal structures surveyed for this study, augmented by crystallographic metadata and stringified neighbor contacts in **Suppl Dataset S1**. Included are 10 structures of 2P-ERK2 and 167 structures of 0P-ERK2, along with RSCB metadata and all crystal contacts formed with the A-loop or MKI. For all PDBs, each chain within the asymmetric unit was treated as an independent entry, in keeping with the parametric treatment (coordinates, occupancies, and temperature factors) by the crystallographer, all totaling to 186 chain entries.

Crystal contacts found in the 2P-ERK2 apoenzyme (2ERK) are shown in **Fig. 12A**, highlighting crystal lattice atoms within 5 Å of heavy atoms in the structure. Except for 2ERK, all 2P-ERK2 structures were ligand-bound, either to a small molecule or a polypeptide. In four of five ligand-bound structures (5V60, 6OPG, 6OPH, 6OPK), the resolved A-loop was modeled by the same solvent-exposed conformation described in 2ERK. Like 2ERK, each structure displayed multiple crystal contacts with A-loop residues P174-F181, despite differences in space groups. One structure of 2P-ERK2 bound to an ATP-competitive inhibitor (6OPI) showed a disordered A-loop, in a space accessible to solvent. Thus, the similarities in crystal contacts found in the majority of 2P-ERK2 structures suggest stabilization of the A-loop conformation by crystal packing, which is supported by the observations that all MD trajectories rapidly deviated away from the canonical A-loop structure (**Fig. 2**). Notably, 2P-ERK2 cocrystallized with a polypeptide bound to the ERK DEF docking site (PEA15 death effector domain (DBD), PDBid: 4IZA) showed an A-loop that was solvent-protected by multiple PEA15-DBD interactions [32]. Here, the A-loop formed a conformer that broke the canonical salt bridge between pY185, R189 and R192, and moved pY185 away from the C-lobe to interact with R65 on helix *α*C. The similarity between this A-loop movement and the 2erk 2P.pY-R65 settled state further supports the latter as an accessible conformation of 2P-ERK2 that can accommodate protein interactions with the DEF docking site.

**Figure 12.**
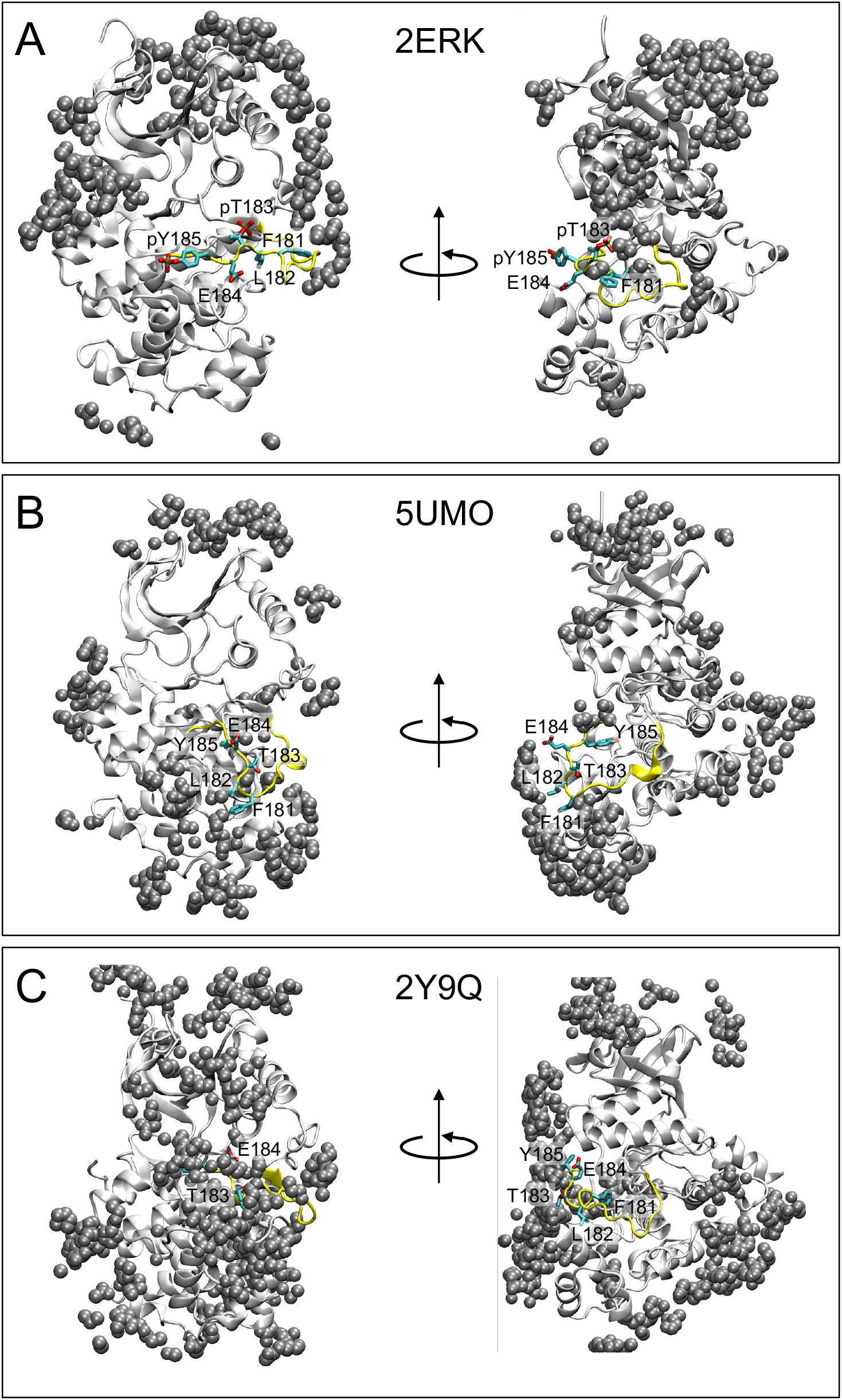
Crystal contacts with the A-loop in X-ray structures of ERK2. Structures of **(A)** 2ERK, **(B)** 5UMO, and **(C)** 2Y9Q, with the ERK2 asymmetric unit rendered in white cartoon, and atoms from protein neighbors within 5 Å shown as black spheres. Alternative orientations correspond to rotation by ∼90° about the vertical axis.

In the 5UMO crystal structure of 0P-ERK2, Y185 is buried in a pocket formed by I196, I207, and R146, and stabilized by interactions of F181 and L182 with L232 and Y261, located in the helix *α*G-MKI pocket. **Fig. 12B** highlights crystal lattice contacts with A-loop atoms in 5UMO. These reveal extensive contacts with residues in the pocket between helix *α*G and the helix-turn-helix of MKI (I254-R259) and 16 atoms on neighboring ERK2 molecules. Notably, the size of the pocket for F181 and L182 is extended by crystal contacts with Y315 and P317 from an ERK2 neighbor. At the same time, the A-loop residues preceding F181 (H178-T179-G180) are in close proximity to multiple atoms in T157 and T158 from a second ERK2 neighbor. By computing the intersection between the crystal contacts in 5UMO and those in all other entries, we observed that 112 entries retained *≥* 75% of the same contact pairs, all sharing the same spacegroup (P12_1_1) and A-loop conformation, and only one with a partially unresolved A-loop (**Suppl. Dataset S1**). Seven additional entries displayed the same A-loop conformation, except that A-loop residues H176-G180 formed contacts with multiple atoms from S120 and D121 on helix *α*E from an ERK2 neighbor (space group P2_1_2_1_2). Two other entries (4QTE, 6G54, space group P3_2_21) displayed the same A-loop conformation, but with multiple contacts with the MKI from an ERK2 neighbor. Therefore, the majority of 0P-ERK2 structures shared crystal contacts in the regions surrounding the A-loop and *α*G-MKI pocket that might be expected to stabilize the observed A-loop conformation.

By contrast, in 36 PDB entries (**Suppl. Dataset S1**) the A-loop of 0P-ERK2 was partially or completely disordered (e.g. 5K4I, 4QTA, 6OTS). Of these, only one (4XOY) displayed F181 and L182 interactions with MKI that were similar to those in 5UMO. In all others, interactions between the A-loop and its canonical binding site in 5UMO were replaced, either by other residues within the kinase core or by crystal contacts with a neighboring molecule. In 29 cases, crystal contacts with the MKI distorted the *α*G-MKI pocket, precluding interactions with with F181 and L182 (e.g. 2Y9Q, **Fig. 12C**).

In summary, crystal packing interactions appear to strongly influence the A-loop conformations of both 2P-ERK2 and 0P-ERK2, either by stabilizing intramolecular interactions of the A-loop with the kinase core, or by interfering with them. Together, they support the indications by MD of variable A-loop conformations as reasonable representations of solution behavior.

## Discussion

In this study, extended conventional molecular dynamics simulations reveal unexpected conformational heterogeneity of the A-loop in both 2P- and 0P-ERK2. At least three settled states of the A-loop can be observed in 2P-ERK2, each diverging appreciably from the starting 2ERK crystal structure. At least four states can be observed in 0P-ERK2, three of which disrupt canonical A-loop interactions with the C-lobe seen in the 5UMO crystal structure. Importantly, the different A-loop states are associated with distinct effects in regions within the kinase core. In particular, variations in the range of motions in the active site are observed. Overall, 2P-ERK2 shows greater dynamic restraint and compactness between conserved residues in the N-lobe and active site, in contrast to 0P-ERK2 which shows a higher level of disorganization. Our findings suggest that dual phosphorylation at T183 and Y185 serve in part to restrain N-lobe dynamics in the active form of ERK2, which may assist in catalytic turnover. The results from MD support conclusions reached from NMR relaxation dispersion and HX-MS experiments, that motions of the A-loop can be coupled to motions at the active site.

In 2P-ERK2, the A-loop conformation varies widely among the three states that can be distinguished. One state (2erk 2P.L16) forms multiple side chain contacts from the A-loop to the L16 segment in the N-lobe, while another (2erk 2P.MKI) disrupts these interactions with L16 and forms strong contacts with the C-lobe and MKI. Remarkably, a third conformation (2erk 2P.pY-R65) breaks the salt-bridges between pY185, R189 and R192 in the 2ERK crystal structure, releasing pY185 to move towards the N-lobe, while R189 moves towards the C-lobe and shifts the position of the loop between helices *α*F and *α*G. The settled states observed in key trajectories imply thermal accessibility of these states as well as a more dynamic state (2erk 2P.solv) that connects them. This suggests that the salt bridges formed between the two phosphorylated residues and multiple Arg residues in the 2ERK X-ray model are in fact not highly stable, possibly reflecting their exposure to solvent.

In 0P-ERK2, two states remain similar to their starting crystal structures (5umo 0P.MKI, 2y9q 0P.F/Y), while two other states move the A-loop to very different conformations (5umo 0P.FL, 2erk 0P.Y-*α*C). Noteworthy is the settled state, 5umo 0P.FL, that transitions from 5umo 0P after substantial disruption of contacts between A-loop residues F181 and L182 and the helix *α*G-MKI pocket. Following this trajectory (**Fig. 9A,B**) illustrates how the F181 side chain initially undergoes reversible excursions away from the pocket while L182 remains docked, and then the new settled A-loop forms when L182 moves away from the C-lobe and connects with the P+1 segment. The disruption of F181 and L182 contacts with the C-lobe, as seen in 5umo 0P.FL, 2y9q 0P.F/Y, and 2erk 0P.Y-*α*C, in each case moves Y185 away from its buried position in 5UMO and exposes it to solvent. This may explain why Y185 appears more inaccessible than T183 in the crystal structure, yet is kinetically favored for phosphorylation by MKK1/2 [18].

Such behavior suggests the importance of F181 and especially L182 in remodeling the A-loop. Like 0P-ERK2, contacts with F181 and L182 also control A-loop conformational states in 2P-ERK2. Thus, L182 forms close interactions with R170 and Y203 in 2erk 2P.pY-R65, while F181 is solvent exposed. Likewise, L182 again interacts with Y203 in 2erk 2P.MKI, and packs against P174 and side chain methyls in A172 and T179. By contrast, F181 forms contacts with the L16 loop in 2erk 2P.L16, at the cost of exposing L182 to solvent. Interestingly, 2° seed trajectories show *µ*s timescale decay of the 2erk 2P.L16 state more often than 2erk 2P.MKI or 2erk 2P.pY-R65. Conceivably, L182 may be an important contributor to the lifetime of the settled state and a driver for conformational exchange, where breaking and making contacts with this residue underlies A-loop remodeling.

These novel settled states of 2P- and 0P-ERK2 highlight the advantages of using extended conventional MD simulations to explore conformational changes. Previous MD studies on 0P-ERK2 [31, 57], carried out using trajectory lengths of (0.5 to 1) *µ*s, showed no major changes in the A-loop from the initial crystal structures, even when hundreds of parallel runs were conducted totalling to 2 ms [57]. In our simulations, 1 *µ*s was never long enough to capture deviations from the initial state into new settled states. The longer continuous individual trajectory run times, together with a change of the water model from TIP3P [21] to OPC [59] in our study may have helped alleviate biases introduced by crystal lattice contacts in the starting models, as suggested by Tian et al. [65].

The large quantities of simulation data and the mixing of A-loop conformational states arising at different temperatures necessitated new structural analysis tools to classify states and identify them across multiple trajectories. To achieve this, we used the varying fluctuations of the loop as an intuitive identifier and then applied a “native contact” parameter (Q), developed for protein folding, to refine and separate. The application *Q_A−loop_* allowed large collections of frames to be separated as a function of A-loop conformation. In doing so, it established the important dynamic role of the A-loop.

Correlations between residue distances within the active site and states of the A-loop were revealed by dCNA after applying the *Q_A−loop_* metric. In 2P-ERK2, greater compaction between catalytic residues in the N-lobe was observed in 2erk 2P.L16, which was accompanied by reduced fluctuations across the Gly loop, *β*1-*β*5, and helix *α*C. Although small shifts to longer distances were seen in the other states of 2P-ERK2, the active sites in all forms remained structured. By contrast, uniformly longer distances and larger fluctuations were seen in states of 0P-ERK2, increasing to levels as high as RMSF > 3 Å in trajectories of 5umo 0P.FL. Together, the results link the conformational states of the A-loop to the dynamics of the active site. Notably, stronger A-loop interactions with L16 are associated with lower N-lobe dynamics in 2P-ERK2, while states of 0P-ERK2 reflect higher dynamics in general. That 2P- and 0P-ERK2 differ in their active site dynamics suggests that motions of the A-loop underlie a dynamical switch between active and inactive states of ERK2. This leads us to a working model in which salt bridges formed by phosphorylation promote residue contacts between the A-loop and L16 to increase compactness and restrain dynamics within the active site, while loss of phosphate allows larger A-loop movements that release these dynamic restraints. Thus, we propose that the switch between inactive and active states of ERK2 involves the control of active site dynamics.

Our MD results expand the structural understanding of dynamics in ERK2 inferred from solution measurements. Previous NMR relaxation dispersion studies of 2P-ERK2 demonstrated conformational exchange of the A-loop between thermally accessible states (designated “R” and “L”), differing by only 3.3 kJ/mol (0.8 kcal/mol) [69]. Furthermore, both NMR and HX-MS experiments revealed coupling between the A-loop and the active site, based on mutations in the A-loop that blocked exchange within the active site, and inhibitor binding to the active site which altered hydrogen-deuterium exchange adjacent to the A-loop [20, 42]. Although our simulations reveal multiple “long-lived” conformational states that might undergo exchange, the (10 to 100) *µ*s timescale implied by the trajectories is one to two orders of magnitude faster than the millisecond timescale associated with L⇌R exchange. Most likely, the conformational states observed in this study are not the only ones that exist in 2P-ERK2. Instead, we propose that the R state in 2P-ERK2 may be represented by an ensemble of organized states with relatively restrained active site dynamics, while the L state in 0P-ERK2 may be represented by a more disorganized set of states. The latter includes 5umo 0P.FL, which disrupts the catalytic salt bridge, K52-E69, and shifts K52-D165 and D147-D165 towards longer distances apart (**Fig. 11G**). Lengthening of contacts to D165 reflects a rotation of the F166 side chain in the DFG motif partially towards a “DFG-out” conformation. Lengthening of the Lys-Glu salt bridge and rotation of the DFG backbone are widely regarded hallmarks of inactive kinases [36, 64]. That 5umo 0P.FL may be a major solution structure of 0P-ERK2 representing its true “inactive” form is supported by HX-MS measurements showing less protection from hydrogen-deuterium exchange in the A-loop and P+1 segment of 0P-ERK2 compared to 2P-ERK2 [42]. This is consistent with the higher solvent exposure seen in 5umo 0P.FL compared to the other states of 0P-ERK2 or any of the 2P states [5, 67]. Finally, the lack of conformational exchange in 0P-ERK2 seen by NMR [69] is consistent with the patterns of disorganization between contacts in dCNA analyses of 5umo 0P.FL (**Fig. 9C-E**, **Fig. 10F-H**).

The functional importance of A-loop dynamics is an emerging concept in the kinase field [41]. Like ERK2, recent solution measurements of other kinases have revealed multiple substates of the A-loop available to both active and inactive forms [15, 16, 27, 33, 53, 70]. In Aurora A, single molecule fluorescence quenching and Forster resonance energy transfer (FRET) experiments show shifts in A-loop populations between active, open *vs* inactive, closed states, which are respectively coupled to active DFG-in *vs* inactive DFG-out conformations [15, 16, 53]. Conformational selection for the open A-loop by Aurora A inhibitors correlates with binding of an allosteric activator, TPX2. This suggests that the availability of A-loop substates in the inactive, unphosphorylated form of Aurora A enables its binding and activation by TPX2. Similarly, site-directed spin labeling EPR measurements of CDK2 reveal heterogeneous populations of the A-loop in open vs closed states, which correlate with active helix *α*C-in *vs* inactive *α*C-out conformations [33]. Transient formation of the open A-loop conformation enables allosteric recognition and binding of cyclin A to promote CDK2 activation [33]. The Abl tyrosine kinase provides a third example, where type II kinase inhibitors promote conformation selection for a DFG-out/A-loop-closed configuration, which in turn disrupts SH3-linker interactions leading to detachment of the SH3-SH2 domains and disassembly of kinase domain interactions [62].

In the same way, conformational variants of the A-loop in ERK2 might expand the range of substrates or effectors that bind with conformation selection, either through direct contacts with the A-loop or through propagation from distal allosteric sites. For example, the switch in pY185 interactions from R189/R192 in the C-lobe to R65 in the N-lobe in 2erk 2P.pY-R65 is in line with the 2P-ERK2:PEA15 cocrystal structure, which features disruptions of both the canonical pY and pT salt bridges [32]. Thus, structural and flexibility changes near pY185 and the MKI might affect recognition of substrates and effectors, especially those that bind the DEF docking motif site formed between helix *α*G and MKI. Models proposed for substrate binding to the DEF site invariably involve interactions with hydrophobic residues positioned by the canonical A-loop fold in 2ERK [7, 30, 45, 46, 61]. Movement of the A-loop to allow contacts with helix *α*C suggests that plasticity of the A-loop fold should also be considered in models for substrate binding. For example, the explanation of why DEF ligands such as ELK1 show reduced binding to 0P-ERK2 [6] should be reevaluated to include A-loop flexibility. Conceivably, the assumption that the DEF binding site is created only when the enzyme is phosphorylated [30] may be incorrect. Likewise, malleability of the A-loop has implications for QM/MM models of catalysis, which up to now have inferred proximity of pY185 to substrate in the pre-chemistry conformation ensemble [14, 66].

The suggestion by our study that motions within the ATP-binding site can be coupled to A-loop motions by communication through the L16 segment and helix *α*C may inform our understanding of binding and dissociation rates for nucleotides and small-molecule inhibitors [63]. Rigidification of target enzymes has been associated with slow-onset/slow-offset inhibition [40]. In Abl kinase, metadynamics was used to show how a resistance mutation could alter protein flexibility and shorten the residence time of imatinib inhibitor [60]. In ERK2, tight-binding inhibitors Vertex-11e and SCH772984 have been shown to modulate the conformational equilibrium between the energetically similar R and L states, in a manner associated with differences in the dissociation rate constant [52]. Furthermore, conformational selection by these ERK inhibitors has been shown to modulate the rate of 2P-ERK2 dephosphorylation by MAPK phosphatase 3 (MKP3/DUSP6) [42], providing a way for ATP-competitive inhibitors to control ERK inactivation by regulating exchange between A-loop populations. The ability of active ERK2 to control the flexibility of the active site by toggling between multiple conformations of the A-loop presents an intriguing new behavior to exploit for drug design.

In addition to the potential regulation of enzyme turnover, our findings of new A-loop conformations for 2P-ERK2 could have implications for alternative functions of ERK2. Even in its “kinase-dead” mutant state (K52R), 2P-ERK2 has been reported to allosterically activate topoisomerase II*α* and bind DNA in a manner that requires dual phosphorylation [26, 49], suggesting that the A-loop fold might accommodate noncatalytic functions of ERK2. Together, these observations suggest that A-loop motions in ERK2 could help explain its multifunctional properties and broad recognition of substrates and effectors.

Analyses of the A-loop conformations and crystal contacts in the 0P-ERK2 PDB entries lead to several insights. In each case, the canonical A-loop structures seen in 2ERK and 5UMO are accompanied by crystal packing interactions that may stabilize the fold. In 0P-ERK2, the majority of structures of wild-type, mutant, and ligand-cocrystallized forms share the same space group (P12_1_1), and without exception, display the canonical A-loop conformation. Those with other crystallization symmetries showed alternative A-loop conformations or missing electron density. Like ERK2, MD simulations of Aurora A and p38*α* MAP kinase have also suggested that majority conformers in solution may differ from the loop states observed in crystal structures, due to packing interactions [27, 53]. Effects of crystal packing on protein structure more broadly [24, 74] and specifically on loop conformations have been demonstrated in other systems [47]. In fact, anisotropic displacement parameters (ADP) derived from an ensemble of myoglobin structures were found to agree well with the positional variance of the solution NMR ensemble but were suppressed by crystal contacts [24]. Crystallization co-solutes at high concentrations may also shift conformational equilibria due to preferential interactions with varying molecular surfaces [23, 43], as observed experimentally for solvent-exposed loops [22].

On the other hand, structures of 0P-ERK2 with unresolved A-loop regions suggest that the loop can be flexible. The position of the MKI appears to be an important determinant of the A-loop conformation, where crystal contacts that stabilize the MKI closer to helix *α*G block canonical loop formation, while contacts that stabilize the canonical fold open and extend the pocket to allow insertion of F181 and L182. Most of the 10 deposited structures of 2P-ERK2 model the A-loop as in 2ERK. However, the unresolved A-loop in two inhibitor-bound structures (4QTB, 6OPI) and the alternative loop conformation in PEA15-bound 2P-ERK2 (4IZA)[10, 32] suggest that other loop conformers are possible in solution. Thus, for both unphosphorylated and phosphorylated ERK2, crystallographic evidence exists to support the flexibility of the A-loop and its ability to adopt multiple conformations.

## Materials and Methods

### Preparation of structural models

Heavy-atom models of the phosphorylated (2P) and the unphosphorylated (0P) forms of ERK2 were constructed from crystal structures PDBid 2ERK, 5UMO, and 2Y9Q using HackaMol [48]. All crystal water molecules were removed. Starting states from these structures after energy minimization, described below, are referred to as 2erk 2P, 5umo 0P, and 2y9q 0P throughout; a third form of 0P-ERK2 (2erk 0P) was constructed by removing the two phosphate groups from the 2erk 2P model. Amino acid sequences for all models were referenced to that of 2ERK, omitting the first five residues in the protein sequence (MAAAA) because they were unresolved in the crystal structure. The 2Y9Q sequence was converted from the human sequence to rat, by renumbering residues (resids) and introducing a single V44L mutation using Chimera [44]; the alternative location (altloc) setting “A” was selected for all residues with multiple occupancies, which was relevant only for 2Y9Q. Missing N- and C-terminal residues of 5UMO (resids 11-14 and 355-358) and 2Y9Q (resids 6, 357 and 358) were grafted from the 2ERK structure using superposition to neighboring backbone atoms. Residues were named according to the AMBER forcefield, with protonation states assigned (by name) for pH 7. Histidine was determined to be the only ambiguous residue. The singly protonated His (HIE and HID) and doubly protonated His (HIP) forms were assigned by searching for non-carbon neighbors within 3 Å of the ND1 and NE2 atoms. Potential ring flips (HIE *vs* HID) during crystal modeling were considered but determined to be unnecessary; all His assignments (with the exception of H178 in 2erk 2P) were manually inspected and applied identically to all models. In each model, H139 was assigned as HIP, due to its proximity to D208. The H178 residue was assigned as HIP in 2erk 2P, due to an observed salt-bridge with E332, and assigned as HIE in 5umo 0P, 2y9q 0P, and 2erk 0P. Overall, each model contained 353 amino acids, with 5umo 0P, 2y9q 0P, and 2erk 0P being chemically identical, and 2erk 2P adding phosphates to residues T183 and Y185.

The Amber LEaP program was used to construct the starting points for each model. Each model was treated with the ff19SB force field [65] and immersed in a truncated octahedral box of OPC water molecules [59]; the edge of the box was set to satisfy a minimum of 12 Å from any protein atom. Phosphorylated residue parameters were loaded (from leaprc.phosaa19SB) as described in the Amber20 manual. Each state was neutralized with Na^+^ (5 ions for 2erk 2P and 2 ions each for 5umo 0P, 2y9q 0P, and 2erk 0P). A NaCl concentration of ∼150 mmol/L was imposed by adding 54 Na^+^ and 54 Cl*^−^* ions (using the Amber LEaP addIonsRand function); the Na^+^ and Cl*^−^* ions were treated with the Li/Merz monovalent ion parameters for the OPC water model (frcmod.ions1lm 126 hfe opc) [59]. There were 15,565; 15,576; 15,555; and 15,571 water molecules for 2erk 2P, 5umo 0P, 2y9q 0P, and 2erk 0P, respectively.

### Molecular dynamics simulations

All calculations were carried out using the GPU-enabled CUDA version of the pmemd executable (pmemd.cuda) in AMBER Version 20 [9, 29, 56]. For 2erk 2P, 5umo 0P, and 2erk 0P, two independent production runs were carried out at 300 K and 315 K, and one run was carried out at 285 K and 330 K. Each of these consisted of (15 to 27) *µ*s of continuous simulation and are referred to as “1° seeds”. For 2y9q 0P, four independent seeds were run independently at 300 K for 9.9 *µ*s of continuous simulation. Nonbonded interactions were treated using a 9 Å cutoff; long-range van der Waals interactions were approximated using the Amber2020 default setting (vdwmeth = 1); long-range electrostatics were calculated using the particle mesh Ewald method [11]. The SHAKE algorithm was applied to all bonds including hydrogen to allow the 2 fs timestep [35, 55]. All starting points (2erk 2P, 5umo 0P, 2y9q 0P, and 2erk 0P) were minimized with a 10.0 kcal/mol/Å^2^ (1 kcal/mol/Å^2^ = 418.4 kJ/mol/nm^2^) restraint on all protein atoms using up to 15,000 steps of steepest descent. Heating and equilibration were carried out in a series of steps: first, each system was heated to 285 K for 2 ns and then run at 285 K for another 7 ns with the temperature maintained using a Langevin thermostat with a collision frequency of 5 ps*^−^*^1^. Next, a series of NPT equilibration steps were used to gradually reduce the protein-position restraints in four 10 ns steps; the first step retained the 10.0 kcal/mol/Å^2^(1 kcal/mol/Å^2^ = 418.4 kJ/mol/nm^2^) position restraint on all protein atoms, and the next three steps applied restraints to the protein backbone atoms only, at 10 kcal/mol/Å^2^, 1 kcal/mol/Å^2^, and 0.1 kcal/mol/Å^2^ (1 kcal/mol/Å^2^ = 418.4 kJ/mol/nm^2^). Each system was run without restraints at 285 K for another 310 ns and then brought up to the production temperatures in 5 K increments each for 5 ns (285 K *→* 290 K *→* 295 K *→* 300 K *→* 305 K *→* 310 K *→* 315 K *→* 320 K *→* 325 K *→* 330 K). For 2y9q 0P, four seeds were run independently at 285 K and then heated to 300 K in 5 K increments for 5 ns. For 2erk 2P, 5umo 0P, and 2erk 0P, the additional production runs at 300 K and 315 K were equilibrated with a slightly different protocol, where the system was brought up to 300 K during the initial heating stage, and the 315 K runs were started from the 300 K runs after 10 *µ*s, raising the temperature in 5 K increments for 5 ns. All NPT production runs used a Langevin thermostat with a collision frequency of 5 ps*^−^*^1^ to maintain the temperature, and a Monte Carlo barostat with a coupling constant of 2 ps to maintain the pressure at 1.01325 bar.

Additional seeds were run at 300 K, initiated from activation loop conformational states observed in the ten 1° seed production runs described above. The initial coordinates for each new seed were taken from restart files corresponding to the temporal region of the A-loop conformation; generally, the initial configuration of each consecutive new seed was taken from restart files separated by 300 ns with respect to the original trajectory. These are referred to as “2° seeds”. For 2erk 2P, eight 2° seeds were started from a long-lived conformation found in the 1° seed at 330 K; 14 seeds were started from a conformation at 285 K; and 8 seeds were started from a conformation at 300 K. For 5umo 0P, five 2° seeds were started from a conformation at 315 K where the A-loop deviated from the X-ray structure. For 2erk 0P, 8 seeds were started from a conformation at 315 K. Each 2° seed for 2erk 2P and 2erk 0P was run for 5.70 *µ*s while the secondary seeds for 5umo 0P ranged between 4.92 and 6.30 *µ*s. There were no 2° seeds run for 2y9q 0P. The total accumulated production sampling for all forms of ERK2, summed over all temperatures (285 K to 330 K) and seeds was 727 *µ*s (**Table 1**).

In general, for both 1° and 2° seeds, each trajectory was run continuously, 300 ns at a time, and saved every 100 ps (i.e., every 50,000 steps). File system problems corrupted frames for 13 individual, 300 ns trajectories of the 1° seeds (7 in March 2021 and 6 in June 2021). The skipbadframes function of CPPTRAJ [51] was used to remove these frames with respect to integrity checks on all 353 protein residues (“check: 1-353 skipbadframes” where resid 1 in the models corresponds to resid 6 for the canonical 2ERK sequence). There was no issue apparent for the restart files. Examples in **Fig. S12A** show total energy plots at different temperatures before and after restarts. For visualization (in VMD) and analysis, all trajectories were stripped of water and NaCl, downsampled to 2.5 ns between frames, and aligned to the respective, state-appropriate minimized 2erk 2P and 2erk 0P structures using the C, CA, and N backbone atoms excluding the activation loop (resids 170 to 186) and the ten residues at the N- and C-termini (resids 6-15 and 349-358).

### Activation loop conformer search and classification *via* native contacts

Long-lived (*≥* 5 *µ*s) activation loop conformations were identified in the simulations that started from 2erk 2P and 2erk 0P. These were found by splitting each 1° seed trajectory into 1 *µ*s segments and calculating the RMSF of the activation loop C*α* atoms (resids 170-186) for each segment. Any contiguous segment *≥* 5 *µ*s with an average A-loop RMSF *≤* 1.2 Å was indexed as a “settled” A-loop conformer. New 2° seed trajectories at 300 K (described above) were initiated from frames in these segments. These criteria worked well for 2erk 2P and 2erk 0P, but were not relevant for the majority of 5umo 0P and 2y9q 0P runs, where the A-loop remained close to their crystallographic conformations with low RMSD. The additional A-loop conformation discovered in seed 2 for 5umo 0P at 315 K (5umo 0P.FL) was *≤* 1.23 Å for two 2 *µ*s segments connected by a 1 *µ*s segment with an average RMSF of < 1.5 Å; the selection criteria were relaxed slightly for 5umo 0P.FL due to clearly settled region (**Fig. 7B**).

The A-loop conformation reference structures were defined using coordinate averaging, followed by minimization of the RMSD from the averaged structure to identify the reference frame for the trajectory. Secondary seed trajectories exhibiting the A-loop conformer for the length of the trajectory were used for 2erk 2P and 2erk 0P due to the presence of multiple A-loop conformers across seeds; the single seed with the lowest averaged A-loop RMSD (from the starting frame) was used to calculate the averaged protein structure. For the 5umo 0P and 2y9q 0P states, all 300 K seeds were used to calculate the averaged protein structure, which was then used to scan all trajectories for the reference frame.

The collective fraction of native contacts for the activation loop, *Q_A−loop_* was used to quantify and further refine the long-lived conformational states according to the following equation [4]:

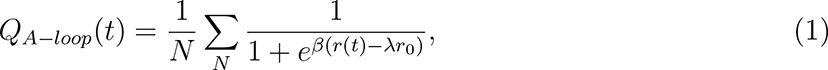

where *N* is the number of atom-atom pairs (within 4.5 Å) between residues in the A-loop (resids 170-186) and the remainder of the protein in the reference structure (excluding neighboring resids 169 and 187); *r*_0_ is the distance for a given reference pair of atoms; and *r*(*t*) is the corresponding distance at any time *t* in the trajectory (*β* = 5 Å*^−^*^1^ and *λ* = 1.8). Intuitively, *Q_A−loop_* ranges from 0 to 1. In Eq. 1, for *λ* = 1.8, a contact distance will contribute ∼ 1 to the sum while the sampled distance is less than the reference distance (i.e., the contact is present); the contribution drops to 0.5 as the sampled distance lengthens to 1.8 times the distance, and then further lengthening reduces the contribution to zero (i.e., the contact is not present). The sharpness of the transition is governed by the *β* parameter. Summing over all contacts and normalizing by the preceding 1*/N* yields a value between 0 (no reference contacts present) to 1 (all reference contacts present). For time-independent analyses, all trajectories (1° and 2° seeds) at 300 K were combined and aligned to either the 2erk 2P or 2erk 0P structures, and then split into A-loop conformational state collections using a threshold for *Q_A−loop_* of 0.67. This threshold was selected as a conservative estimate based on visual inspection of the *Q_A−loop_* charts; if a frame fell below this threshold for all 2P (or 0P) reference structures, it was added to the 2erk 2P.solv (2erk 0P.solv) collection.

The calculated *Q_A−loop_* values for each settled state provided effective frame separation for all trajectories. For any overlapping frames (i.e. *Q_A−loop_ >* 0.67 for more than one conformer) precedence was given to states in the order 2erk 2P.MKI, 2erk 2P.L16, then 2erk 2P.pY-R65. There were a small number of overlapping frames compared to the total (115,046) between 2erk 2P.pY-R65 and the other two states (125 and 296 overlapping frames with 2erk 2P.L16 and 2erk 2P.MKI, respectively). There were 0 overlapping frames between 2erk 2P.L16 and 2erk 2P.MKI.

### Naming and color palette for conformational states

Here we summarize naming of initial states and branched A-loop conformational states used in MD simulations of 0P-ERK2 and 2P-ERK2. The names used for all 1° and 2° seeds combine the starting PDBid and the phosphorylation state in lowercase (2erk 2P, 5umo 0P, 2y9q 0P, and 2erk 0P). The resulting A-loop conformational states are enumerated as follows:

- 2P: 2erk 2P.MKI, 2erk 2P.L16, 2erk 2P.pY-R65, and 2erk 2P.solv
- 0P: 5umo 0P.MKI, 2erk 0P.Y-*α*C, 2y9q 0P.F/Y, 5umo 0P.FL, and 2erk 0P.solv

The color palette for the A-loop states was designed using an online resource [39] to be accessible for colorblind readers, and is provided in **Fig. S12B**.

### Contact Map Analysis

MDAnalysis was used to calculate the residue-residue contact matrices for trajectory segments (for time-dependent contact analysis) and for the state collections described above (for state-dependent contact analysis). For each frame, all distances (*≤* 4.5 Å) between heavy atoms for residue pairs were calculated using the capped distances method from the MDAnalysis Distances library. A contact between residues was counted once if any heavy atom pair was within the cutoff distance. The cutoff distance 4.5 Å was used based on recent validation studies [72]. The contact matrices had 353 rows and columns and were very sparse. For time-dependent analyses, the sparse matrices were saved for each 300 ns trajectory segment (over 2,400 segments) and indexed in a data frame. For state-dependent analyses, a single contact matrix was saved for each state frame collection.

Differences between residue-residue contact probability matrices were used to compare associated changes between states or regions of the trajectories. The associated contact probability matrices were computed from the residue-residue contact matrices normalized by the total number of frames (diagonal elements of the matrix), and accumulated across multiple trajectory segments. Two states were compared by subtracting the contact probability matrix for the initial state from that of the final state. Bar analysis R functions from the Bio3D difference contact network analysis (dCNA) project repository (https://bitbucket.org/xinqyao/dcna/src/master/) were used to generate VMD visualizations for residue-residue contact probability differences [13, 68, 72]. In the visualizations, probability differences less than 0.1 were ignored; blue (red) bars corresponded to increased (decreased) contact probability changes, going from the initial state to the final state; and the radius of the bar was proportional to the magnitude of the difference.

**Fig. S13** compares the dCNA bar plots using frames (saved every 0.100 ns) and to bar plots generated from down-sampled trajectories (saved every 2.5 ns) 5umo 0P.MKI, 2y9q 0P.F/Y, and 2erk 0P.Y-*α*C. The quantitative agreement (**Fig. S13**) validated the use of the down-sampled trajectories, which were more convenient.

### Structural Analysis

MDAnalysis was used to collect data frames of structural measurements for each down-sampled frame (2.5 ns between frames for continuous runs) over all states, seeds, and temperatures. Consistent with the bar analysis described above, the minimum heavy-atom distance between residue pairs was used to measure the separation between residues. Using the minimum heavy-atom distance has the advantage of simplifying the information encoded in measuring the distance; the caveat is that the identities of the atoms associated with the distance may change from frame to frame. In addition to residue-pair distances, the pseudo-dihedral angle between helices *α*C (resids 62-75) and *α*E (resids 120-140) was calculated for each frame. Two vectors, centered on the C*α* atoms for each helix and scaled to the length of the helix (computed as the distance between the first and last C*α* atom), were used to generate the four points for the dihedral angle.

### RCSB Survey

All Protein Databank X-ray crystal structures corresponding to the MAPK1 gene were identified using the Search API of the RCSB (Research Collaboratory for Structural Bioinformatics, https://search.rcsb.org)[3]. The Data API of the RCSB (https://data.rcsb.org/) was then used to retrieve the corresponding metadata containing sequence information (human or rat) in order to use the sequence information referenced to rat; other information was also cached (included bound ligands, resolution, and space-group). Queries of both APIs were carried out with the user agent provided by the Perl Mojolicious web framework (mojolicious.org). The structural data were collected in November 2021.

HackaMol Perl scripts were used to carry out the analysis of the crystal structures. The coordinates of the crystal contacts within 25 Å of the asymmetric unit were reintroduced by applying the symmetry information contained in each PDB file. A translated CrysFML [50] subroutine was used for transformations between Cartesian and fractional coordinates. The crystal coordinates were all stored as chain X in the output PDB to simplify the selection of atom groups. The crystal contacts within 5 Å of specific residues including the A-loop (resids 168-186) and the MAPK insert (resids 254-259) were determined. Only the ATOM record names (and phosphorylated residues, if present) were used in the crystal contact analyses (i.e., all water or other HETATM cosolutes were ignored). PDB files containing erroneous residue numbering with respect to the listed organism were manually identified and corrected. For PDB files with multiple independent chains of the ERK2 molecule in the asymmetric unit, the crystal contacts included all other chains identified, along with crystal neighbors (e.g. in 4QP1, the chain A crystal contacts included those from chains B and X; the chain B crystal contacts included those from chains A and X).

## Supplemental Materials

In support of the results presented here, reference structures, analysis dataframes, dCNA visualization states, trajectory seeds (downsampled at 2.5 ns between frames, total over 290,000 frames) and scripts used for analysis are available for download from https://data.nist.gov/od/id/mds2-2988. All 0P and 2P structures are aligned to the 2erk 0p and 2erk 2p minimized structures, respectively. The energy-minimized structure for 2erk 2p is very similar to that of 2erk 0p (backbone RMSD is 0.01 Å) and the crystal structure (2ERK, backbone RMSD is 0.06 Å).

## Note

This article is, in part, a contribution of NIST, and is not subject to copyright in the United States for the authors. Trade names are provided only to specify the source of information and procedures adequately and do not imply endorsement by the National Institute of Standards and Technology. Similar products by other developers may be found to work as well or better.

## Acknowledgments

LMP and NGA were supported by NIH award R35GM136392 (NGA). All simulations were carried out on NIST/MML GPU clusters CTCMS, Nisaba, and Simba. We are grateful to Andrew Reid, NIST, who maintains the CTCMS resources, and to Chris Muzny, NIST, who facilitated the simulation work on Nisaba and Simba, and provided valuable assistance identifying a filesystem storage solution for housing and accessing the associated large quantities of data. We are also indebted to Xin-qiu Yao, Georgia State University, for help adapting the dCNA workflow to accommodate sparse matrices.

