## Supporting Information for "Activation loop plasticity and active site coupling in the MAP kinase, ERK2"

Provided here are additional tables and figures in support of the main text along with descriptions of Supplementary Dataset ERK2_xtal_contacts.csv. Here is the itemized list of SI contents:


- **Description of Supplementary Dataset 1**, ERK2_xtal_contacts.csv
- **Table S1 -** Selected dCNA matrix elements
- **Table S2 -** Tabulated summary of **Supplementary Dataset 1**
- **Figure S1 -** Heavy atom contacts for A-loop residues in reference conformations for 2P-ERK2
- **Figure S2 -** $Q_{A-loop}$ and RMSD plots for 2P-ERK2 1° seed trajectories
- **Figure S3 -** $Q_{A-loop}$ plots for 2P-ERK2 2° seed trajectories
- **Figure S4 -** RMSD plots for 2P-ERK2 2° seed trajectories
- **Figure S5 -** Overlay plots for settled states of 2P-ERK2
- **Figure S6 -** Side-chain connectivity between L16 and the active site
- **Figure S7 -** Reference conformations and heavy atom contacts for settled states of 0P-ERK2
- **Figure S8 -** $Q_{A-loop}$ and RMSD plots for 1° seed trajectories of 0P-ERK2
- **Figure S9 -** $Q_{A-loop}$ and RMSD plots for 2° seed trajectories of 0P-ERK2
- **Figure S10 -** Temporal dCNA for the 5umo_0P.FL 1° trajectory
- **Figure S11 -** Temporal dCNA for the 2erk_2P.MKI 1° trajectory
- **Figure S12 -** Rare filesystem imposed errors and color palette used in figures
- **Figure S13 -** Comparison of dCNA bar analyses calculated with different numbers of frames

### Supplementary Dataset S1.

The file, ERK2_xtal_contacts.csv, provides the crystallographic metadata and string-encoded crystal packing interactions (within 5 Å) for the A-loop and six MKI residues (I254 to R259) in different X-ray structures of ERK2. **Table S2** summarizes this dataset, but with columns describing the enzyme state (e.g., 0P, 2P, ligands, peptide linking compounds) removed.

The following PDBIDs listed in ERK2_xtal_contacts.csv represent phosphorylated forms of ERK2:

- 2ERK
- 3ZUV
- 4IZA
- 5V60, 5V61, 5V62
- 6OPG, 6OPH, 6OPI, 6OPK

All other PDBIDs listed represent 0P-ERK2.

In ERK2_xtal_contacts.csv, selected crystal packing interactions are encoded in strings. For example, in the string "I254-H123_Y126_T157_T158", the residue before the dash represents the asymmetric unit residue (I254), and residues following the dash (H123_Y126_T157_T158) represent crystal packing residues within 5 Å. Only those asymmetric unit residues with crystal packing interactions are included and are separated by semicolons. For each entry, the string-encoded crystal-packing interactions with the six MKI residues are compared against those in 5UMO, by counting intersecting elements. 5UMO is used as the reference and sets the maximum number of intersecting contacts. In 5UMO, crystal contacts are formed in five of the six MKI residues (5UMO A258 has no crystal neighbor within 5 Å), and these form 16 contacts with residues in crystal neighbors:

- I254-H123, I254-T157, I254-T158, I254-C159
- N255-T157, N255-C159
- L256-L113, L256-Q117, L256-H118, L256-L119, L256-H123, L256-F127, L256-C159
- K257-Q117
- R259-S120, R259-H123

Thus, the contact intersections with other crystal structures have a maximum overlap of 5 for the MKI residue identities (column label INTERSECTION_5UMO_MKI_RESIDUE in the dataset) and 16 for all the pairs listed above (column label INTERSECTION_5UMO_MKI_XTAL_RESIDUE). See entry 5UMO compared to self. Selected columns with contact intersections are included in **Table S2**. Data represent RSCB entries collected in November 2021.

### Supplementary Tables

**Table S1. Selected dCNA matrix elements (res. 33, 52, 62, 69, 165, 166, 167) calculated between settled states of 2P-ERK2.** Listed are contact probability differences between residue pairs that are greater than |0.1|. Positive or negative values respectively signify closer or farther distances in the final state.

| **res_i** | **res_j** | **2erk_2P.MKI → 2erk_2P.L16** | **2erk_2P.pY-R65 → 2erk_2P.L16** | **2erk_2P.solv → 2erk_2P.L16** |
|---|---|---|---|---|
| A33 | Y62 | 0.193 | 0.252 | 0.177 |
| K52 | Q103 | – | -0.276 | -0.103 |
| K52 | C164 | – | 0.117 | – |
| K52 | D165 | 0.173 | – | 0.138 |
| E69 | I54 | – | 0.215 | – |
| E69 | I101 | – | 0.136 | – |
| E69 | D165 | 0.200 | – | – |
| D165 | D147 | -0.247 | -0.243 | -0.366 |
| D165 | S151 | – | -0.466 | – |
| D165 | I163 | 0.102 | 0.157 | – |
| F166 | R68 | – | -0.546 | -0.144 |
| F166 | I72 | 0.110 | – | – |
| F166 | I82 | 0.186 | 0.379 | 0.245 |
| F166 | V143 | – | – | -0.115 |
| G167 | Y34 | 0.145 | 0.477 | 0.211 |
| G167 | R68 | 0.241 | -0.426 | – |

**Table S2. Tabulated summary of Supplementary Dataset S1.** Crystallographic metadata is provided in the first five columns ["PDBID", "SpaceGroup", Chain, organism ("Org."), and resolution ("Resol. (Å)")]. The next two columns ("Count A-loop" and "Count MKI") contain the count of backbone atoms (C, O, CA, N) modeled in the asymmetric A-loop (res. L168-V186) and selected residues of the MKI (res. I254-R259); fully modeled regions with occupancies of 1.0 for all atoms yields 76 and 24 for the 19 A-loop and six MKI residues, respectively. The next column ("Intersect. MKI") contains the count of MKI residues (I254, N255, L256, K257, and R259) that have any crystal contact (as found in 5UMO). The final column ("Intersect. MKI-XTAL") contains the count of crystal contacts that intersect with those of 5UMO (maximum of 16 listed for I254, N255, L256, K257, and R259 in the description of Supplementary Dataset S1).

| PDBID | SpaceGroup | Chain | Org. | Resol. (Å) | Count A-loop | Count MKI | Intersect. MKI | Intersect. MKI-XTAL |
|---|---|---|---|---|---|---|---|---|
| **1ERK** | P 1 21 1 | A | rat | 2.3 | 76 | 24 | 5 | 15 |
| **1GOL** | P 1 21 1 | A | rat | 2.8 | 76 | 24 | 5 | 15 |
| **1PME** | P 1 21 1 | A | human | 2 | 76 | 24 | 5 | 13 |
| **1TVO** | P 1 21 1 | A | human | 2.5 | 76 | 24 | 5 | 16 |
| **1WZY** | P 1 21 1 | A | human | 2.5 | 76 | 24 | 5 | 15 |
| **2ERK** | P 41 21 2 | A | rat | 2.4 | 76 | 24 | 0 | 0 |
| **2FYS** | P 1 21 1 | A | rat | 2.5 | 76 | 24 | 5 | 2 |
| **2FYS** | P 1 21 1 | B | rat | 2.5 | 44 | 24 | 5 | 1 |
| **2GPH** | P 21 21 21 | A | rat | 1.9 | 76 | 24 | 5 | 2 |
| **2OJG** | P 1 21 1 | A | human | 2 | 76 | 24 | 5 | 16 |
| **2OJI** | P 1 21 1 | A | human | 2.6 | 76 | 24 | 5 | 15 |
| **2OJJ** | P 1 21 1 | A | human | 2.4 | 76 | 24 | 5 | 16 |
| **2Y9Q** | P 21 21 21 | A | human | 1.55 | 76 | 24 | 2 | 0 |
| **2Z7L** | P 1 21 1 | A | rat | 2.41 | 76 | 24 | 5 | 15 |
| **3C9W** | P 1 21 1 | A | rat | 2.5 | 28 | 24 | 5 | 2 |
| **3C9W** | P 1 21 1 | B | rat | 2.5 | 28 | 24 | 5 | 2 |
| **3ERK** | P 1 21 1 | A | rat | 2.1 | 76 | 24 | 5 | 14 |
| **3I5Z** | P 21 21 21 | A | human | 2.2 | 20 | 24 | 5 | 2 |
| **3I60** | P 21 21 21 | A | human | 2.5 | 24 | 24 | 5 | 2 |
| **3O71** | P 3 2 1 | A | rat | 1.95 | 56 | 24 | 5 | 0 |
| **3QYW** | P 1 21 1 | A | rat | 1.5 | 76 | 24 | 5 | 13 |
| **3QYZ** | P 1 21 1 | A | rat | 1.46 | 76 | 24 | 5 | 13 |
| **3R63** | P 1 21 1 | A | rat | 1.7 | 76 | 24 | 5 | 15 |
| **3SA0** | P 1 21 1 | A | human | 1.59 | 76 | 26 | 5 | 13 |

**Table S2 continued from previous page**

| PDBID | SpaceGroup | Chain | Org. | Resol. (Å) | Count A-loop | Count MKI | Intersect. MKI | Intersect. MKI-XTAL |
|---|---|---|---|---|---|---|---|---|
| **3TEI** | P 21 21 21 | A | human | 2.4 | 48 | 24 | 1 | 0 |
| **3W55** | P 21 21 21 | A | human | 3 | 28 | 24 | 4 | 1 |
| **3ZU7** | P 21 21 21 | A | rat | 1.97 | 76 | 24 | 1 | 0 |
| **3ZUV** | P 21 21 21 | A | rat | 2.72 | 76 | 28 | 5 | 0 |
| **3ZUV** | P 21 21 21 | C | rat | 2.72 | 76 | 24 | 5 | 0 |
| **4ERK** | P 1 21 1 | A | rat | 2.2 | 76 | 24 | 5 | 16 |
| **4FMQ** | P 21 21 21 | A | human | 2.1 | 76 | 24 | 0 | 0 |
| **4FUX** | P 1 21 1 | A | human | 2.2 | 76 | 24 | 5 | 13 |
| **4FUY** | P 1 21 1 | A | human | 2 | 76 | 25 | 5 | 13 |
| **4FV0** | P 1 21 1 | A | human | 2.1 | 76 | 24 | 5 | 13 |
| **4FV1** | P 1 21 1 | A | human | 1.99 | 77 | 24 | 5 | 13 |
| **4FV2** | P 1 21 1 | A | human | 2 | 78 | 24 | 5 | 13 |
| **4FV3** | P 1 21 1 | A | human | 2.2 | 77 | 25 | 5 | 13 |
| **4FV4** | P 21 21 21 | A | human | 2.5 | 28 | 24 | 5 | 2 |
| **4FV5** | P 21 21 21 | A | human | 2.4 | 20 | 24 | 5 | 2 |
| **4FV6** | P 21 21 21 | A | human | 2.5 | 20 | 24 | 5 | 2 |
| **4FV7** | P 1 21 1 | A | human | 1.9 | 76 | 24 | 5 | 13 |
| **4FV8** | P 21 21 21 | A | human | 2 | 24 | 24 | 5 | 2 |
| **4FV9** | P 21 21 21 | A | human | 2.11 | 20 | 24 | 5 | 2 |
| **4G6N** | P 1 21 1 | A | human | 2 | 76 | 24 | 5 | 13 |
| **4G6O** | P 1 21 1 | A | human | 2.2 | 76 | 24 | 5 | 13 |
| **4GSB** | P 1 21 1 | A | rat | 1.8 | 77 | 28 | 5 | 13 |
| **4GT3** | P 1 21 1 | A | rat | 1.68 | 84 | 29 | 5 | 13 |
| **4GVA** | P 1 21 1 | A | rat | 1.83 | 76 | 29 | 5 | 13 |
| **4H3P** | P 1 | A | human | 2.3 | 48 | 24 | 0 | 0 |
| **4H3P** | P 1 | D | human | 2.3 | 36 | 24 | 0 | 0 |
| **4H3Q** | P 21 21 21 | A | human | 2.2 | 76 | 24 | 0 | 0 |
| **4I5H** | P 31 2 1 | A | rat | 1.9 | 48 | 24 | 5 | 4 |

**Table S2 continued from previous page**

| **PDBID** | SpaceGroup | Chain | Org. | Resol. (Å) | Count A-loop | Count MKI | Intersect. MKI | Intersect. MKI-XTAL |
|---|---|---|---|---|---|---|---|---|
| **4IZ5** | P 1 21 1 | A | human | 3.19 | 76 | 24 | 5 | 0 |
| **4IZ5** | P 1 21 1 | B | human | 3.19 | 76 | 24 | 3 | 0 |
| **4IZ5** | P 1 21 1 | C | human | 3.19 | 76 | 24 | 5 | 0 |
| **4IZ5** | P 1 21 1 | D | human | 3.19 | 76 | 24 | 3 | 0 |
| **4IZ7** | P 21 21 2 | A | human | 1.8 | 52 | 24 | 5 | 0 |
| **4IZ7** | P 21 21 2 | C | human | 1.8 | 40 | 24 | 5 | 0 |
| **4IZA** | P 21 21 2 | A | human | 1.93 | 80 | 24 | 5 | 0 |
| **4IZA** | P 21 21 2 | C | human | 1.93 | 28 | 24 | 5 | 0 |
| **4N0S** | P 1 21 1 | A | human | 1.8 | 77 | 26 | 5 | 13 |
| **4N4S** | P 1 | A | rat | 2.2 | 48 | 24 | 5 | 2 |
| **4N4S** | P 1 | B | rat | 2.2 | 28 | 24 | 5 | 3 |
| **4NIF** | P 1 21 1 | B | human | 2.15 | 64 | 24 | 0 | 0 |
| **4NIF** | P 1 21 1 | E | human | 2.15 | 64 | 24 | 0 | 0 |
| **4O6E** | P 1 21 1 | A | human | 1.95 | 32 | 24 | 5 | 2 |
| **4QP1** | P 41 21 2 | A | human | 2.7 | 76 | 24 | 3 | 0 |
| **4QP1** | P 41 21 2 | B | human | 2.7 | 76 | 16 | 1 | 0 |
| **4QP2** | P 41 21 2 | A | human | 2.23 | 52 | 24 | 1 | 0 |
| **4QP2** | P 41 21 2 | B | human | 2.23 | 64 | 0 | 0 | 0 |
| **4QP3** | P 41 21 2 | A | human | 2.6 | 52 | 24 | 3 | 0 |
| **4QP3** | P 41 21 2 | B | human | 2.6 | 64 | 24 | 1 | 0 |
| **4QP4** | P 1 21 1 | A | human | 2.2 | 44 | 24 | 5 | 2 |
| **4QP4** | P 1 21 1 | B | human | 2.2 | 32 | 24 | 5 | 3 |
| **4QP6** | P 41 21 2 | A | human | 3.1 | 52 | 24 | 3 | 0 |
| **4QP6** | P 41 21 2 | B | human | 3.1 | 64 | 24 | 1 | 0 |
| **4QP7** | P 1 21 1 | A | human | 2.25 | 44 | 24 | 5 | 2 |
| **4QP7** | P 1 21 1 | B | human | 2.25 | 32 | 24 | 5 | 2 |
| **4QP8** | P 1 21 1 | A | human | 2.45 | 32 | 24 | 5 | 2 |
| **4QP8** | P 1 21 1 | B | human | 2.45 | 44 | 24 | 5 | 2 |

**Table S2 continued from previous page**

| **PDBID** | SpaceGroup | Chain | Org. | Resol. (Å) | Count A-loop | Count MKI | Intersect. MKI | Intersect. MKI-XTAL |
|---|---|---|---|---|---|---|---|---|
| **4QP9** | P 1 21 1 | A | human | 2 | 44 | 24 | 5 | 2 |
| **4QPA** | P 41 21 2 | A | human | 2.85 | 64 | 24 | 0 | 0 |
| **4QPA** | P 41 21 2 | B | human | 2.85 | 68 | 24 | 0 | 0 |
| **4QTA** | P 21 21 21 | A | human | 1.45 | 24 | 24 | 1 | 0 |
| **4QTE** | P 32 2 1 | A | human | 1.5 | 79 | 32 | 5 | 2 |
| **4QYY** | P 21 21 2 | A | rat | 1.65 | 76 | 24 | 5 | 3 |
| **4S2Z** | P 1 21 1 | A | rat | 1.48 | 76 | 24 | 5 | 14 |
| **4S30** | P 1 21 1 | A | rat | 2 | 76 | 24 | 5 | 15 |
| **4S31** | P 1 21 1 | A | rat | 1.45 | 76 | 24 | 5 | 14 |
| **4S32** | P 1 21 1 | A | rat | 1.34 | 76 | 24 | 5 | 16 |
| **4S33** | P 1 21 1 | A | rat | 1.48 | 76 | 24 | 5 | 15 |
| **4S34** | P 1 21 1 | A | rat | 2.5 | 76 | 24 | 5 | 16 |
| **4XJ0** | P 41 21 2 | A | human | 2.58 | 76 | 20 | 2 | 0 |
| **4XJ0** | P 41 21 2 | B | human | 2.58 | 76 | 12 | 2 | 0 |
| **4XNE** | P 1 21 1 | A | rat | 1.8 | 76 | 24 | 5 | 13 |
| **4XOY** | P 1 21 1 | A | rat | 2.1 | 64 | 24 | 5 | 13 |
| **4XOZ** | P 1 21 1 | A | rat | 1.95 | 76 | 24 | 5 | 13 |
| **4XP0** | P 1 21 1 | A | rat | 1.46 | 76 | 24 | 5 | 13 |
| **4XP2** | P 1 21 1 | A | rat | 1.75 | 76 | 24 | 5 | 13 |
| **4XP3** | P 1 21 1 | A | rat | 1.78 | 76 | 24 | 5 | 13 |
| **4XRJ** | P 1 21 1 | A | rat | 1.69 | 76 | 26 | 5 | 13 |
| **4XRL** | P 1 21 1 | A | rat | 2.55 | 76 | 25 | 5 | 13 |
| **4ZXT** | P 1 21 1 | A | human | 2 | 76 | 26 | 5 | 13 |
| **4ZZM** | P 1 21 1 | A | human | 1.89 | 76 | 24 | 5 | 13 |
| **4ZZN** | P 1 21 1 | A | human | 1.33 | 76 | 24 | 5 | 13 |
| **4ZZO** | P 1 21 1 | A | human | 1.63 | 76 | 24 | 5 | 13 |
| **5AX3** | P 21 21 21 | A | human | 2.98 | 64 | 24 | 0 | 0 |
| **5BUE** | P 21 21 21 | A | human | 2.4 | 20 | 24 | 5 | 2 |

**Table S2 continued from previous page**

| **PDBID** | SpaceGroup | Chain | Org. | Resol. (Å) | Count A-loop | Count MKI | Intersect. MKI | Intersect. MKI-XTAL |
|---|---|---|---|---|---|---|---|---|
| **5BUI** | P 21 21 21 | A | human | 2.12 | 40 | 24 | 5 | 2 |
| **5BUJ** | P 21 21 21 | A | human | 1.85 | 20 | 24 | 5 | 2 |
| **5BVD** | P 21 21 21 | A | human | 1.9 | 16 | 24 | 5 | 2 |
| **5BVE** | P 21 21 21 | A | human | 2 | 16 | 25 | 5 | 2 |
| **5BVF** | P 21 21 21 | A | human | 1.9 | 16 | 24 | 5 | 2 |
| **5HD4** | P 21 21 2 | A | rat | 1.45 | 76 | 24 | 5 | 3 |
| **5HD7** | P 21 21 2 | A | rat | 1.69 | 76 | 28 | 5 | 3 |
| **5K4I** | P 21 21 21 | A | human | 1.76 | 24 | 24 | 5 | 2 |
| **5KE0** | P 21 21 2 | A | rat | 1.68 | 76 | 24 | 5 | 4 |
| **5LCJ** | P 1 21 1 | A | human | 1.78 | 76 | 25 | 5 | 13 |
| **5LCK** | P 1 21 1 | A | human | 1.89 | 76 | 26 | 5 | 13 |
| **5NGU** | P 1 21 1 | A | human | 2.74 | 76 | 24 | 5 | 13 |
| **5NHF** | P 1 21 1 | A | human | 2.14 | 76 | 24 | 5 | 12 |
| **5NHH** | P 1 21 1 | A | human | 1.94 | 76 | 24 | 5 | 14 |
| **5NHJ** | P 1 21 1 | A | human | 2.12 | 76 | 24 | 5 | 14 |
| **5NHL** | P 1 21 1 | A | human | 2.07 | 76 | 24 | 5 | 15 |
| **5NHO** | P 1 21 1 | A | human | 2.24 | 76 | 24 | 5 | 15 |
| **5NHP** | P 1 21 1 | A | human | 1.99 | 76 | 24 | 5 | 15 |
| **5NHV** | P 1 21 1 | A | human | 2 | 76 | 24 | 5 | 16 |
| **5U6I** | P 21 21 2 | A | rat | 1.69 | 76 | 24 | 5 | 3 |
| ***5UMO*** | **P 1 21 1** | **A** | **rat** | **2.26** | **76** | **24** | **5** | **16** |
| **5V60** | P 21 21 21 | A | human | 2.18 | 76 | 24 | 4 | 0 |
| **5V61** | P 21 21 21 | A | human | 2.2 | 76 | 24 | 4 | 0 |
| **5V62** | P 21 21 21 | A | human | 1.9 | 76 | 24 | 5 | 0 |
| **5WP1** | P 1 21 1 | A | human | 1.4 | 80 | 25 | 5 | 13 |
| **6CPW** | P 21 21 2 | A | rat | 1.85 | 76 | 24 | 5 | 3 |
| **6D5Y** | P 21 21 21 | A | human | 2.86 | 44 | 24 | 5 | 2 |
| **6DCG** | P 21 21 2 | A | rat | 1.45 | 76 | 24 | 5 | 4 |

**Table S2 continued from previous page**

| **PDBID** | SpaceGroup | Chain | Org. | Resol. (Å) | Count A-loop | Count MKI | Intersect. MKI | Intersect. MKI-XTAL |
|---|---|---|---|---|---|---|---|---|
| **6DMG** | P 1 21 1 | A | human | 2.2 | 77 | 25 | 5 | 13 |
| **6FI3** | P 1 21 1 | A | rat | 1.52 | 76 | 24 | 5 | 13 |
| **6FI6** | P 1 21 1 | A | rat | 1.65 | 77 | 24 | 5 | 13 |
| **6FJ0** | P 1 21 1 | A | rat | 1.66 | 77 | 24 | 5 | 13 |
| **6FJB** | P 1 21 1 | A | rat | 1.85 | 76 | 24 | 5 | 13 |
| **6FJZ** | P 1 21 1 | A | rat | 1.86 | 76 | 25 | 5 | 13 |
| **6FLE** | P 1 21 1 | A | rat | 1.48 | 76 | 24 | 5 | 13 |
| **6FLV** | P 1 21 1 | A | rat | 1.91 | 76 | 25 | 5 | 13 |
| **6FMA** | P 1 21 1 | A | rat | 1.67 | 93 | 24 | 5 | 13 |
| **6FN5** | P 1 21 1 | A | rat | 1.93 | 76 | 25 | 5 | 13 |
| **6FQ7** | P 1 21 1 | A | rat | 1.6 | 92 | 24 | 5 | 13 |
| **6FR1** | P 1 21 1 | A | rat | 1.56 | 76 | 25 | 5 | 13 |
| **6FRP** | P 1 21 1 | A | rat | 1.53 | 76 | 24 | 5 | 13 |
| **6FXV** | P 1 21 1 | A | rat | 1.53 | 77 | 24 | 5 | 13 |
| **6G54** | P 32 2 1 | A | human | 2.05 | 76 | 24 | 5 | 2 |
| **6G8X** | P 1 21 1 | A | human | 1.76 | 76 | 24 | 5 | 13 |
| **6G91** | P 1 21 1 | A | human | 1.8 | 76 | 24 | 5 | 13 |
| **6G92** | P 1 21 1 | A | human | 1.99 | 76 | 24 | 5 | 13 |
| **6G93** | P 1 21 1 | A | human | 1.67 | 76 | 24 | 5 | 13 |
| **6G97** | P 1 21 1 | A | human | 1.9 | 76 | 24 | 5 | 13 |
| **6G9A** | P 1 21 1 | A | human | 1.91 | 76 | 24 | 5 | 13 |
| **6G9D** | P 1 21 1 | A | human | 1.8 | 76 | 28 | 5 | 13 |
| **6G9H** | P 1 21 1 | A | human | 1.73 | 76 | 24 | 5 | 13 |
| **6G9J** | P 1 21 1 | A | human | 1.98 | 77 | 24 | 5 | 13 |
| **6G9K** | P 1 21 1 | A | human | 1.94 | 76 | 25 | 5 | 13 |
| **6G9M** | P 1 21 1 | A | human | 1.86 | 76 | 25 | 5 | 13 |
| **6G9N** | P 1 21 1 | A | human | 1.76 | 77 | 25 | 5 | 13 |
| **6GDM** | P 1 21 1 | A | human | 1.91 | 76 | 25 | 5 | 13 |

**Table S2 continued from previous page**

| PDBID | SpaceGroup | Chain | Org. | Resol. (Å) | Count A-loop | Count MKI | Intersect. MKI | Intersect. MKI-XTAL |
|---|---|---|---|---|---|---|---|---|
| **6GDQ** | P 1 21 1 | A | human | 1.86 | 76 | 24 | 5 | 13 |
| **6GE0** | P 1 21 1 | A | human | 1.82 | 76 | 24 | 5 | 13 |
| **6GJB** | P 1 21 1 | A | human | 1.82 | 76 | 28 | 5 | 13 |
| **6GJD** | P 1 21 1 | A | human | 1.58 | 76 | 28 | 5 | 13 |
| **6NBS** | P 1 21 1 | A | rat | 1.9 | 76 | 24 | 5 | 15 |
| **6OPG** | P 21 21 21 | A | human | 2.9 | 76 | 24 | 4 | 0 |
| **6OPH** | P 21 21 21 | A | human | 2.4 | 76 | 24 | 4 | 0 |
| **6OPI** | P 21 21 21 | A | human | 3 | 32 | 24 | 3 | 1 |
| **6OPK** | P 21 21 21 | A | rat | 2.54 | 76 | 24 | 4 | 0 |
| **6OT6** | P 1 21 1 | A | rat | 1.65 | 76 | 28 | 5 | 16 |
| **6OTS** | C 1 2 1 | A | rat | 2.1 | 36 | 20 | 2 | 0 |
| **6Q7K** | P 1 21 1 | A | human | 1.84 | 76 | 24 | 5 | 13 |
| **6Q7S** | P 1 21 1 | A | human | 1.73 | 76 | 25 | 5 | 13 |
| **6Q7T** | P 1 21 1 | A | human | 1.6 | 76 | 25 | 5 | 13 |
| **6QA1** | P 1 21 1 | A | human | 1.58 | 76 | 24 | 5 | 13 |
| **6QA3** | P 1 21 1 | A | human | 1.57 | 76 | 24 | 5 | 13 |
| **6QA4** | P 1 21 1 | A | human | 1.6 | 76 | 24 | 5 | 13 |
| **6QAG** | P 1 21 1 | A | human | 2.07 | 76 | 24 | 5 | 12 |
| **6QAH** | P 1 21 1 | A | human | 1.58 | 76 | 24 | 5 | 13 |
| **6QAL** | P 1 21 1 | A | human | 1.57 | 76 | 24 | 5 | 13 |
| **6QAQ** | P 1 21 1 | A | human | 1.58 | 77 | 26 | 5 | 12 |
| **6QAW** | P 1 21 1 | A | human | 1.84 | 76 | 24 | 5 | 13 |
| **6RFP** | P 1 21 1 | A | rat | 1.74 | 76 | 24 | 5 | 16 |
| **6RQ4** | P 1 21 1 | A | human | 1.96 | 76 | 25 | 5 | 13 |
| **6SLG** | P 1 21 1 | A | human | 1.33 | 76 | 24 | 5 | 14 |
| **7AUV** | P 1 21 1 | A | human | 1.76 | 76 | 25 | 5 | 13 |
| **7NQQ** | P 1 21 1 | A | human | 1.94 | 76 | 25 | 5 | 13 |
| **7NQW** | P 1 21 1 | A | human | 1.77 | 76 | 24 | 5 | 13 |

**Table S2 continued from previous page**

| **PDBID** | SpaceGroup | Chain | Org. | Resol. (Å) | Count A-loop | Count MKI | Intersect. MKI | Intersect. MKI-XTAL |
|---|---|---|---|---|---|---|---|---|
| **7NR3** | P 1 21 1 | A | human | 1.9 | 76 | 25 | 5 | 13 |
| **7NR5** | P 1 21 1 | A | human | 1.77 | 76 | 26 | 5 | 13 |
| **7NR8** | P 1 21 1 | A | human | 1.63 | 76 | 24 | 5 | 13 |
| **7NR9** | P 1 21 1 | A | human | 1.91 | 76 | 24 | 5 | 12 |

### Supplementary Figures

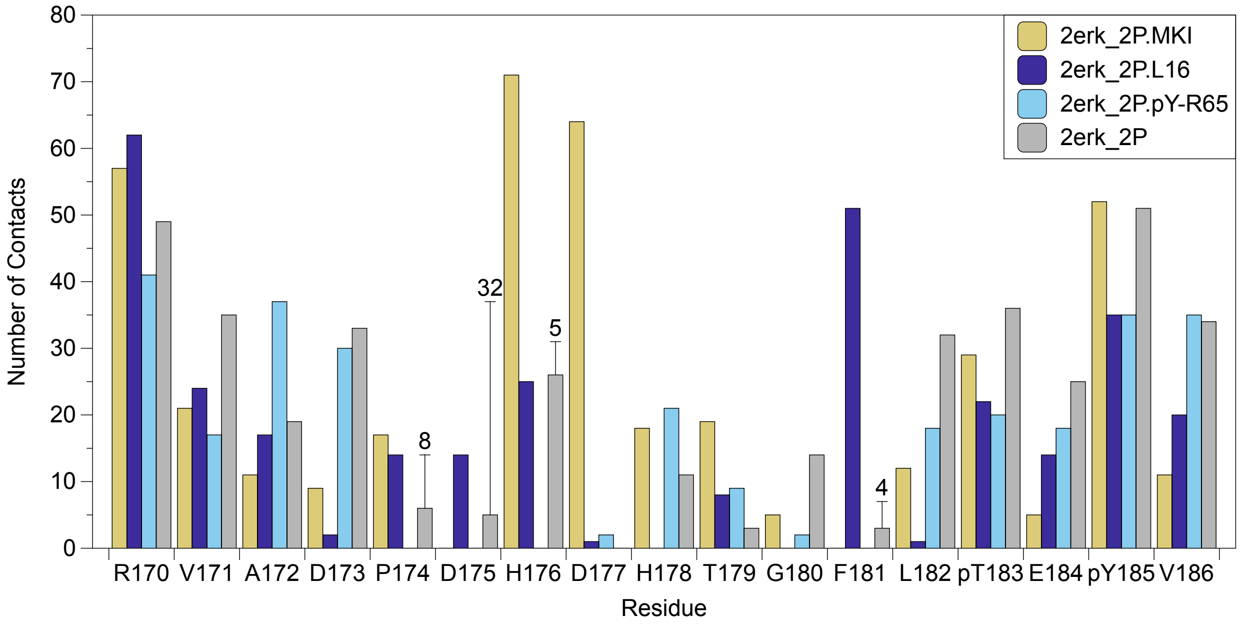


**Figure S1. Heavy atom contacts for A-loop residues in reference conformations for 2P-ERK2.** The numbers of heavy-atom contacts within 4.5 Å of residues 170-186 total to 310, 401, 285, and 382 contacts for 2erk_2P.L16, 2erk_2P.MKI, and 2erk_2P.pY-R65 reference structures and the 2ERK_2P starting state, respectively. For 2erk_2P, the contacts calculated from the energy-minimized structure are shown as grey bars, and the number of crystal lattice contacts calculated directly from the crystal coordinates of 2ERK (totaling 49) are shown as vertical lines.

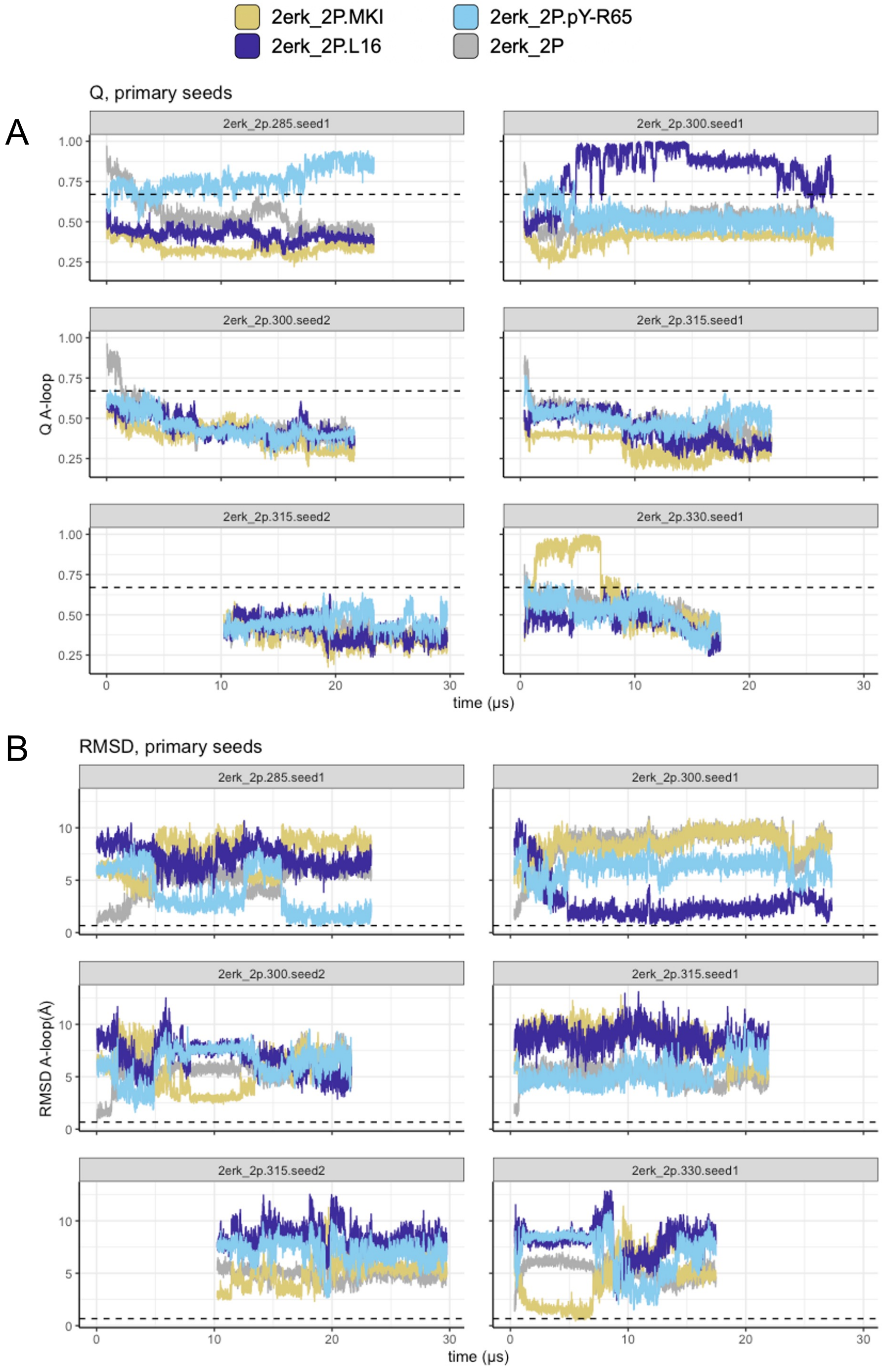


**Figure S2.** $Q_{A-loop}$ **and RMSD plots for 2P-ERK2 1° seed trajectories.** Plots of **(A)** $Q_{A-loop}$ and **(B)** RMSD for A-loop Cα atoms, calculated against the reference structures for 2erk_2P.L16, 2erk_2P.MKI, 2erk_2P.pY-R65, and the 2erk_2p starting state.

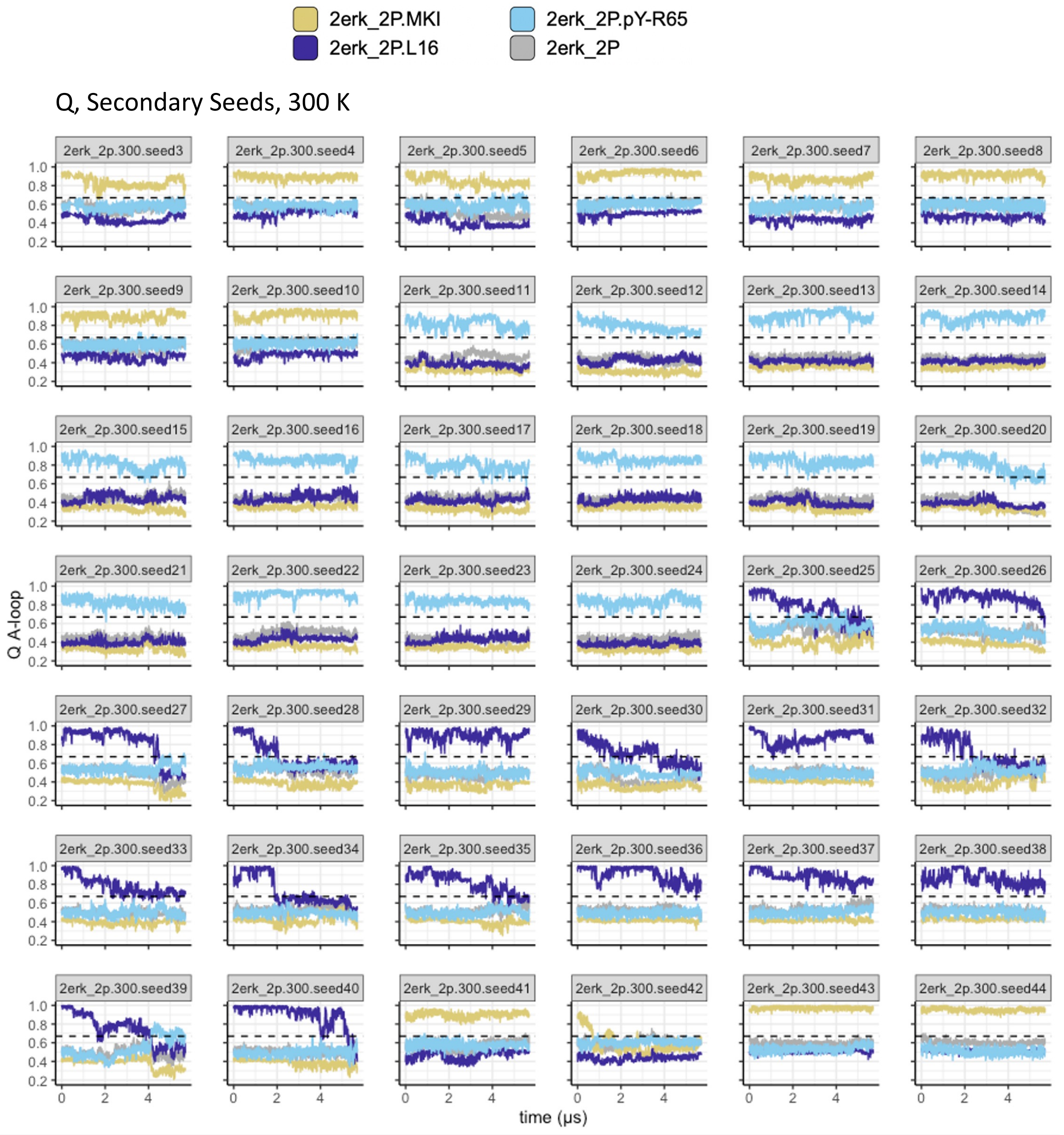


**Figure S3.** $Q_{A-loop}$ **plots for 2P-ERK2 2° seed trajectories.** Plots of $Q_{A-loop}$ calculated against the reference structures for 2erk_2P.L16, 2erk_2P.MKI, 2erk_2P.pY-R65, and the 2erk_2p starting state.

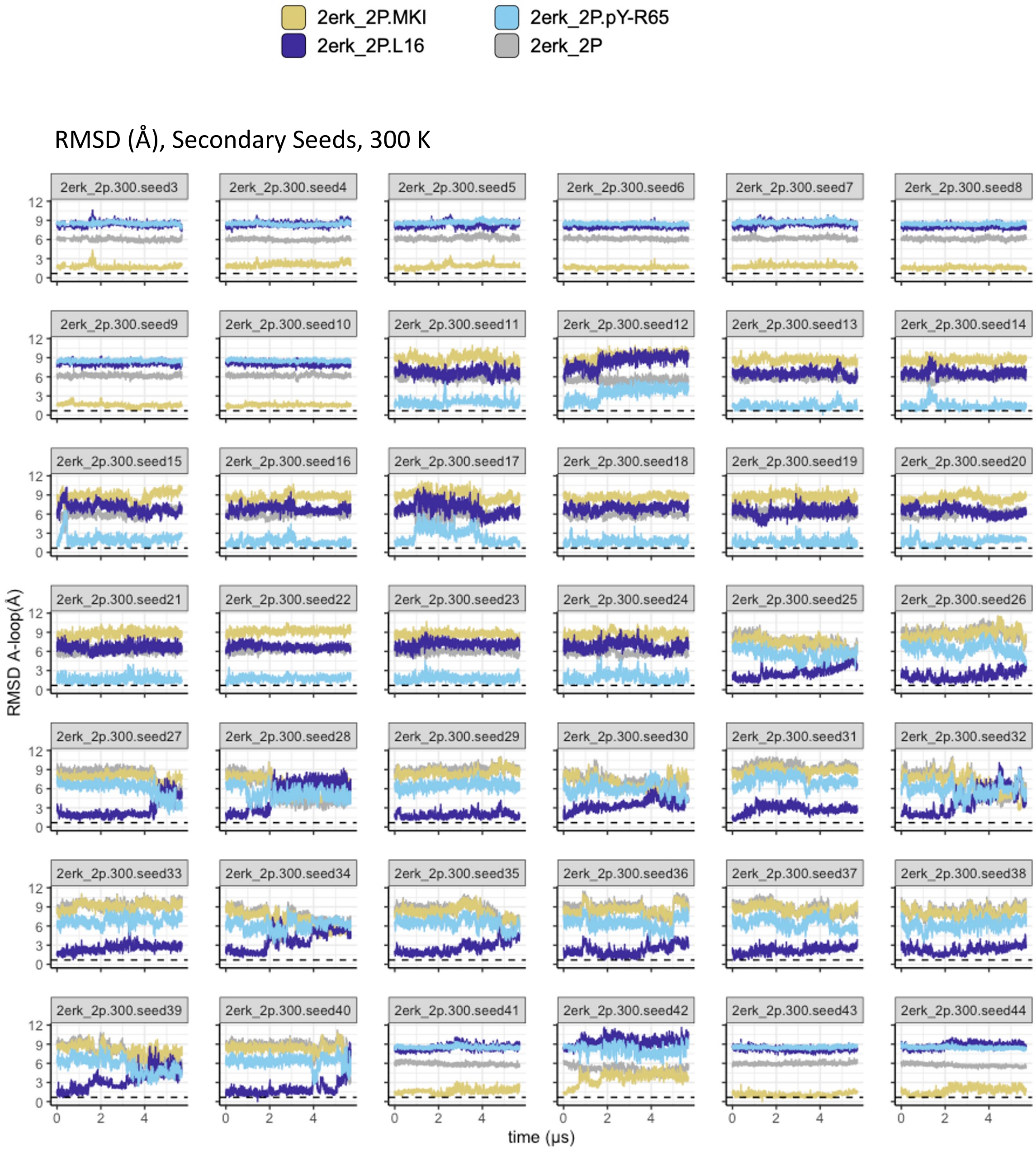


**Figure S4. RMSD plots for 2P-ERK2 2° seed trajectories.** RMSD plots for A-loop C$\alpha$ atoms calculated against the reference structures for 2erk_2P.L16, 2erk_2P.MKI, 2erk_2P.pY-R65, and the 2erk_2p starting state.

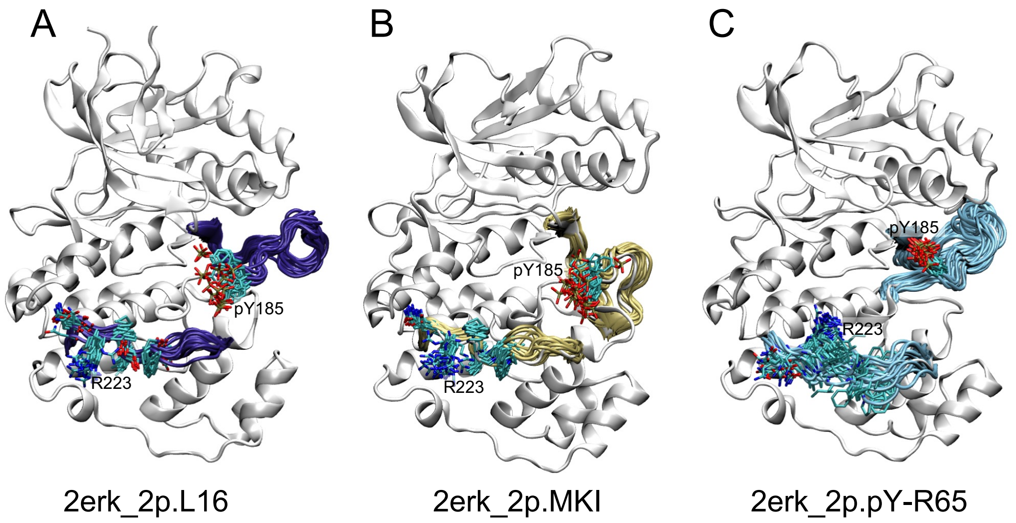


**Figure S5. Overlay plots for settled states of 2P-ERK2.** Overlays display conformational shifts of the A-loop and the loop connecting helices $\alpha$F and $\alpha$G, between **(A)** 2erk_2P.L16, **(B)** 2erk_2P.MKI, and **(C)** 2erk_2P.pY-R65. Each overlay contains 20 frames separated by 250 ns.

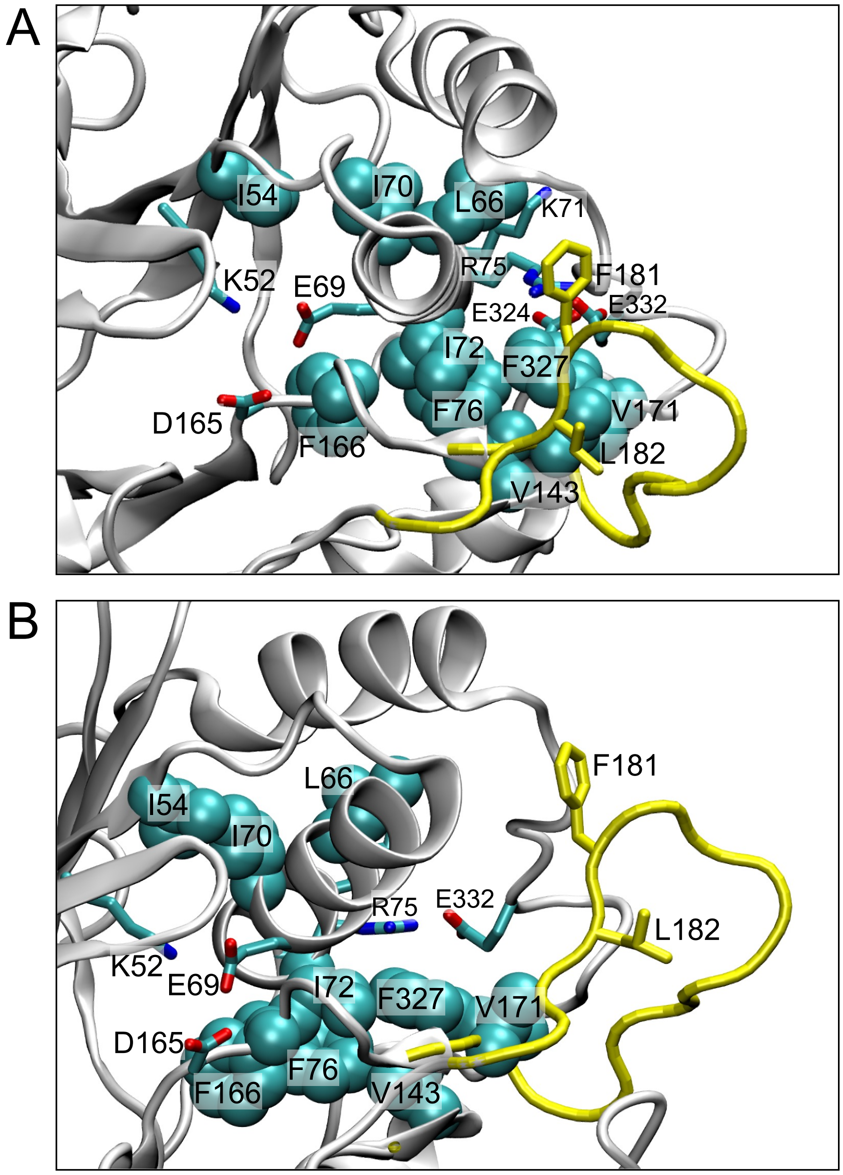


**Figure S6. Side-chain connectivity between L16 and the active site. (A, B)** Two orientations of the 2erk_2P.L16 reference structure, showing hydrophobic packing interactions between F327 in L16 and residues I72, R75 and F76 in helix $\alpha$C. Residues in helix $\alpha$C in turn form second sphere interactions with residues nearby catalytic residues in $\beta$3 (I54, nearby K52), $\beta$8-$\beta$9 (F166, nearby D165), and $\beta$6 (V143, nearby D147).

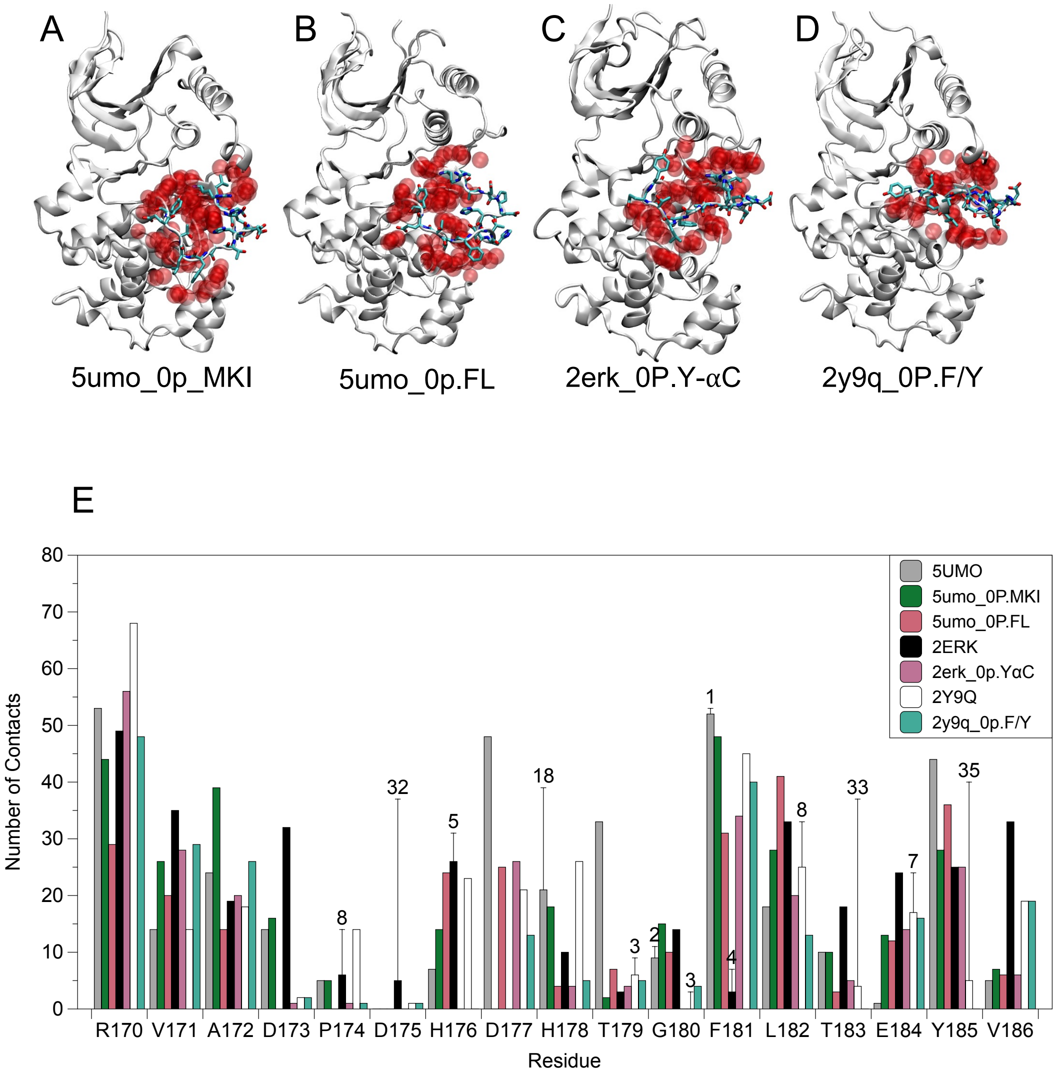


**Figure S7. Reference conformations and heavy atom contacts for settled states of 0P-ERK2.** Reference structures for **(A)** 5umo_0P.MKI, **(B)** 5umo_0P.FL, **(C)** 2erk_0P.Y-$\alpha$C, and **(D)** 2y9q_0P.F/Y. Red spheres show heavy atom contacts within 4.5 Å of A-loop residues (licorice sticks). **(E)** Numbers of heavy atom contacts for residues 170-186 in reference and X-ray structures. Heavy atom contacts total 313, 262, 244, and 222 for reference states 5umo_0P.MKI, 5umo_0P.FL, 2erk_0P.Y-$\alpha$C, and 2y9q_0P.F/Y, respectively. Also shown are A-loop contacts for energy-minimized X-ray structures, totaling 358, 308, and 335 for 5UMO, 2Y9Q, and 2ERK, respectively. Generally, the reference structures extracted from simulations showed fewer heavy atom contacts than their corresponding starting states. The numbers of crystal lattice contacts with A-loop residues, calculated using the crystal coordinates from X-ray structures, are shown as vertical lines.

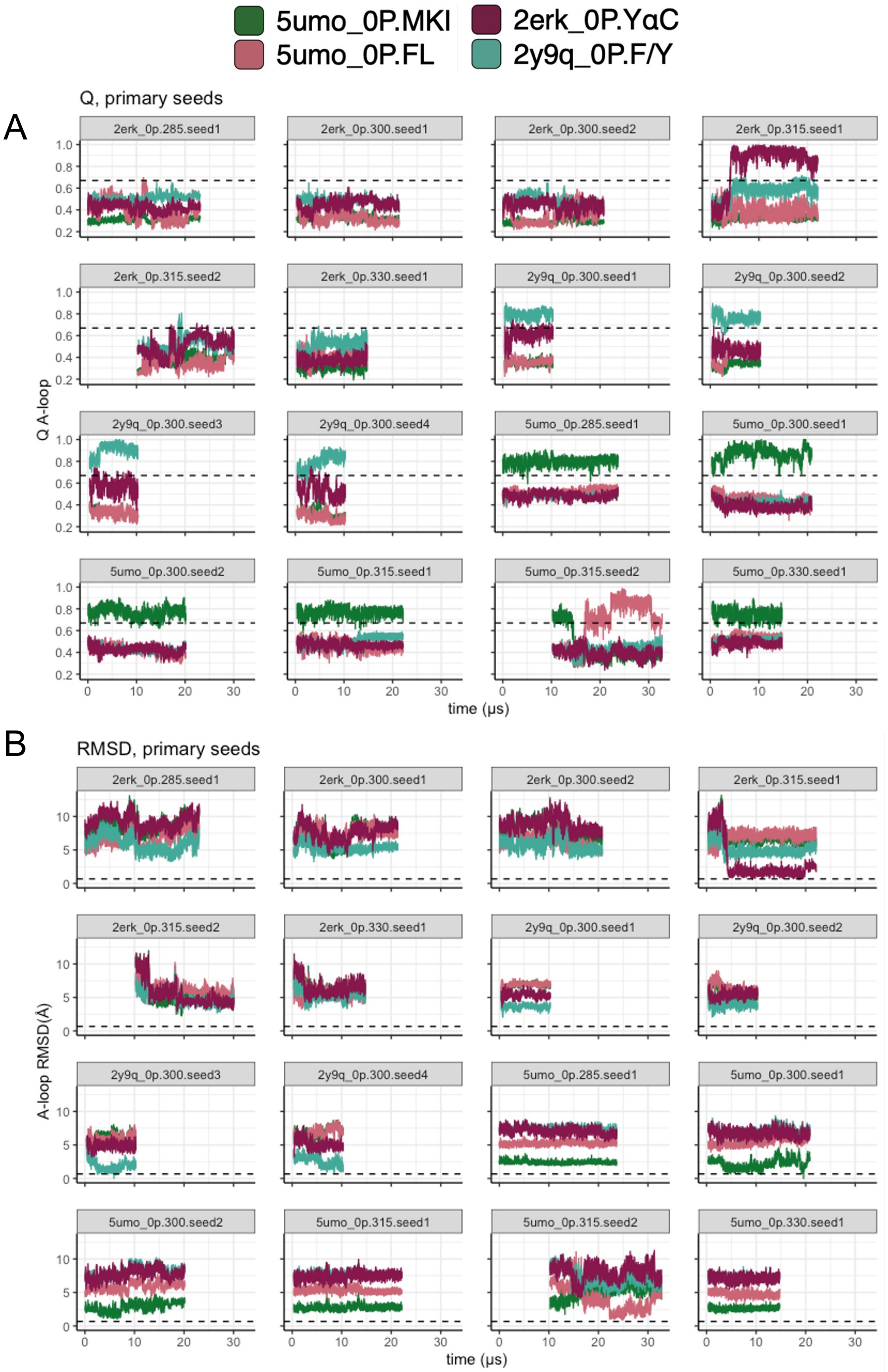


**Figure S8.** $Q_{A-loop}$ **and RMSD plots for 1° seed trajectories of 0P-ERK2.** Plots of **(A)** $Q_{A-loop}$ and **(B)** RMSD for A-loop C$\alpha$ atoms, calculated against the reference structures for 5umo_0P.MKI, 5umo_0P.FL, 2erk_0P.Y-$\alpha$C, and 2y9q_0P.F/Y.

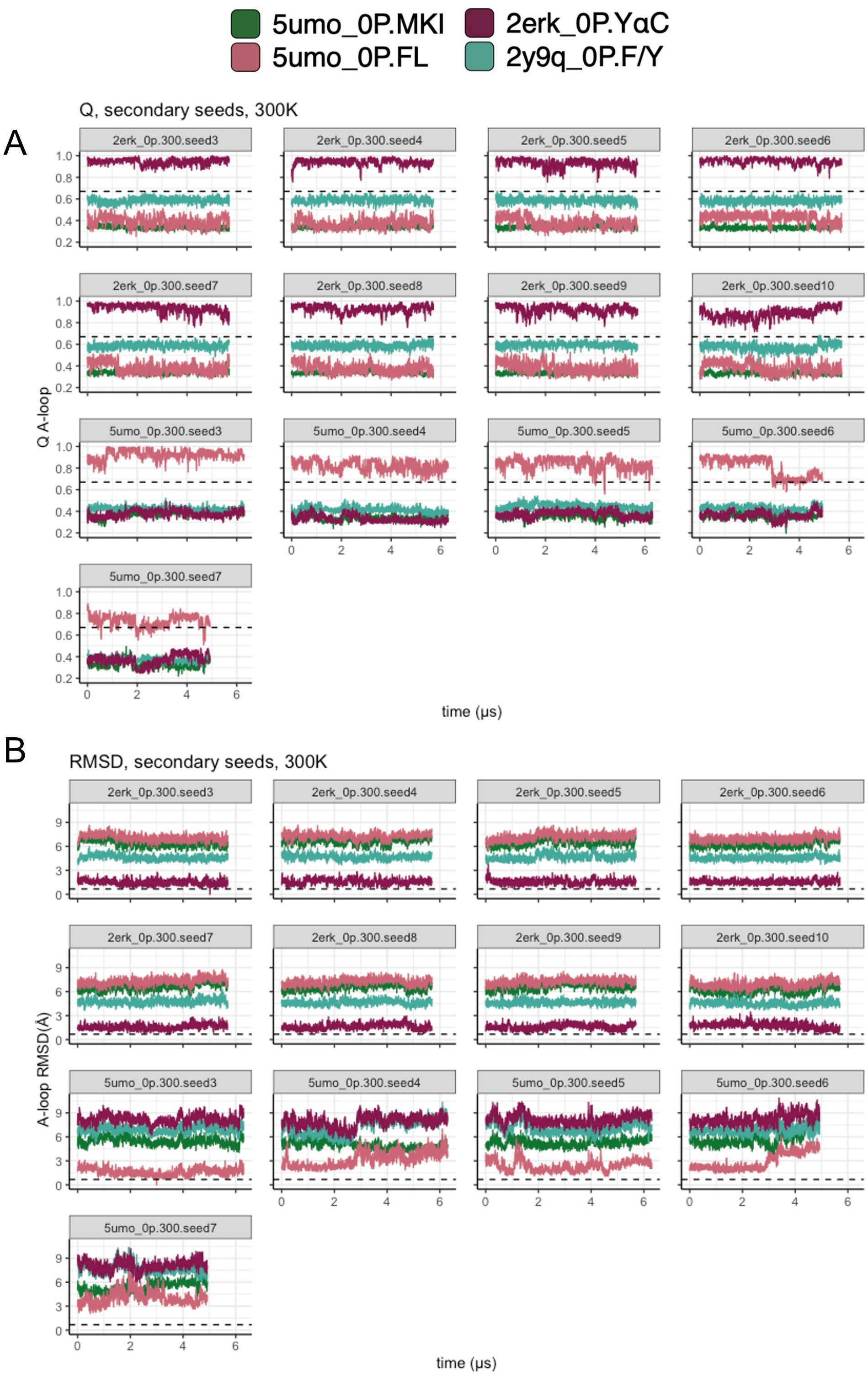


**Figure S9.** $Q_{A-loop}$ **and RMSD plots for 2° seed trajectories of 0P-ERK2.** Plots of **(A)** $Q_{A-loop}$ and **(B)** RMSD for A-loop C$\alpha$ atoms, calculated against the reference structures for 5umo_0P.MKI, 5umo_0P.FL, 2erk_0P.Y-$\alpha$C, and 2y9q_0P.F/Y.

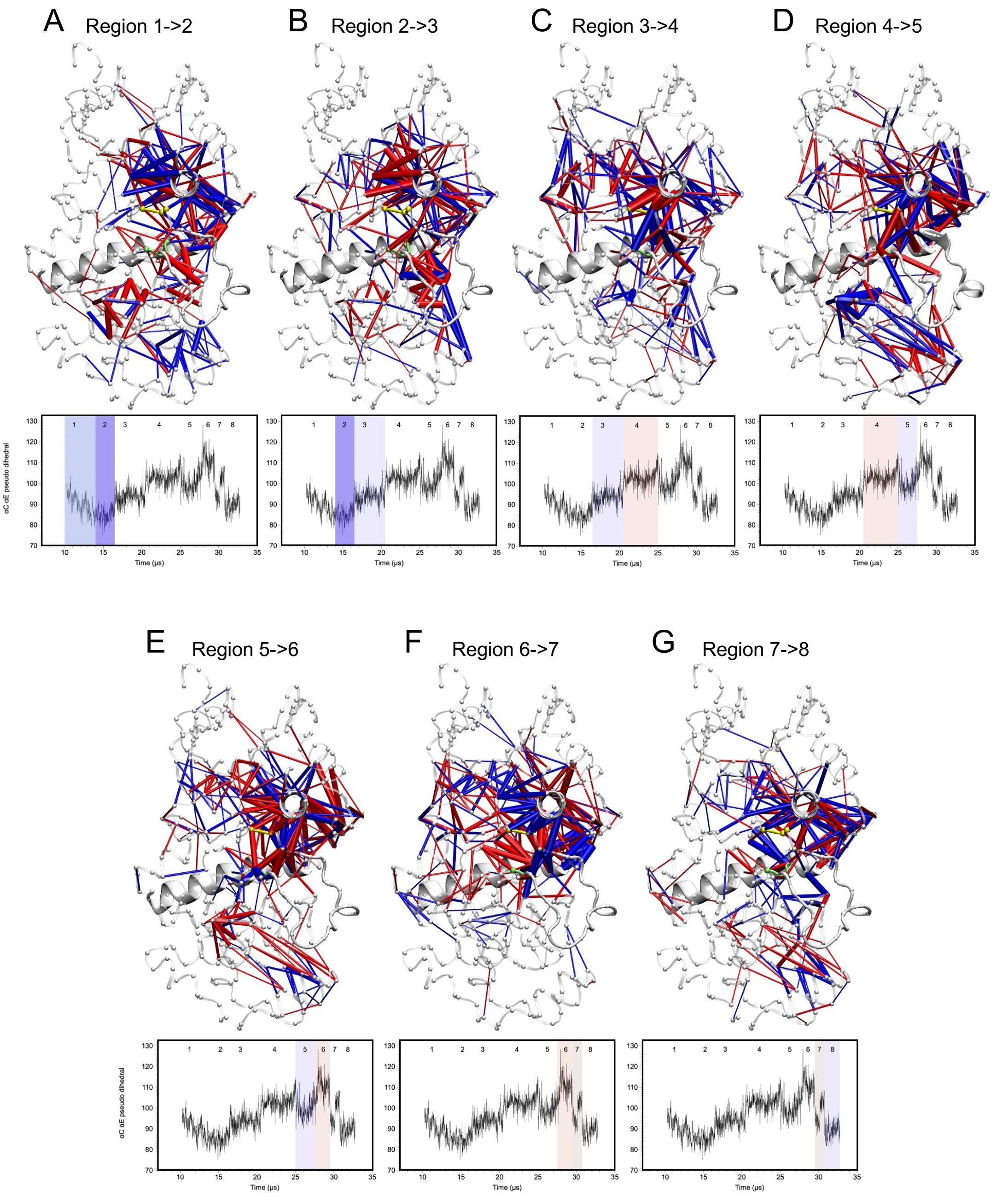


**Figure S10. Temporal dCNA for the 5umo_0P.FL 1° trajectory.** dCNA was applied to consecutive segments of the trajectory for 5umo_0p seed 2 at 315 K, which was split into eight regions based on the pseudo-dihedral angle between helices $\alpha$C and $\alpha$E. The temporal analysis revealed large fluctuations in conformation across this trajectory, leading up to and including the 5umo_0P.FL settled state.

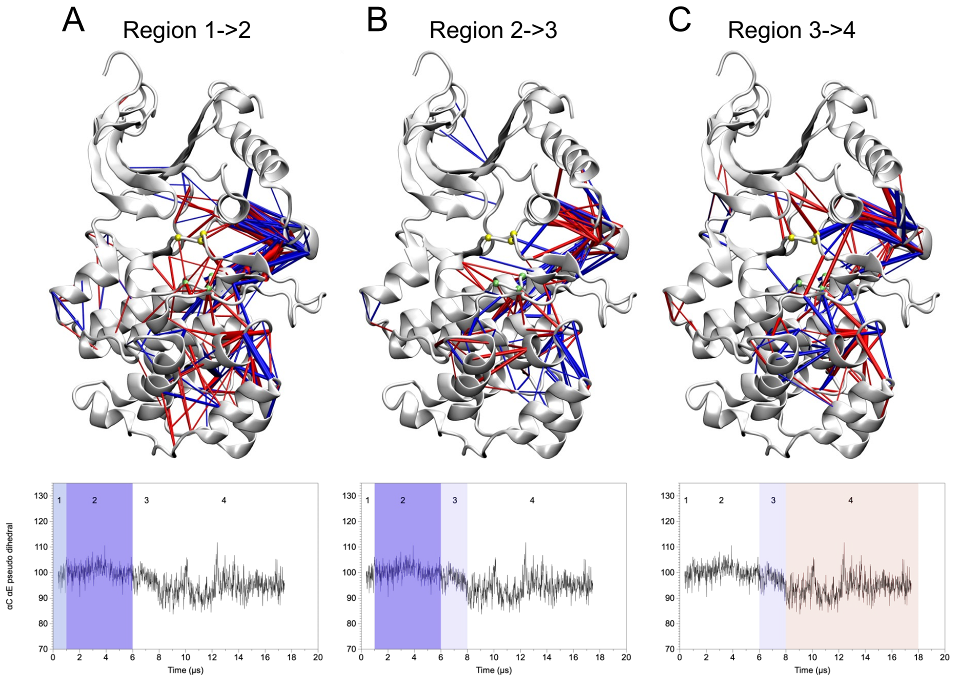


**Figure S11. Temporal dCNA for the 2erk_2P.MKI 1° trajectory.**. Time-dependent dCNA was applied to consecutive segments of the trajectory for 2erk_2p seed 1 at 330 K, which was split into four regions based on $Q_{A-loop}$ value for the 2erk_2P.MKI reference conformation. The pseudo-dihedral angle between helices $\alpha$C and $\alpha$E is also plotted, highlighting the four regions. The temporal analysis revealed few conformational fluctuations across this trajectory, corresponding to the 2erk_2P.MKI settled state.

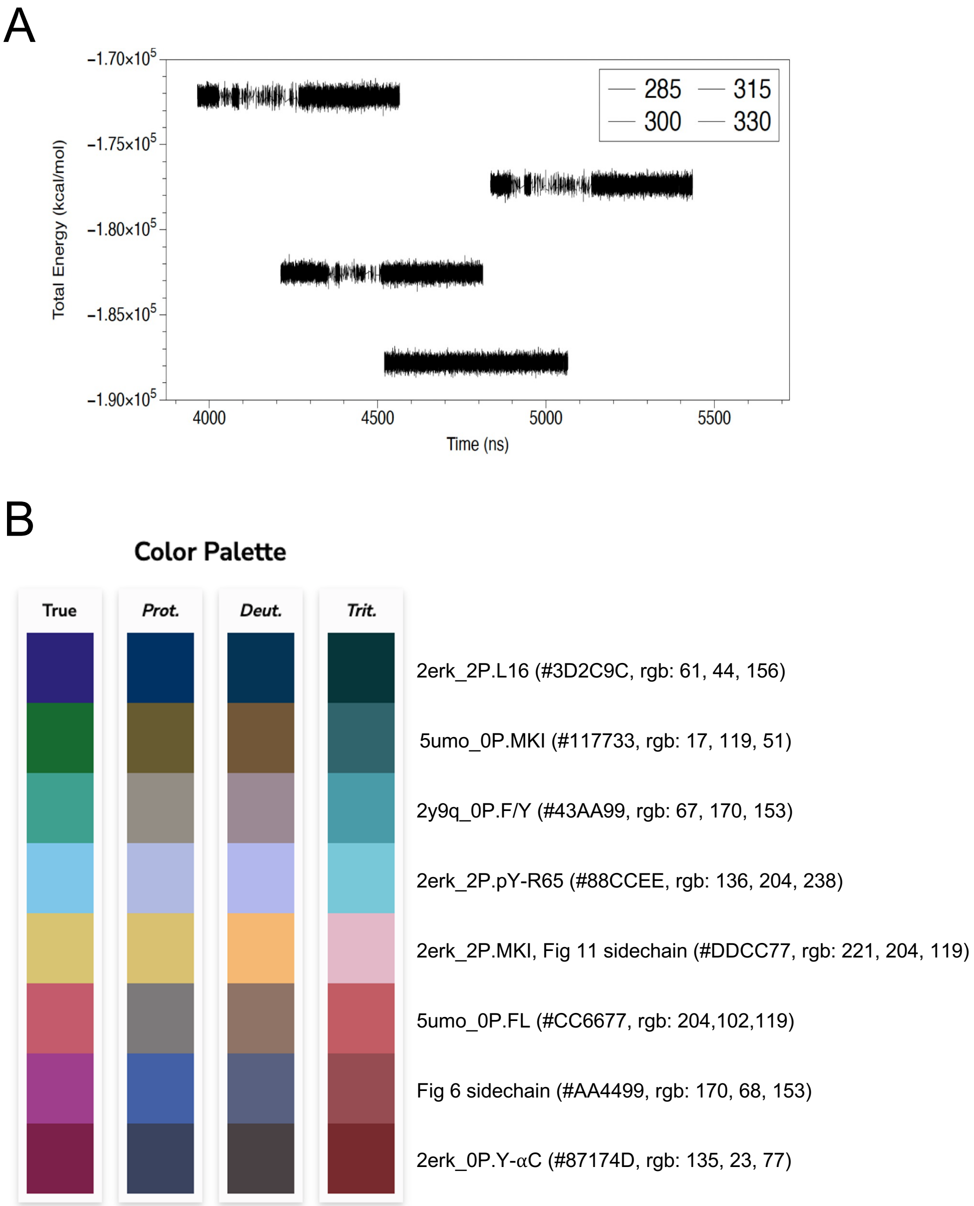


**Figure S12. Rare filesystem imposed errors and color palette used in figures. (A)** Total energy plotted *vs* time for trajectory segments with IO problems. Generally, each trajectory was run for 300 ns at a time. As a result there were around 2,450 trajectories. For a handful of runs of the 1° seeds, there were IO issues with writing the trajectories to the disk. CPPTRAJ was used to discard problematic frames in these trajectories using the "check: 1-353 skipbadframes" arguments during down-sampling and alignment. The runs shown correspond to 2erk_2p simulations performed on March 28, 2021, when the filesystem of the working directory accidentally ran out of space. The lines from highest to lowest total energy correspond to runs at 330 K, 315 K, 300 K and 285 K. For each temperature, the energy distribution is consistent after restart. The 2° seeds showed no IO issues. **(B)** Color palette used throughout for states and sidechains in specified main text figures.

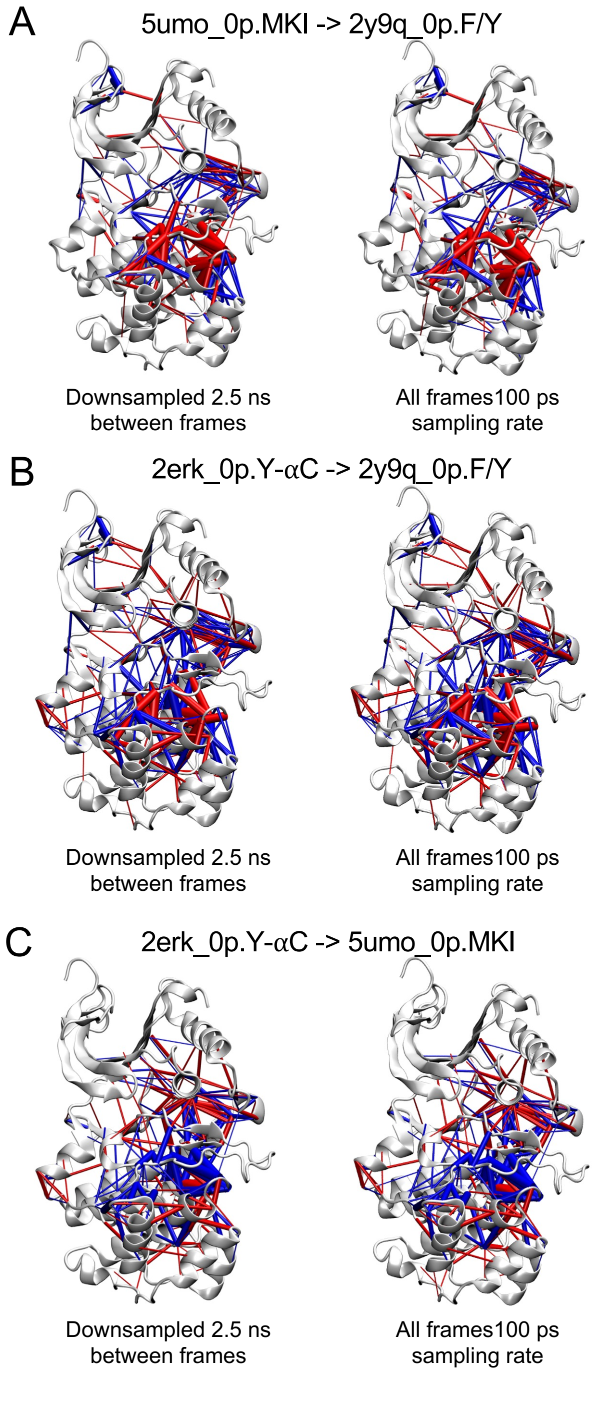


**Figure S13. Comparison of dCNA bar analyses calculated with different numbers of frames.** dCNA plots comparing **(A)** 5umo_0P.MKI to 2y9q_0P.F/Y, **(B)** 2erk_0P.Y-$\alpha$C to 2y9q_0P.F/Y, and **(C)** 2y9q_0P.F/Y to 5umo_0P.MKI were calculated and plotted after **(left panels)** downsampling (a frame every 2.5 ns), or **(right panels)** using all available frames (a frame every 100 ps). The results show that the two methods for calculating dCNA agree quantitatively.
